# scConsensus: combining supervised and unsupervised clustering for cell type identification in single-cell RNA sequencing data

**DOI:** 10.1101/2020.04.22.056473

**Authors:** Bobby Ranjan, Florian Schmidt, Wenjie Sun, Jinyu Park, Mohammad Amin Honardoost, Joanna Tan, Nirmala Arul Rayan, Shyam Prabhakar

## Abstract

Clustering is a crucial step in the analysis of single-cell data. Clusters identified using unsupervised clustering are typically annotated to cell types based on differentially expressed genes. In contrast, supervised methods use a reference panel of labelled transcriptomes to guide both clustering and cell type identification. Supervised and unsupervised clustering strategies have their distinct advantages and limitations. Therefore, they can lead to different but often complementary clustering results. Hence, a consensus approach leveraging the merits of both clustering paradigms could result in a more accurate clustering and a more precise cell type annotation. We present scConsensus, an R framework for generating a consensus clustering by (i) integrating the results from both unsupervised and supervised approaches and (ii) refining the consensus clusters using differentially expressed (DE) genes. The value of our approach is demonstrated on several existing single-cell RNA sequencing datasets, including data from sorted PBMC sub-populations. scConsensus is freely available on GitHub at https://github.com/prabhakarlab/scConsensus.

## Introduction

Since the first single cell experiment was published in 2009 (1), single cell RNA sequencing (scRNA-seq) has become the quasi-standard for transcriptomic profiling of heterogeneous data sets. In contrast to bulk RNA-sequencing, scRNA-seq is able to elucidate transcriptomic heterogeneity at an unmatched resolution and thus allows downstream analyses to be performed in a cell-type-specific manner, easily. This has been proven to be especially important for instance in case-control studies or in studying tumor heterogeneity (2). Nowadays, due to advances in experimental technologies, more than 1 million single cell transcriptomes can be profiled with high-throughput microfluidic systems. Scalable and robust computational frameworks are required to analyse such highly complex single cell data sets.

The clustering of single cells for annotation of cell types is a major step in this analysis. There are two methodologies that are commonly applied to cluster and annotate cell types: (i) unsupervised clustering followed by cluster annotation using marker genes (3) and (ii) supervised approaches that use reference data sets to either cluster cells (4) or to classify cells into cell types (5).

A wide variety of methods exist to conduct unsupervised clustering, with each method using different distance metrics, feature sets and model assumptions. The graph-based clustering method Seurat (6) and its Python counterpart Scanpy (7) are the most prevalent ones. In addition, numerous methods based on hierarchical (8), density-based (9) and k-means clustering (10) are commonly used in the field. Kiselev et al. provide an extensive overview on unsupervised clustering approaches and discuss different methodologies in detail. Importantly, they conclude that there is currently no method available that can robustly be applied to any kind of scRNA-seq data set, as method performance can be influenced by the size of data sets, the number and the nature of sequenced cell types as well as by technical aspects, such as dropouts, sample quality and batch effects.

Unsupervised clustering methods have been especially useful for the discovery of novel cell types. However, the marker-based annotation is a burden for researchers as it is a time-consuming and labour-intensive task. Also, manual, marker-based annotation can be prone to noise and dropout effects. Furthermore, different research groups tend to use different sets of marker genes to annotate clusters, rendering results to be less comparable across different laboratories.

To overcome these limitations, supervised cell type assignment and clustering approaches were proposed. The major advantages of supervised clustering over unsupervised clustering are its robustness to batch effects and its reproducibility. This has been shown to be beneficial for the integrative analysis of different data sets (4). A comprehensive review and benchmarking of 22 methods for supervised cell type classification is provided by (5). While they found that several methods achieve high accuracy in cell type identification, they also point out certain caveats: several sub-populations of CD4+ and CD8+ T cells could not be accurately identified in their experiments. Abdelaal et al. (5) traced this back to inappropriate and/or missing marker genes for these cell types in the reference data sets used by some of the methods tested. This exposes a vulnerability of supervised clustering and classification methods–the reference data sets impose a constraint on the cell types that can be detected by the method. Aside from this strong dependence on reference data, another general observation made was that the accuracy of cell type assignments decreases with an increasing number of cells and an increased pairwise similarity between them. Furthermore, clustering methods that do not allow for cells to be annotated as *Unkown*, in case they do not match any of the reference cell types, are more prone to making erroneous predictions.

In summary, despite the obvious importance of cell type identification in scRNA-seq data analysis, the single-cell community has yet to converge on one cell typing methodology (3). Due to the diverse merits and demerits of the numerous clustering approaches, this is unlikely to happen in the near future. However, as both unsupervised and supervised approaches have their distinct advantages, it is desirable to leverage the best of both to improve the clustering of single-cell data. As exemplified in Supplementary Figure (Sup. Fig.) S1 using FACS-sorted Peripheral Blood Mononuclear Cells (PBMC) scRNA-seq data from (11), both supervised and unsupervised approaches deliver unique insights into the cell type composition of the data set. Specifically, the supervised RCA (4) is able to detect different progenitor sub-types, whereas Seurat is better able to determine T-cell sub-types. Therefore, a more informative annotation could be achieved by combining the two clustering results.

## Approach

Inspired by the consensus approach used in the unsupervised clustering method SC3, which resulted in improved clustering results for small data sets compared to graph-based approaches (3, 10), we propose scConsensus, a computational framework in **R** to obtain a consensus set of clusters based on unsupervised and supervised clustering results.

Firstly, a consensus clustering is derived from the results of unsupervised and supervised methods. This consensus clustering represents cell groupings derived from both clustering results, thus incorporating information from both inputs. Details on how this consensus clustering is generated are provided in Materials and Methods section.

Secondly, the resulting consensus clusters are refined by reclustering the cells using cluster-specific differentially expressed (DE) genes (Fig. 1) as features. Each initial consensus cluster is compared in a pair-wise manner with every other cluster to maximise inter-cluster distance with respect to strong marker genes. Thereby, the separation of distinct cell types will improve, whereas clusters representing identical cell types not exhibiting distinct markers, will be merged together.

**Fig. 1.**
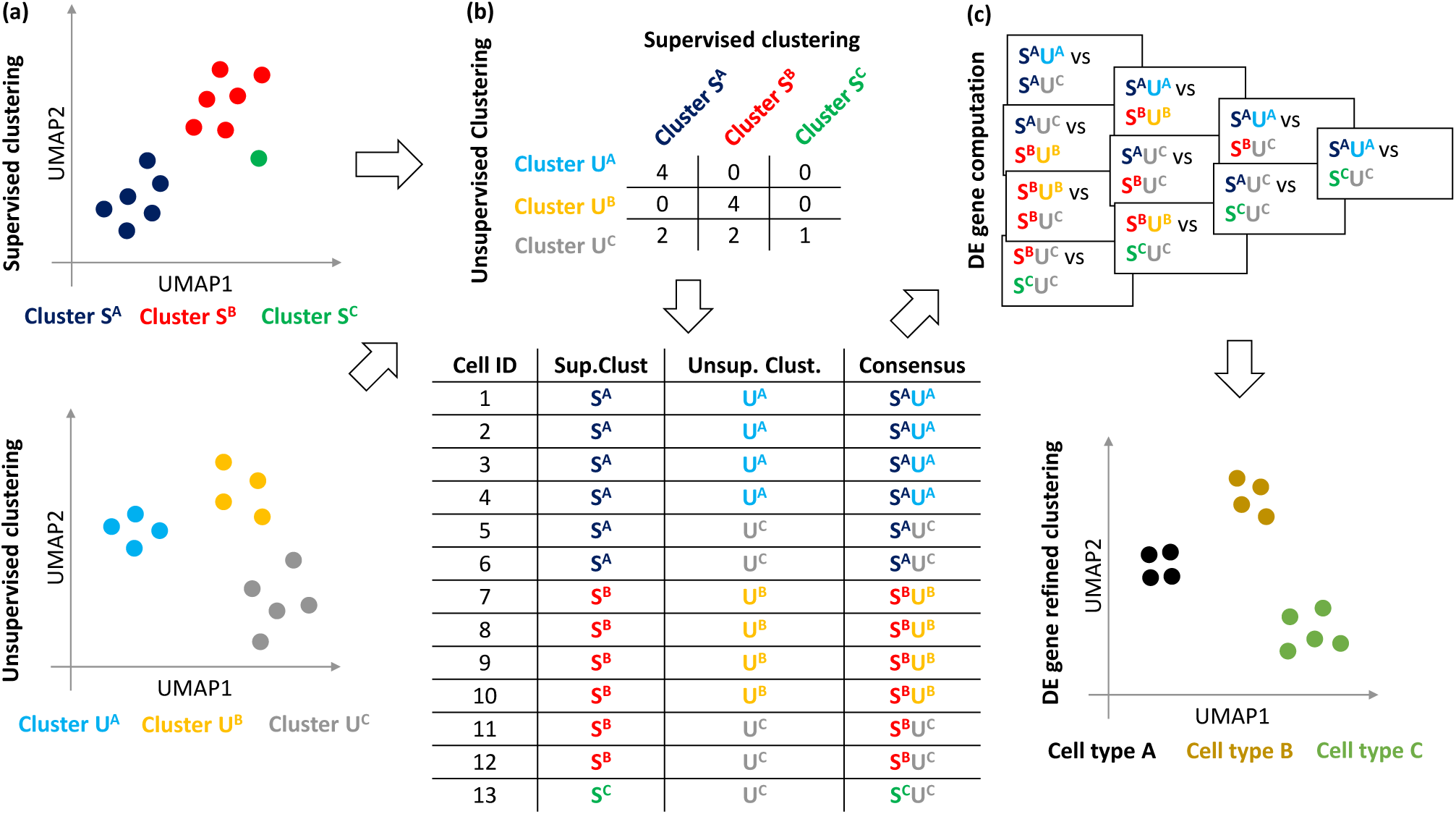
(a) The scConsensus workflow considers two independent cell cluster annotations obtained from any pair of supervised and unsupervised clustering methods. (b) A contingency table is generated to elucidate the overlap of the annotations on the single cell level. A consensus labeling is generated using either an automated method or manual curation by the user. (c) DE genes are computed between all pairs of consensus clusters. Those DE genes are used to re-cluster the data. The refined clusters thus obtained can be annotated with cell type labels.

Here, we illustrate the applicability of the scConsensus workflow by integrating cluster results from the widely used Seurat package (6) with the reference-based RCA clustering method (4).

## Materials and Methods

### Data

In total, we used five 10X CITE-Seq scRNA-seq data sets. Two data sets of 7817 Cord Blood Mononuclear Cells and 7583 PBMC cells respectively from (12) and three from 10X Genomics containing 8242 Mucosa-Associated Lymphoid cells, 7750 and 7627 PBMCs, respectively. Additionally, we downloaded FACS-sorted PBMC scRNA-seq data generated by (11) for CD14+ Monocytes, CD19+ B Cells, CD34+ Cells, CD4+ Helper T Cells, CD4+/CD25+ Regulatory T Cells, CD4+/CD45RA+/CD25-Naive T cells, CD4+/CD45RO+ Memory T Cells CD56+ Natural Killer Cells, CD8+ Cytotoxic T cells and CD8+/CD45RA+ Naive T Cells from the 10X website. Further details and download links are provided in Sup. Table S1. Table 1 provides acronyms used in the remainder of the paper. Details on processing of the FACS sorted PBMC data are provided in Supplementary *Note 3*.

**Table 1.**
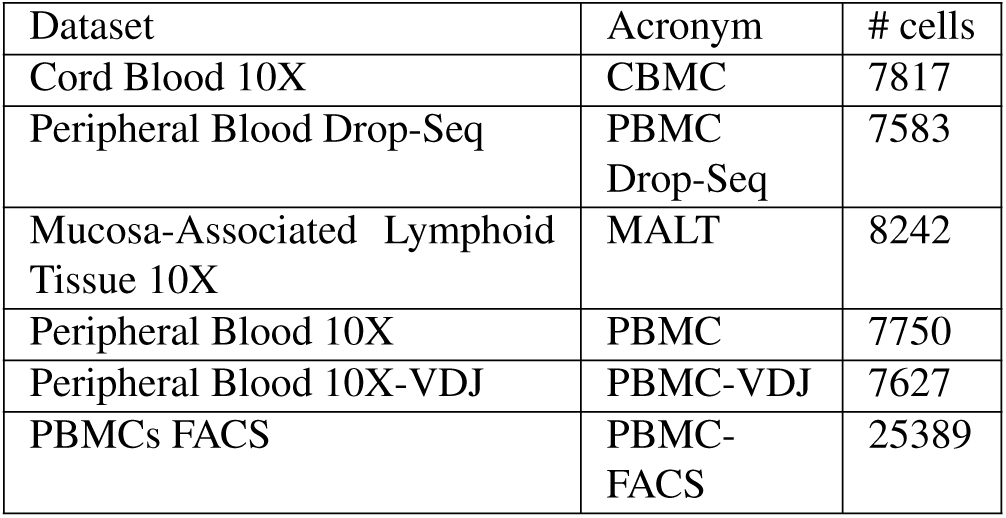
Overview on the number of cells contained in each considered scRNA-seq data set as well as on the acronyms used throughout this article.

| Dataset | Acronym | # cells |
| --- | --- | --- |
| Cord Blood 10X | CBMC | 7817 |
| Peripheral Blood Drop-Seq | PBMC<br>Drop-Seq | 7583 |
| Mucosa-Associated Lymphoid Tissue 10X | MALT | 8242 |
| Peripheral Blood 10X | PBMC | 7750 |
| Peripheral Blood 10X-VDJ | PBMC-VDJ | 7627 |
| PBMCs FACS | PBMC-FACS | 25389 |

### Data pre-processing and initial clustering

We used RCA (version 1.0) for supervised and Seurat (version 3.1.0) for unsupervised clustering (Fig.1a). As the reference panel included in RCA contains only major cell types, we generated an immune-specific reference panel containing 29 immune cell types based on sorted bulk RNA-seq data from (13). Details on the generation of this reference panel are provided in Supplementary *Note 1*.

All data pre-processing was conducted using the Seurat **R**-package. After filtering cells using a lower and upper bound for the *Number of Detected Genes (NODG)* and an upper bound for *mitochondrial rate*, we filtered out genes that are not expressed in at least 100 cells. Data set specific QC metrics are provided in Sup. Table S2. Note that we did not apply a threshold on the *Number of Unique Molecular Identifiers*. **R**-code is available in Supplementary *Note 2*.

### Workflow of scConsensus

scConsensus takes the supervised and unsupervised clustering results as input and performs the following two major steps:

1. Generation of consensus annotation using a contingency table consolidating the results from both clustering inputs,
2. Refinement of the consensus cluster labels by reclustering cells using DE genes.

The entire pipeline is visualized in Fig. 1.

#### Generating a consensus clustering

First, we use the *table* function in **R** to construct a contingency table (Fig.1 b). Each value in the contingency table refers to the extent of overlap between the clusters, measured in terms of number of cells. scConsensus provides an automated method to obtain a consensus set of cluster labels 𝒞. Starting with the clustering that has a larger number of clusters, referred to as ℒ, scConsensus determines whether there are any possible sub-clusters that are missed by ℒ. To do so, we determine for each cluster *l* ∈ ℒ the percentage of overlap for the clustering with fewer clusters (ℱ) in terms of cell numbers: |*l* ∩*f*|. By default, we consider any cluster *f* that has an overlap ≥10% with cluster *l* as a sub-cluster of cluster *l*, and then assign a new label to the overlapping cells as a combination of *l* and *f*. For cells in a cluster *l* ∈ ℒ with an overlap < 10% to any cluster *f*∈ ℱ, the original label will be retained. We note that the overlap threshold can be changed by the user. For instance by setting it to 0, each cell will obtain a label based on both considered clustering results ℱand ℒ. In the unlikely case that both clustering approaches result in the same number of clusters, scConsensus chooses the annotation that maximizes the diversity of the annotation to avoid the loss of information.

In addition to the automated consensus generation and for refinement of the latter, scConsensus provides the user with means to perform a manual cluster consolidation. This approach is especially well-suited for expert users who have a good understanding of cell types that are expected to occur in the analysed data sets.

#### Refinement by re-clustering cells on DE genes

Once the consensus clustering 𝒞 has been obtained, we determine the top 30 DE genes, ranked by the absolute value of the fold-change, between every pair of clusters in 𝒞 and use the union set of these DE genes to re-cluster the cells (Fig.1c). Note that the number of DE genes is a user parameter and can be changed. Empirically, we found that the results were relatively insensitive to this parameter (Supplementary Figure S9), and therefore it was set at a default value of 30 throughout. Typically, for UMI data, we use the edgeR (14) exactTest to determine the statistical significance of differential expression and couple that with a fold-change threshold (absolute fold-change ≥2) to select differentially expressed genes. Upon DE gene selection, Principal Component Analysis (PCA) (15) is performed to reduce the dimensionality of the data using the DE genes as features. The number of principal components (PCs) to be used can be selected using an elbow plot. For the datasets used here, we found 15 PCs to be a conservative estimate that consistently explains majority of the variance in the data (Supplementary Figure S10). We then construct a cell-cell distance matrix in PC space to cluster cells using Ward’s agglomerative hierarchical clustering approach (16).

### Clustering of antibody tags to derive a ground truth for CITE-Seq data

We used antibody-derived tags (ADTs) in the CITE-Seq data for cell type identification by clustering cells using Seurat. The raw antibody data was normalized using the Centered Log Ratio (CLR) (17) transformation method, and the normalized data was centered and scaled to mean zero and unit variance. Dimension reduction was performed using PCA. The cell clusters were determined using Seurat’s default graph-based clustering. More details, along with the source code used to cluster the data, are available in Supplementary *Note 2*.

Since these cluster labels were derived solely using ADTs, they provide an unbiased ground truth to benchmark the performance of scConsensus on scRNA-seq data. For each antibody-derived cluster, we identified the top 30 DE genes (in scRNA-seq data) that are positively up-regulated in each ADT cluster when compared to all other cells using the Seurat FindAllMarkers function. The union set of these DE genes was used for dimensionality reduction using PCA to 15 PCs for each data set and a cell-cell distance matrix was constructed using the Euclidean distance between cells in this PC space. This distance matrix was used for Silhouette Index computation to measure cluster separation.

### Metrics for assessment of clustering quality

#### Normalized Mutual Information (NMI) to compare cluster labels

The Normalized Mutual Information (NMI) determines the agreement between any two sets of cluster labels 𝒞 and 𝒞′. We compute *NMI*(𝒞, 𝒞′) between 𝒞 and 𝒞^*t*^ as

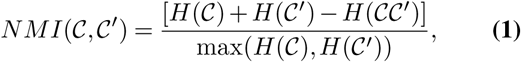

where *H*(𝒞) is the entropy of the clustering 𝒞 (see Chapter 5 of (18) for more information on entropy as a measure of clustering quality). The closer the NMI is to 1.0, the better is the agreement between the two clustering results.

#### Assessment of cluster quality using bootstrapping

We used both (i) Cosine Similarity *cs*_*x,y*_ (19) and (ii) Pearson correlation *r*_*x,y*_ to compute pairwise cell-cell similarities for any pair of single cells (*x, y*) within a cluster *c* according to:

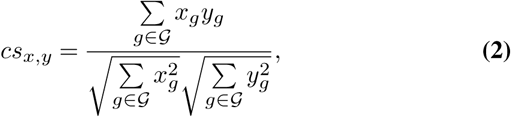

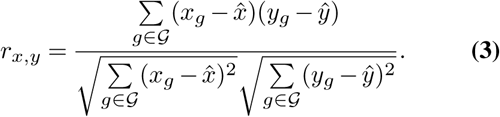

To avoid biases introduced by the feature spaces of the different clustering approaches, both metrics are calculated in the original gene-expression space 𝒢 where *x*_*g*_ represents the expression of gene *g* in cell *x* and *y*_*g*_ represents the expression of gene *g* in cell *y*, respectively. We apply two cut-offs on 𝒢 with respect to the variance of gene-expression (0.5 and 1), thereby neglecting genes that are not likely able to distinguish different clusters from each other. Using bootstrapping, we select 100 genes 100 times from the considered gene-expression space 𝒢 and compute the mean cosine similarity 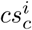 as well as the the mean Pearson correlation 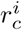 for each cluster *c* ∈ 𝒞 in each iteration *i*:

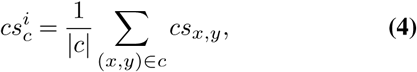

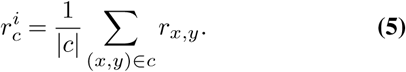

The scores *cs*_*c*_ and *r*_*c*_ are computed for all considered data sets and all three clustering approaches, scConsensus, Seurat and RCA. The closer *cs*_*c*_ and *r*_*c*_ are to 1.0, the more similar are the cells within their respective clusters. Statistical significance is assessed using a one-sided Wilcoxon–Mann–Whitney test.

#### Testing accuracy of cell type assignment on FACS-sorted data

Using the FACS labels as our ground truth cell type assignment, we computed the F1-score of cell type identification to demonstrate the improvement scConsensus achieves over its input clustering results by Seurat and RCA. The F1-score for each cell type *t* is defined as the harmonic mean of precision (*Pre*(*t*)) and recall (*Rec*(*t*)) computed for cell type *t*. In other words,

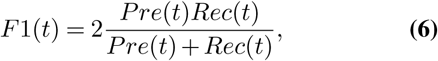

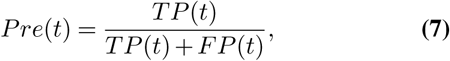

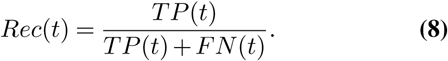

Here, a *TP* is defined as correct cell type assignment, a *FP* refers to a mislabelling of a cell as being cell type *t* and a *FN* is a cell whose true identity is *t* according to the FACS data but the cell was labelled differently.

#### Visualizing scRNA-seq data using UMAP

To visually inspect the scConsensus results, we compute DE genes between every pair of ground-truth clusters and use the union set of those DE genes as the features for PCA. Next, we use the Uniform Manifold Approximation and Projection (UMAP) dimension reduction technique (20) to visualize the embedding of the cells in PCA space in two dimensions.

## Implementation and Availability

scConsensus is implemented in **R** and is freely available on GitHub at https://github.com/prabhakarlab/scConsensus. All data used is available on Zenodo (doi: 10.5281/zenodo.3637700). For generation of the contingency table, scConsensus uses the **R** packages mclust, Complex-Heatmap, circlize and reshape2. For DE gene refinement and cell clustering, scConsensus uses the flash-Clust, calibrate, WGCNA, edgeR, circlize and ComplexHeatmap packages. Silhouette Index was computed using the silhouette function in the cluster package, while Normalized Mutual Information was computed using the NMI function in the aricode package. scConsensus has been tested with **R** versions ≥ 3.6.

## Results

### scConsensus - A hybrid approach for clustering single cell data

scConsensus is a general **R** framework offering a workflow to combine complementary results of two different clustering approaches. Here, we benchmarked scConsensus to combine both unsupervised and supervised scRNA-seq clusters computed with Seurat and RCA on five 10X CITE-Seq data sets and on one 10X GemCode data set.

### scConsensus produces clusters that are more consistent with antibody-derived clusters

We used the Antibody-derived Tag (ADT) signal of the five considered CITE-seq data sets to generate a ground truth clustering for all considered samples (Fig. 2a). Next, we compute all differentially expressed (DE) genes between the antibody based clusters using the scRNA-seq component of the data. As shown in Fig. 2b (Sup. Fig. S2), the expression of DE genes is cluster-specific, thereby showing that the antibody-derived clusters are separable in gene expression space. Therefore, these DE genes are used as a feature set to evaluate the different clustering strategies.

**Fig. 2.**
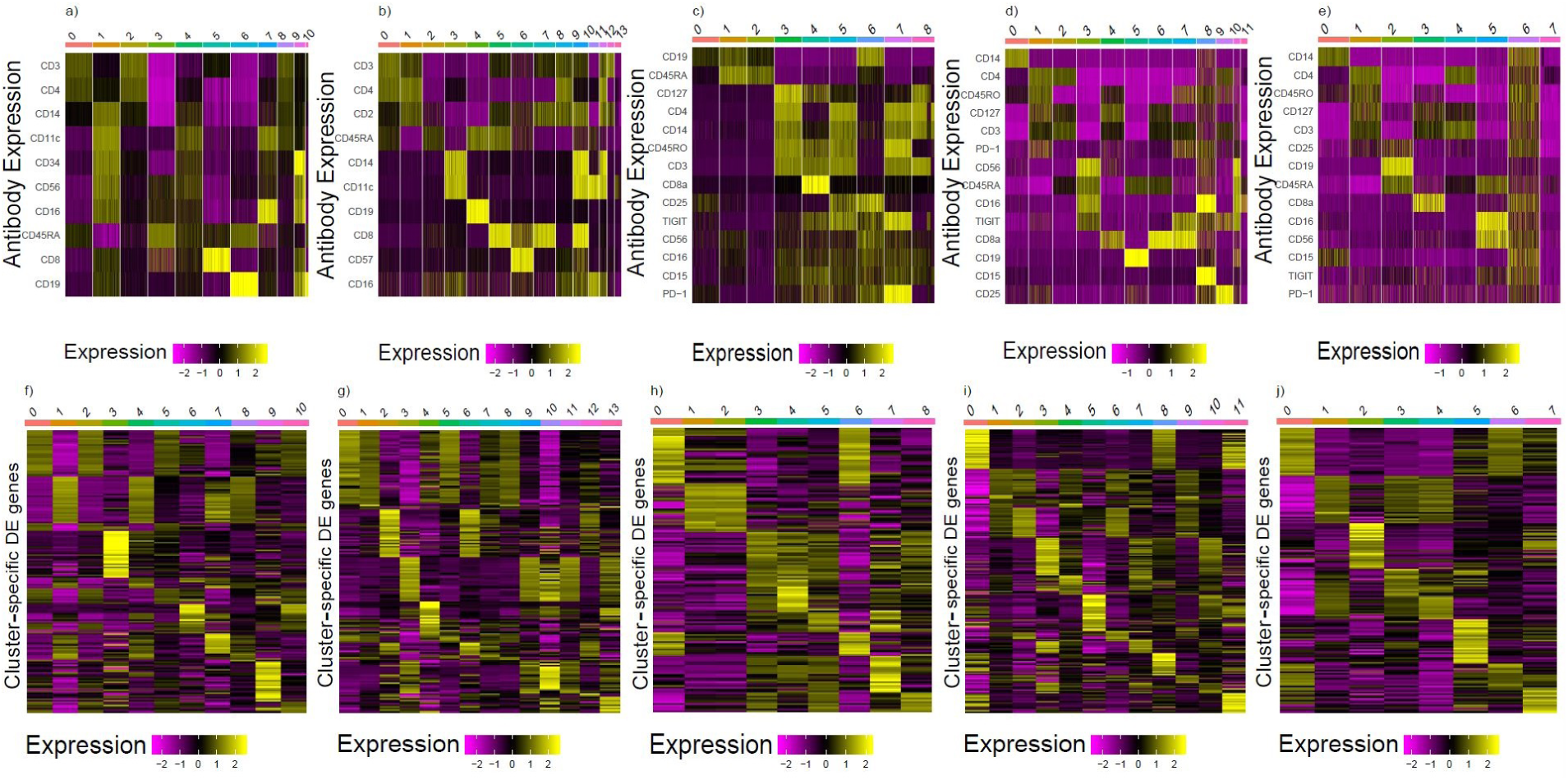
In the heatmaps (a-e), we show ADT cluster specific antibody signal across five CITE-seq data sets per cell, whereas the heatmaps (f-j) show the expression of the top 30 differentially expressed genes averaged across all cells per cluster.

Here, we assessed the agreement of scConsensus, the supervised RCA and the unsupervised Seurat clusters with the antibody-based single-cell clusters in terms of Normalized Mutual Information (NMI), a score quantifying similarity with respect to the cluster labels. scConsensus is more consistent with antibody-based clusters than both Seurat and RCA on all but the MALT data set, where scConsensus is the second best approach (Fig. 3a). Interestingly, we find that there is no consistency in performance for the second best method, depending on the data set this is either the supervised (RCA) or the unsupervised (Seurat) clustering method.

**Fig. 3.**
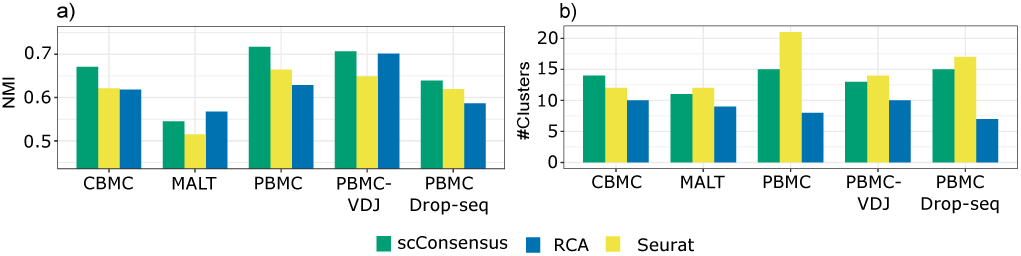
a) Normalized Mutual Information (NMI) quantifies the agreement between the ground truth (antibody based clustering) and the transcriptomic clusters computed using scConsensus, Seurat, and RCA. b) Number of clusters detected using either scConsensus, Seurat, or RCA.

Applying scConsensus, Seurat and RCA to five CITE-seq data sets results suggests that RCA tends to find more clusters than scConsensus and Seurat. On average scConsensus leads to more clusters than Seurat but to less clusters than RCA (Fig. 3b).

For a visual inspection of these clusters, we provide UMAPs visualizing the clustering results in the ground truth feature space based on DE genes computed between ADT clusters, with cells being colored according to the cluster labels provided by one of the tested clustering methods (Sup. Fig S5-S8). In Figure 4, we show the respective UMAPs for the PBMC data set. By visually comparing the UMAPs, we find for instance that Seurat cluster 3 (Fig. 4b), corresponds to the two antibody clusters 4 and 7 (Fig. 4a). In contrast to the unsupervised results, this separation can be seen in the supervised RCA clustering (Fig. 4c) and is correctly reflected in the unified clustering by scConsensus (Fig. 4d). Another illustration for the performance of scConsensus can be found in the supervised clusters 3, 4, 9, and 12 (Fig. 4c), which are largely overlapping. In the ADT cluster space, the corresponding cells should form only one cluster (Fig. 4a). Here scConsensus picks up the cluster information provided by Seurat (Fig. 4b), which reflects the ADT labels more accurately (Fig. 4d). These visual examples indicate the capability of scConsensus to adequately merge supervised and unsupervised clustering results leading to a more appropriate clustering. Similar examples can be found for the other data sets (CBMC, PBMC Drop-Seq, MALT and PBMC-VDJ) in Sup. Fig. S5-S8.

**Fig. 4.**
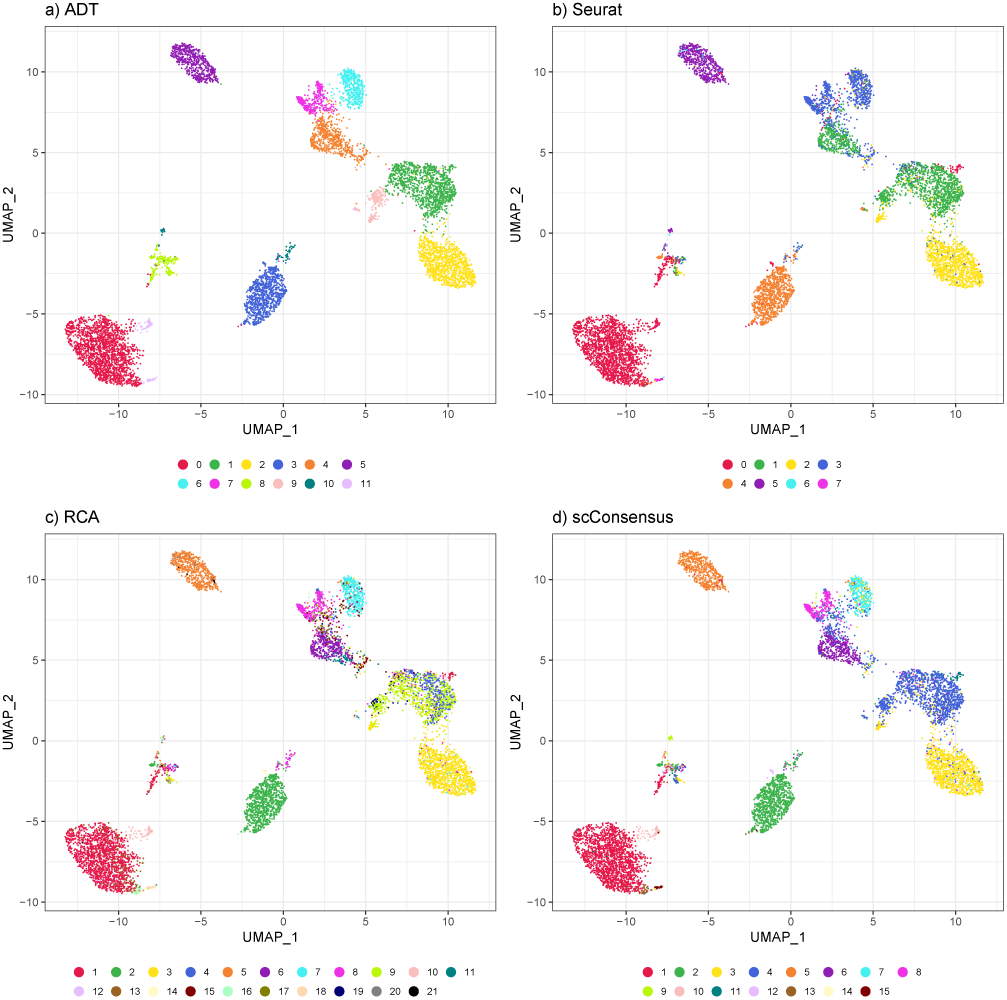
UMAPs anchored in DE-gene space computed for a ADT-based clustering of the PBMC data set colored by cluster IDs obtained for (a) ADT data, (b) Seurat clusters, (c) RCA and (d) scConsensus.

In addition to the NMI, we assessed the performance of scConsensus in yet another complementary fashion. We quantified the quality of clusters in terms of within-cluster similarity in gene-expression space using both Cosine similarity and Pearson correlation. Using bootstrapping, we find that scConsensus consistently improves over clustering results from RCA and Seurat(Sup. Fig. S3 and Sup. Fig S4) supporting the benchmarking using NMI. While the advantage of this comparisons is that it is free from biases introduced through antibodies and cluster method specific feature spaces, one can argue that using all genes as a basis for comparison is not ideal either. However, paired with bootstrapping, it is one of the fairest and most unbiased comparisons possible. A similar approach has been taken previously by (21) to compare the expression profiles of CD4+ T-cells using bulk RNA-seq data. Analogously to the NMI comparison, the number of resulting clusters also does not correlated to our performance estimates using Cosine similarity and Pearson correlation.

### scConsensus accurately reproduces FACS-sorted PBMC cell type labels

Using data from (11), we clustered cells using Seurat and RCA, as in the previous examples. After annotating the clusters, we provided scConsensus with the two clustering results as inputs and computed the F1-score of cell type assignment using the FACS labels as ground truth.

Fig. 5a shows the mean F1-score for cell type assignment using scConsensus, Seurat and RCA, with scConsensus achieving the highest score. Fig. 5b depicts the F1 score in a cell type specific fashion. Fig. 5 shows the visualization of the various clustering results using the FACS labels, Seurat, RCA and scConsensus. A striking observation is that CD4 T Helper cells could neither be captured by RCA nor by Seurat, and hence also not by scConsensus. Fig. 5b also illustrates that scConsensus does not hamper with and can even slightly further improve the already reliable detection of B cells, CD14+ Monocytes, CD34+ cells (Progenitors) and Natural Killer (NK) cells even compared to RCA and Seurat. Importantly, scConsensus is able to isolate a cluster of Regulatory T cells (T Regs) that was not detected by Seurat but was pinpointed through RCA (5c). The scConsensus approach extended that cluster leading to an F1-score of 0.6 for T Regs. However, the cluster refinement using DE genes lead not only to an improved result for T Regs and CD4 T-Memory cells, but it also resulted in a slight drop in performance of scConsensus compared to the best performing method for CD4+ and CD8+ T-Naive as well as CD8+ T-Cytotoxic cells. As indicated by a UMAP representation colored by the FACS labels (Fig.5a), this is likely due to the fact that all immune cells are part of one large immune-manifold, without clear cell type boundaries, at least in terms of scRNA-seq data.

**Fig. 5.**
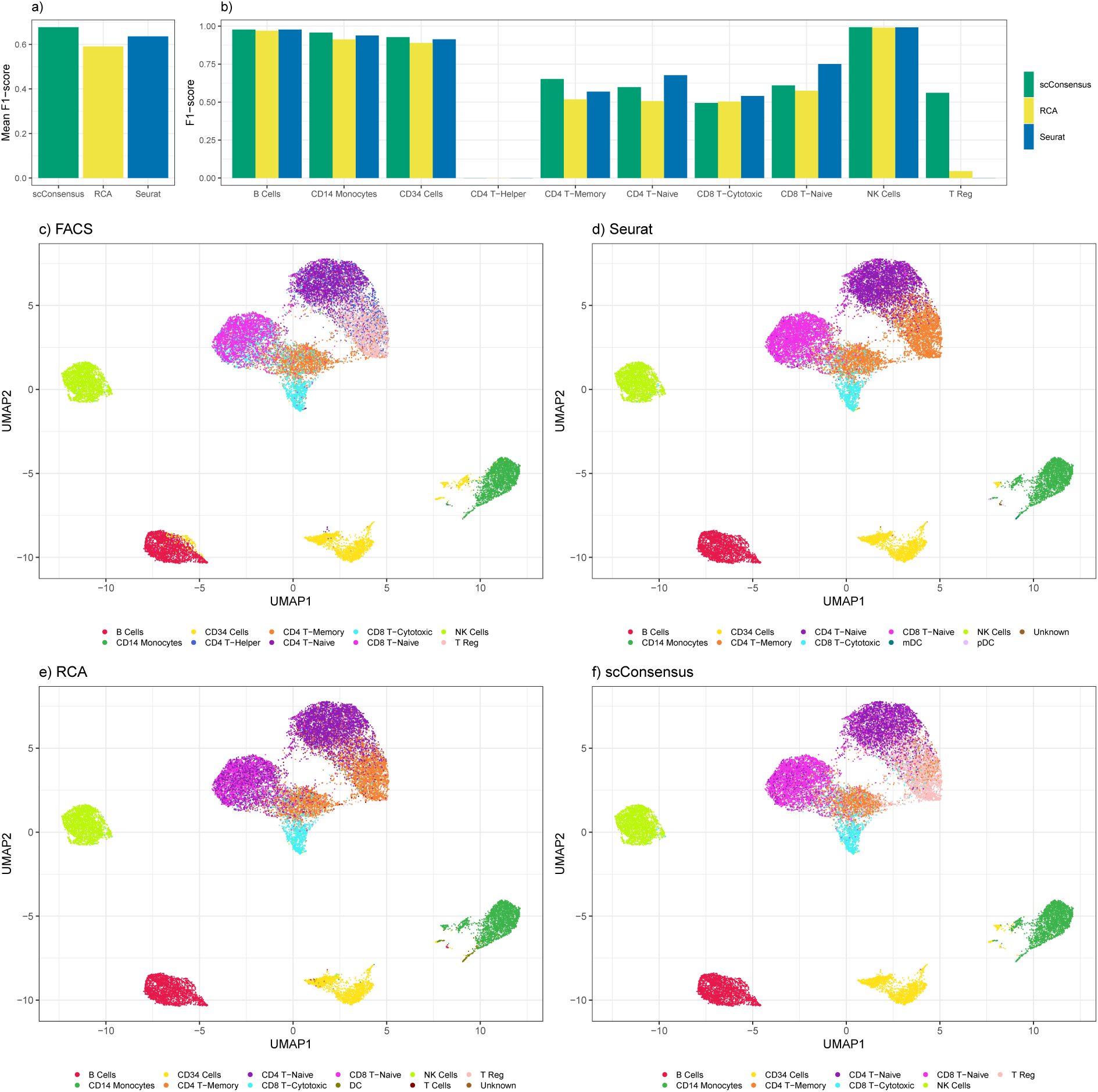
Performance assessment of cell type assignment on FACS sorted PBMC data using scConsensus, Seurat and RCA for clustering. The mean F1-score, which is the harmonic mean of precision and recall, is shown across all cell types in (a) while cell type specific scores are shown in (b). UMAPs (c-f) are anchored in the DE-gene space computed for FACS-based clustering with the UMAP in (c) being colored according to the FACS labels. For the remaining UMAPs, the color code is based on the indicated clustering methods, that is (d) Seurat, (e) RCA and (f) scConsensus.

Another example for the applicability of scConsensus is the accurate annotation of a small cluster shown in Fig. 5 to the left of the CD14 Monocytes cluster. Using Seurat, the majority of those cells are annotated as stem cells, while a minority are annotated as CD14 Monocytes (Fig. 5b). RCA annotates these cells exclusively as CD14+ Monocytes (Fig. 5c). However, according to FACS data (Fig. 5a) these cells are actually CD34+ (Progenitor) cells, which is well reflected by scConsensus (Fig. 5d).

Overall, these examples demonstrate the power of combining reference-based clustering with unsupervised clustering and showcase the applicability of scConsensus to identify and cluster even closely-related sub-types in scRNA-seq data.

## Discussion

Many different approaches have been proposed to solve the single-cell clustering problem, in both unsupervised (3) and supervised (5) ways. However, all approaches have their own advantages and disadvantages and do not necessarily lead to similar results, as exemplified in Sup. Fig. 1. While bench-marking scConsensus we also found that there is no consistent ranking between the tested supervised and unsupervised approaches. On some data sets, e.g. the FACS sorted PBMC data shown in Fig. 5, the unsupervised Seurat performs better than the supervised RCA, while the latter achieves better performance than Seurat on the CITE-seq data sets (Fig. 3). In fact, this observation stresses that there is no ideal approach for clustering and therefore also motivates the development of a consensus clustering approach. With scConsensus we propose a computational strategy to find a consensus clustering that provides the best possible cell type separation for a single-cell data set.

scConsensus builds on known intuition about single-cell RNA sequencing data, i.e. homogeneous cell types will have consistent differentially expressed marker genes when compared with other cell types. scConsensus computes DE gene calls in a pairwise fashion, that is comparing a distinct cluster against all others. Together with a constant number of DE genes considered per cluster, scConsensus gives equal weight to rare sub-types, which may otherwise get absorbed into larger clusters in other clustering approaches. We have demonstrated this using a FACS sorted PBMC data set and the loss of a cluster containing regulatory T-cells in Seurat compared to scConsensus.

A major feature of the scConsensus workflow is its flexibility - it can help leverage information from any two clustering results. Here, we focus on Seurat and RCA, two complementary methods for clustering and cell type identification in scRNA-seq data. However, the intuition behind scConsensus can be extended to any two clustering approaches. For example, even using the same data, unsupervised graph-based clustering and unsupervised hierarchical clustering can lead to very different cell groupings. Upon encountering this issue, users typically tend to pick the clustering result that agrees best with their domain knowledge, while completely ignoring the information provided by the other clustering. Thus, we propose scConsensus as a valuable, easy and robust solution to the problem of integrating different clustering results to achieve a more informative clustering.

## Conclusion

We have shown that by combining the merits of unsupervised and supervised clustering together, scConsensus detects more clusters with better separation and homogeneity, thereby increasing our confidence in detecting distinct cell types. As scConsensus is a general strategy to combine clustering methods, it is apparent that scConsensus is not restricted to scRNA-seq data alone. Any multidimensional singlecell assay whose cell clusters can be separated by differential features can leverage the functionality of our approach. For instance, for single-cell ATAC sequencing data, there are various clustering approaches available that lead to different clustering results (22). scConsensus could be used out of the box to consolidate these clustering results and provide a single, unified clustering result. Therefore, we believe that the clustering strategy proposed by scConsensus is a valuable contribution to the computational biologist’s toolbox for the analysis of single-cell data.

## Acknowledgements

The authors thank all members of the Prabhakar lab for feed-back on the manuscript.

## Author contributions

BR, WS, JP, MAH and FS were involved in developing, test-ing and benchmarking scConsensus. NAR and JT developed the immune reference panel. BR and FS wrote the manuscript. BR, FS and SP edited and reviewed the manuscript.

