## Supplementary Information for "scConsensus: combining supervised and unsupervised clustering for cell type identification in single-cell RNA sequencing data"

**Supplement *scConsensus***

### Supplementary Figures

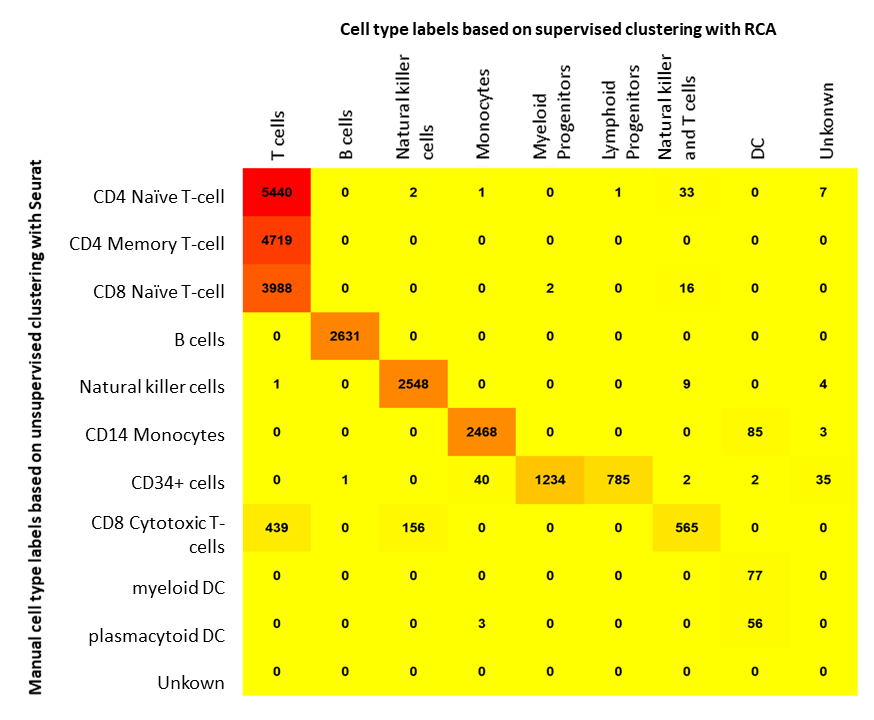

**Supplementary Figure S1:** Consensus matrix of cell type annotation for PBMC data. Columns show cell type labels based on supervised clustering with RCA, rows show cell type labels based on unsupervised clusters using Seurat and a subsequent marker based annotation. The most informative labelling could be obtained by combining both annotations, e.g. the immune cell annotation derived from Seurat clustering and the Progenitor annotation derived by RCA. Figure based on data from Zheng et al. (2017).

A) CBMC

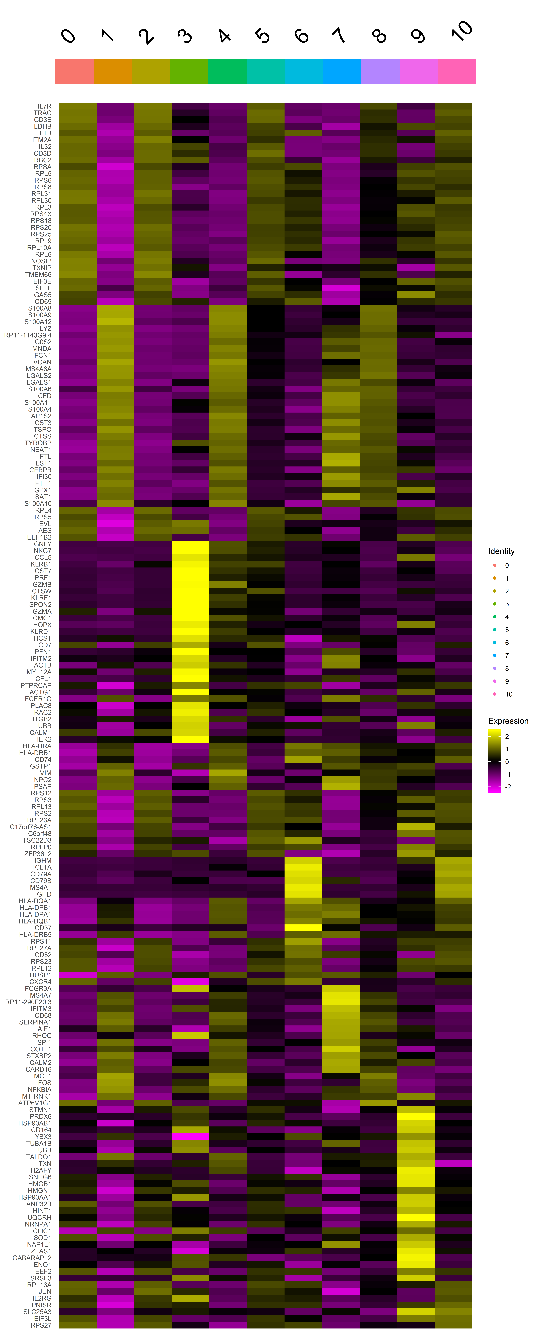

B) PBMC Drop-Seq

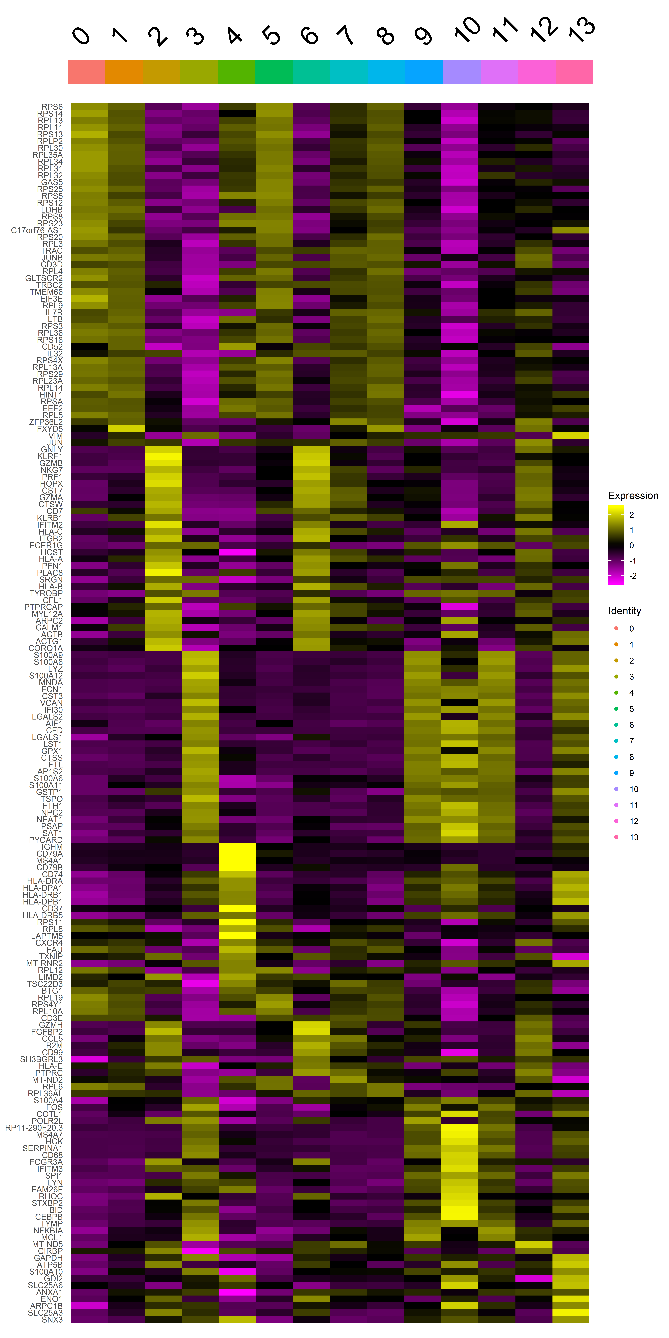

C) MALT

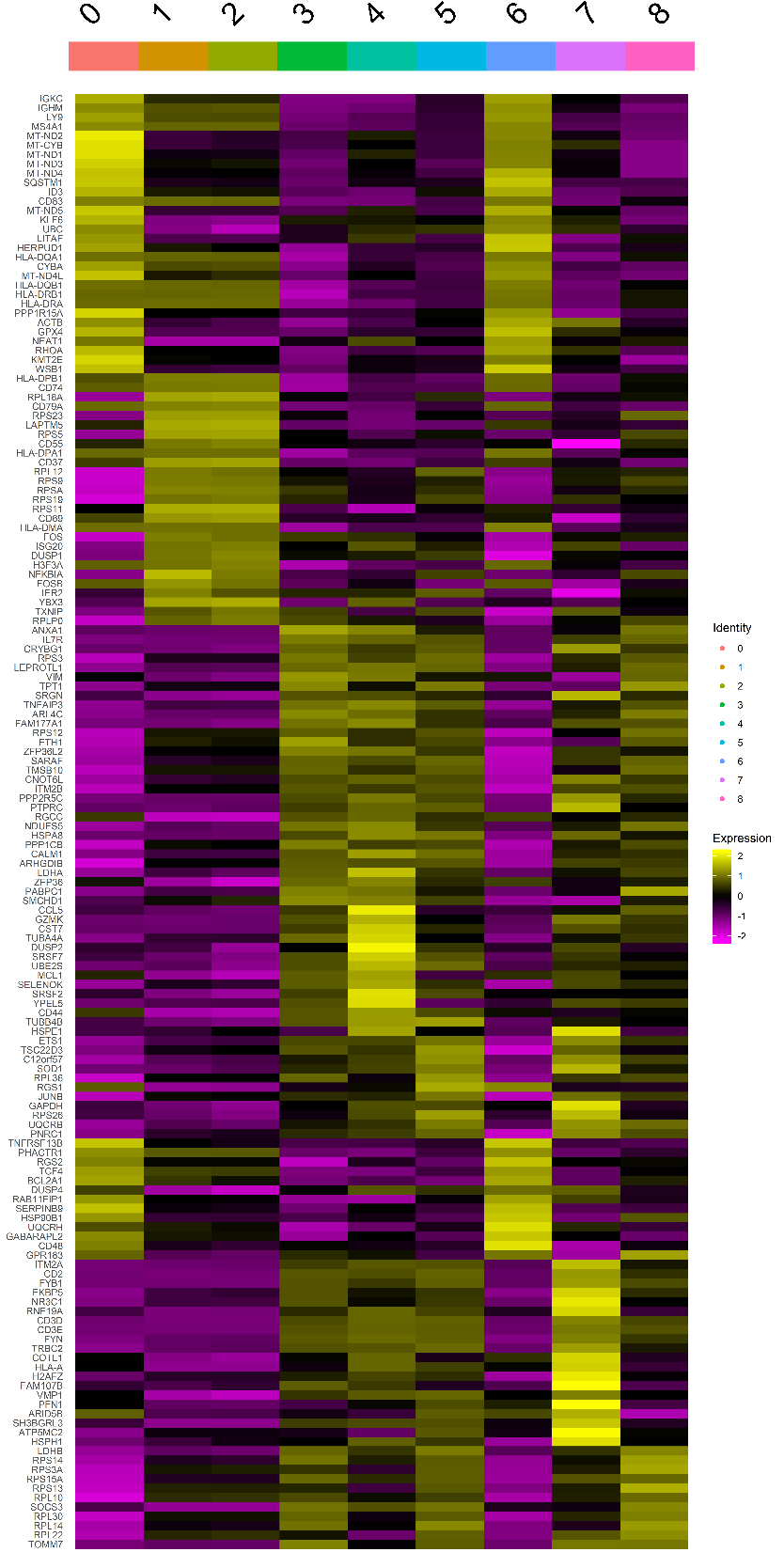

D) PBMC

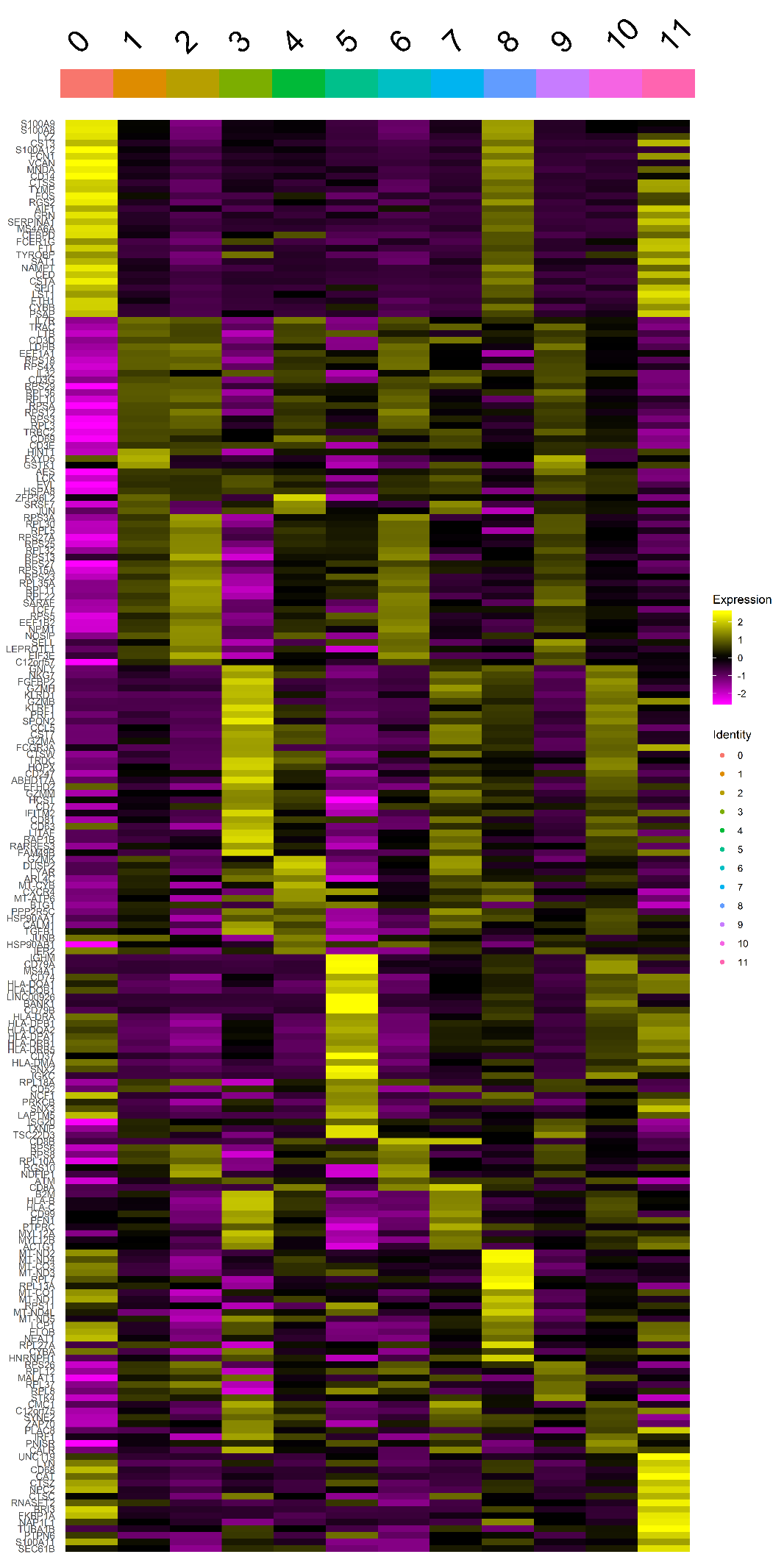

E) PBMC-VDJ

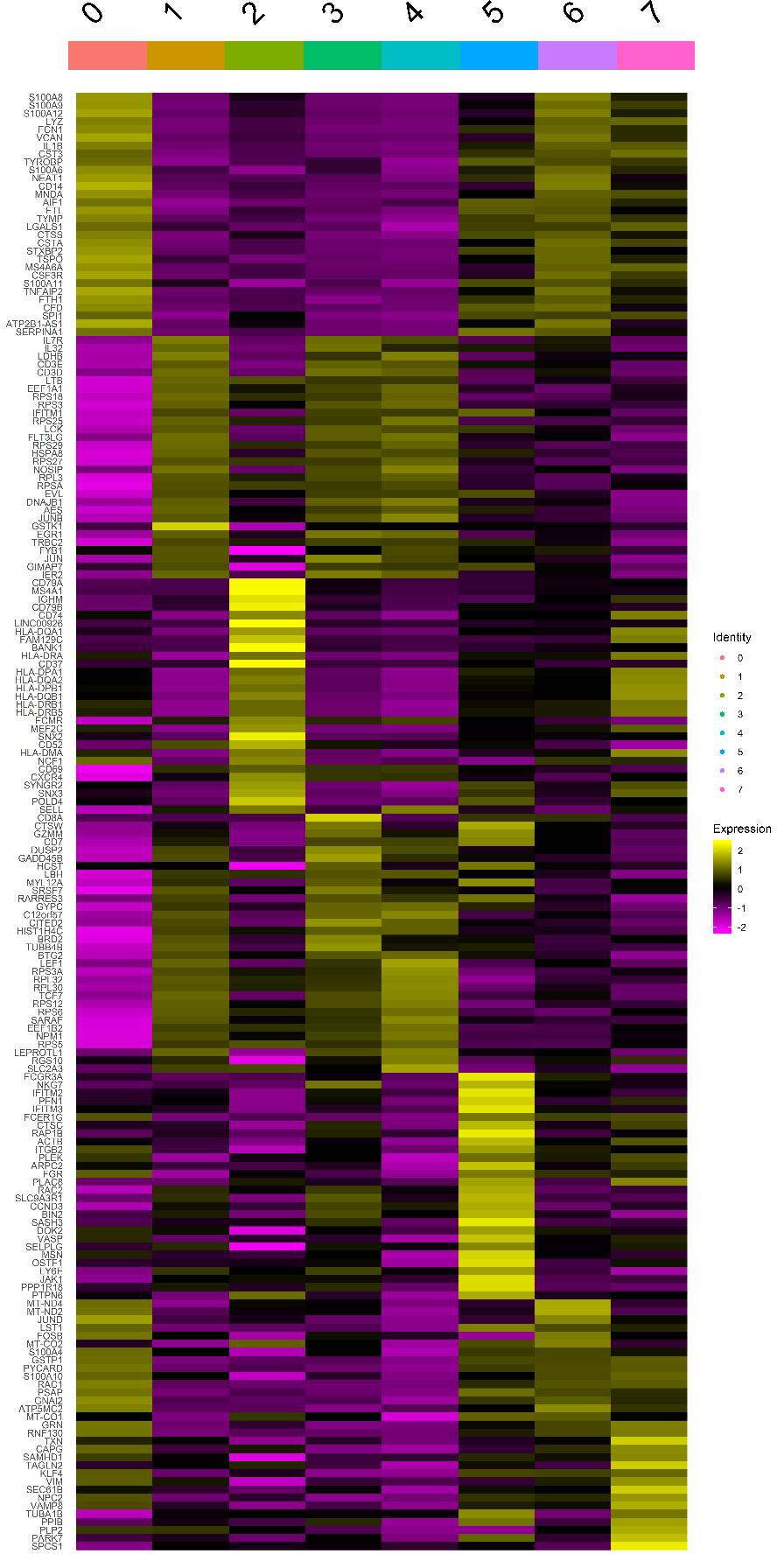

**Supplementary Figure S2:** Heat maps showing cluster-specific DE gene expression for A) CBMC B) PBMC Drop-Seq C) MALT D) PBMC E) PBMC-VDJ

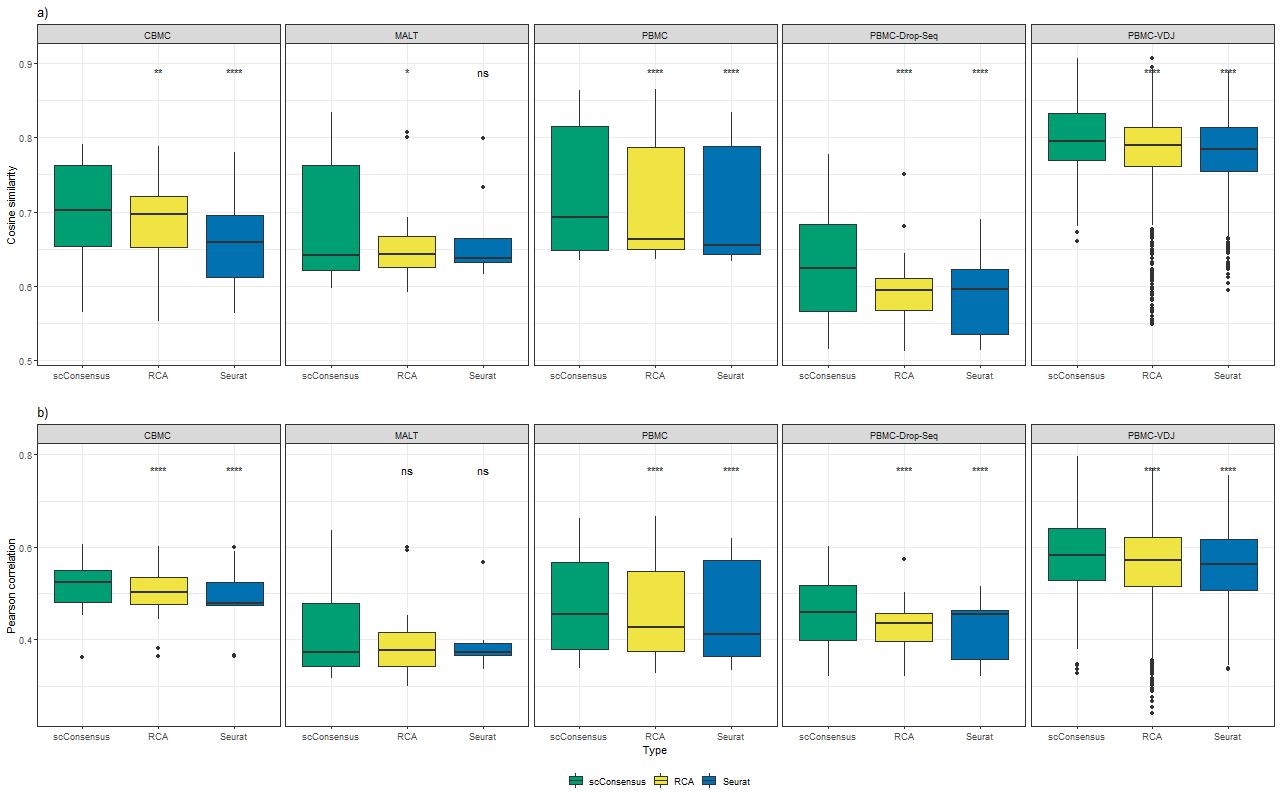

**Supplementary Figure S3:** Cosine and Pearson correlation for bootstrapping on the gene-expression space using a variance threshold of 0.5

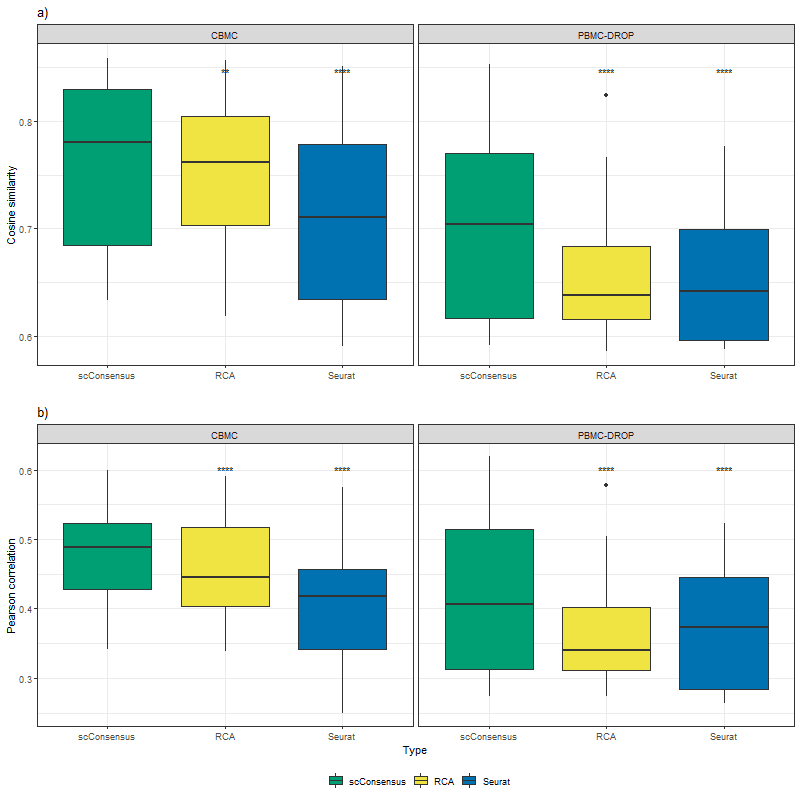

**Supplementary Figure S4:** Cosine and Pearson correlation for bootstrapping on the gene-expression space using a variance threshold of 1.0

A) Ground Truth

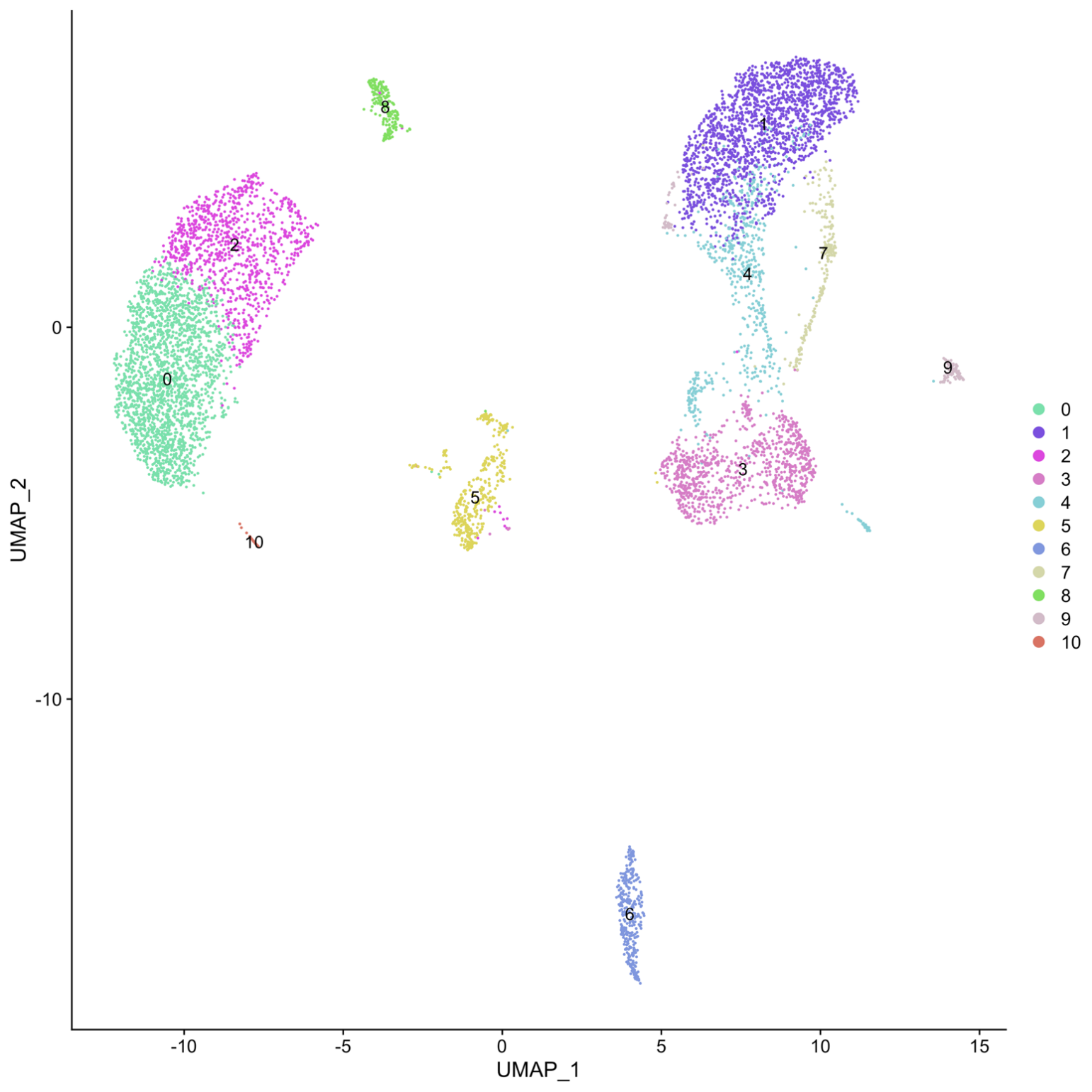

B) scConsensus

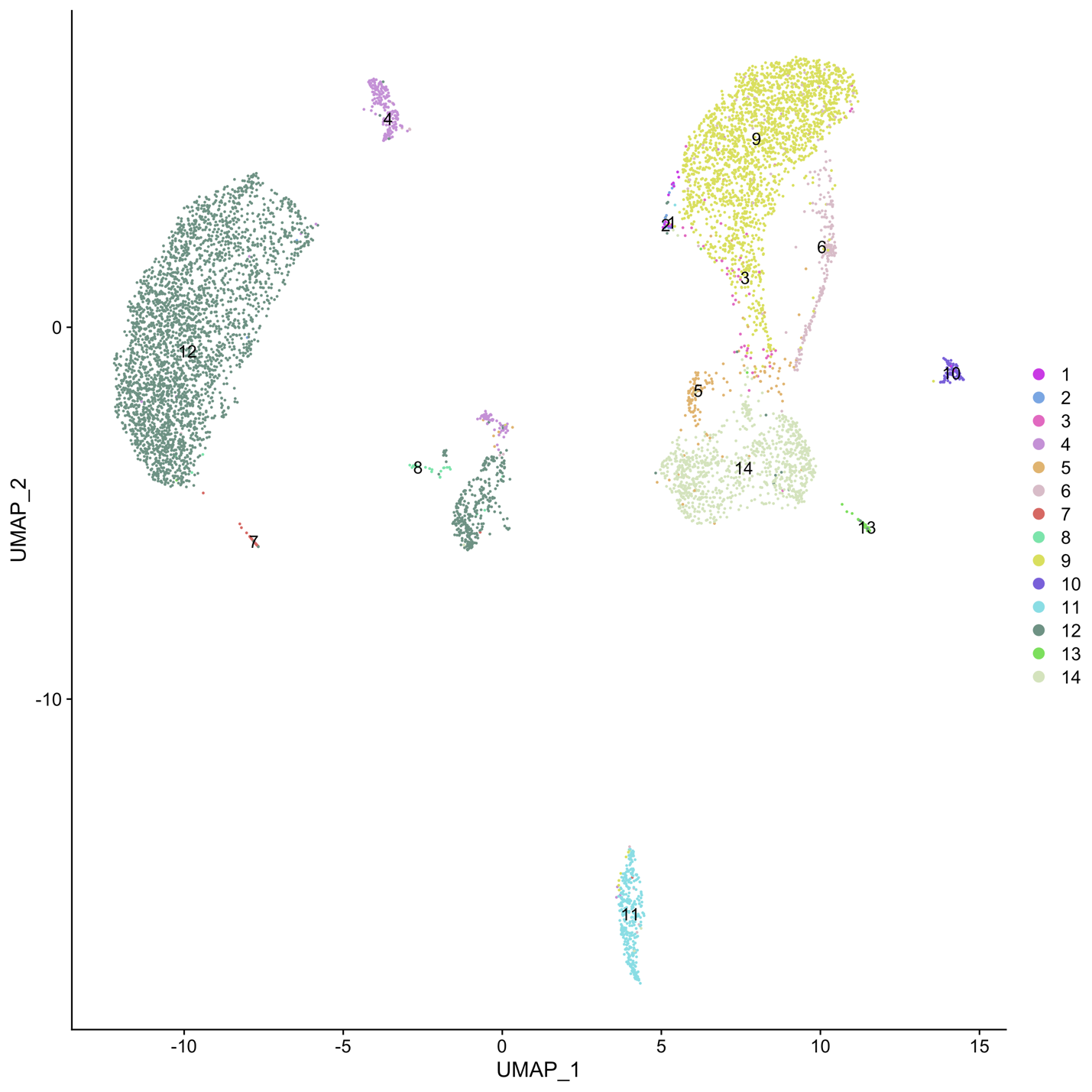

C) Seurat

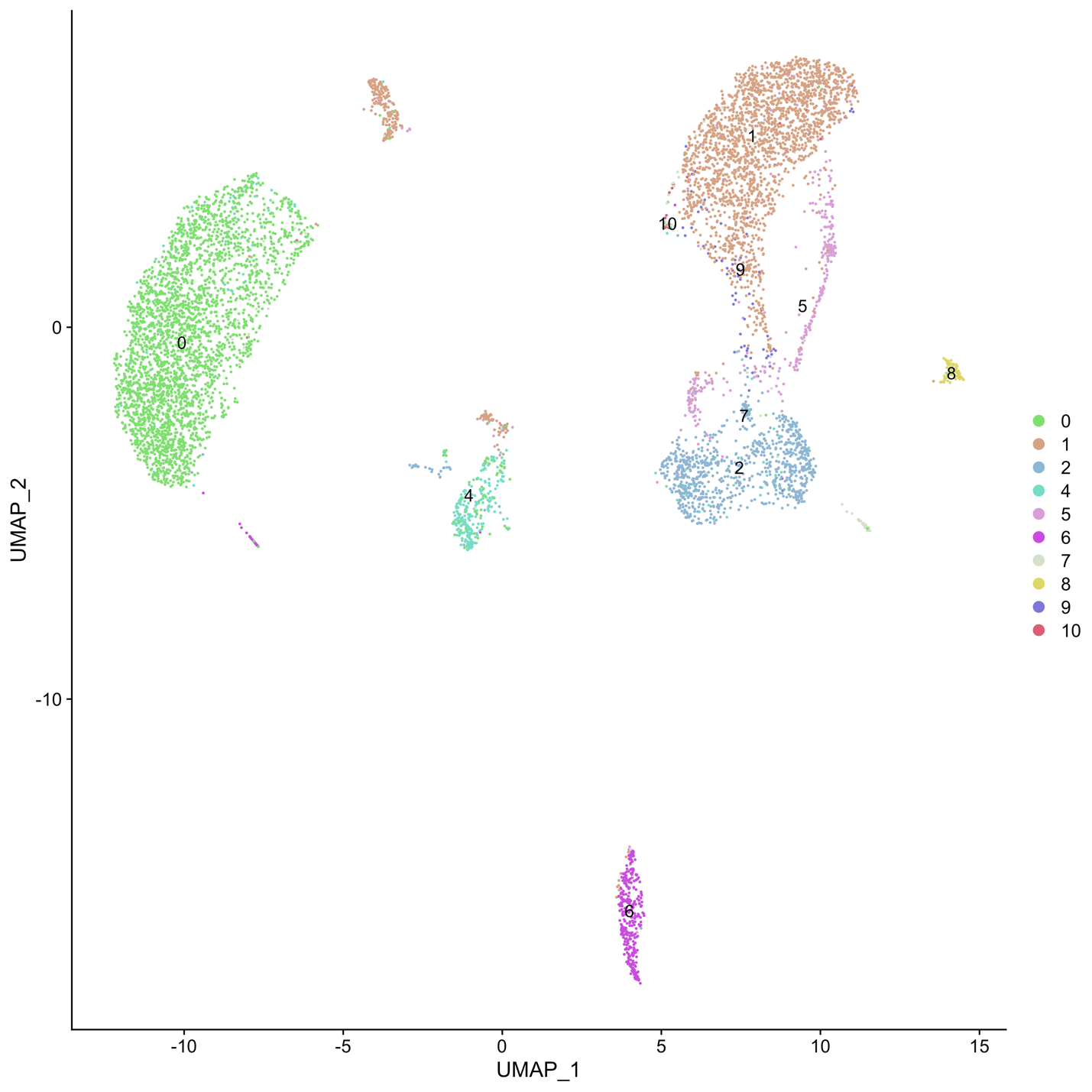

D) RCA

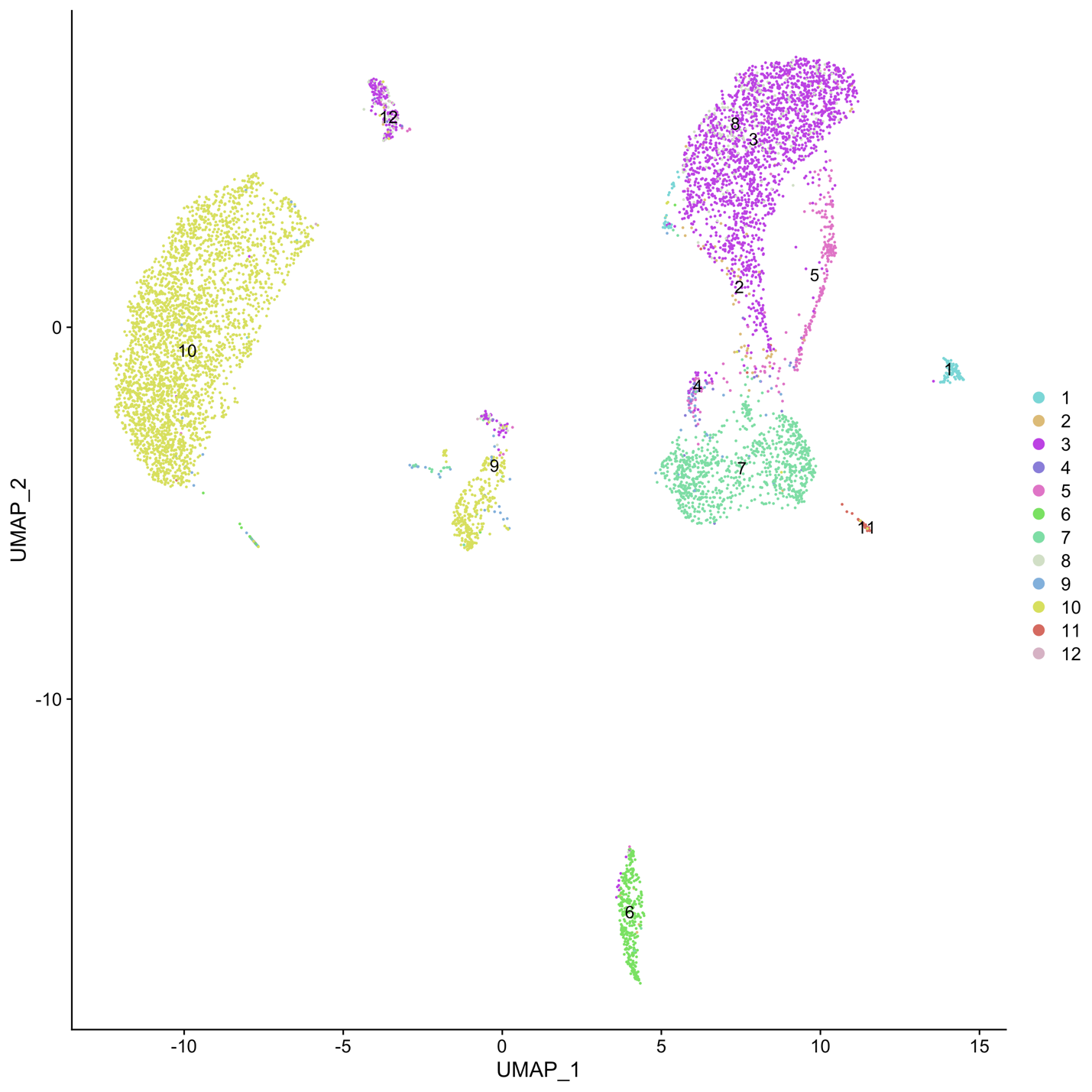

**Supplementary Figure S5:** UMAP visualizations of cells in ADT space colored by A) Antibody-cluster ground truth B) scConsensus C) Seurat and D) RCA in the antibody expression space for the CBMC Dataset

A) Ground Truth

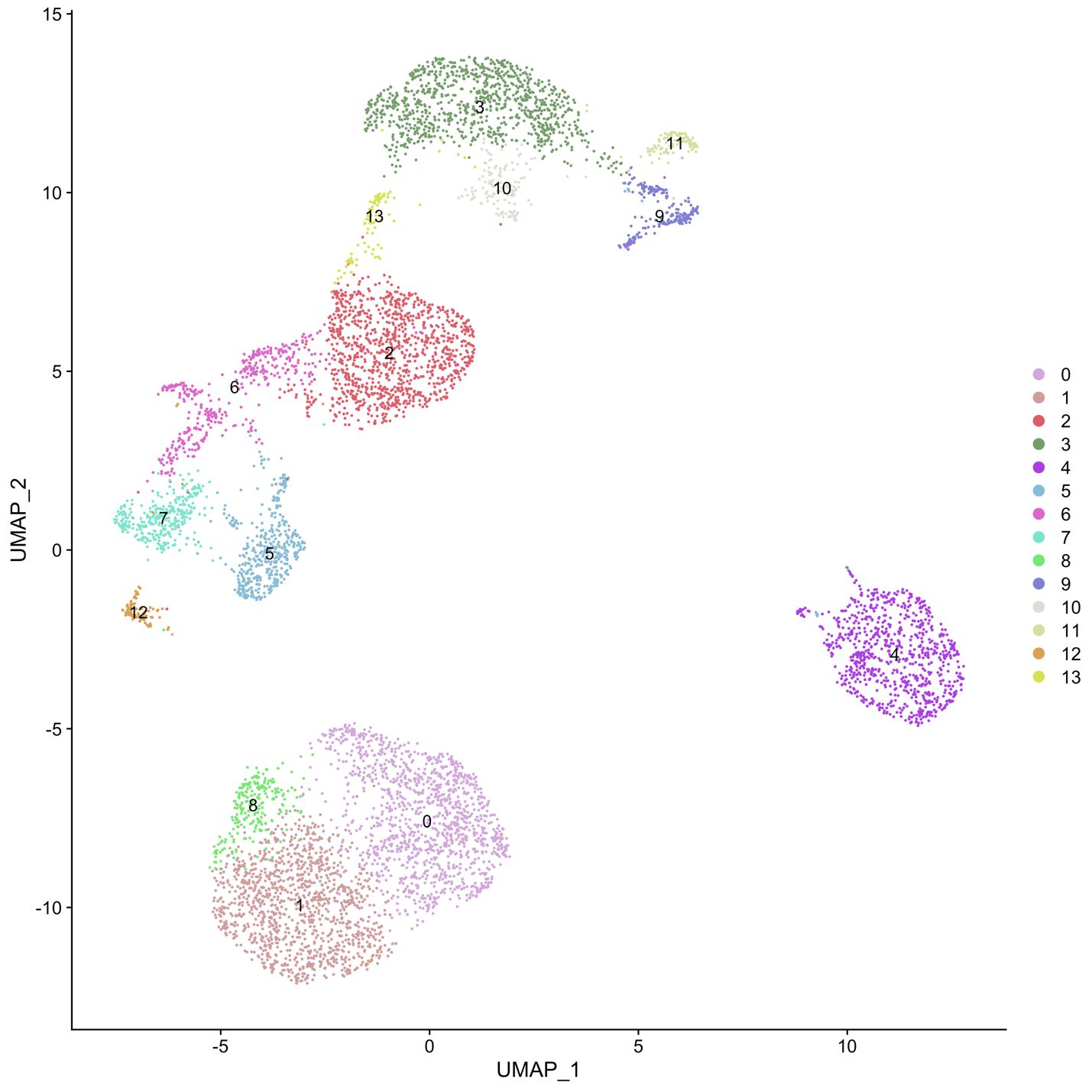

B) scConsensus

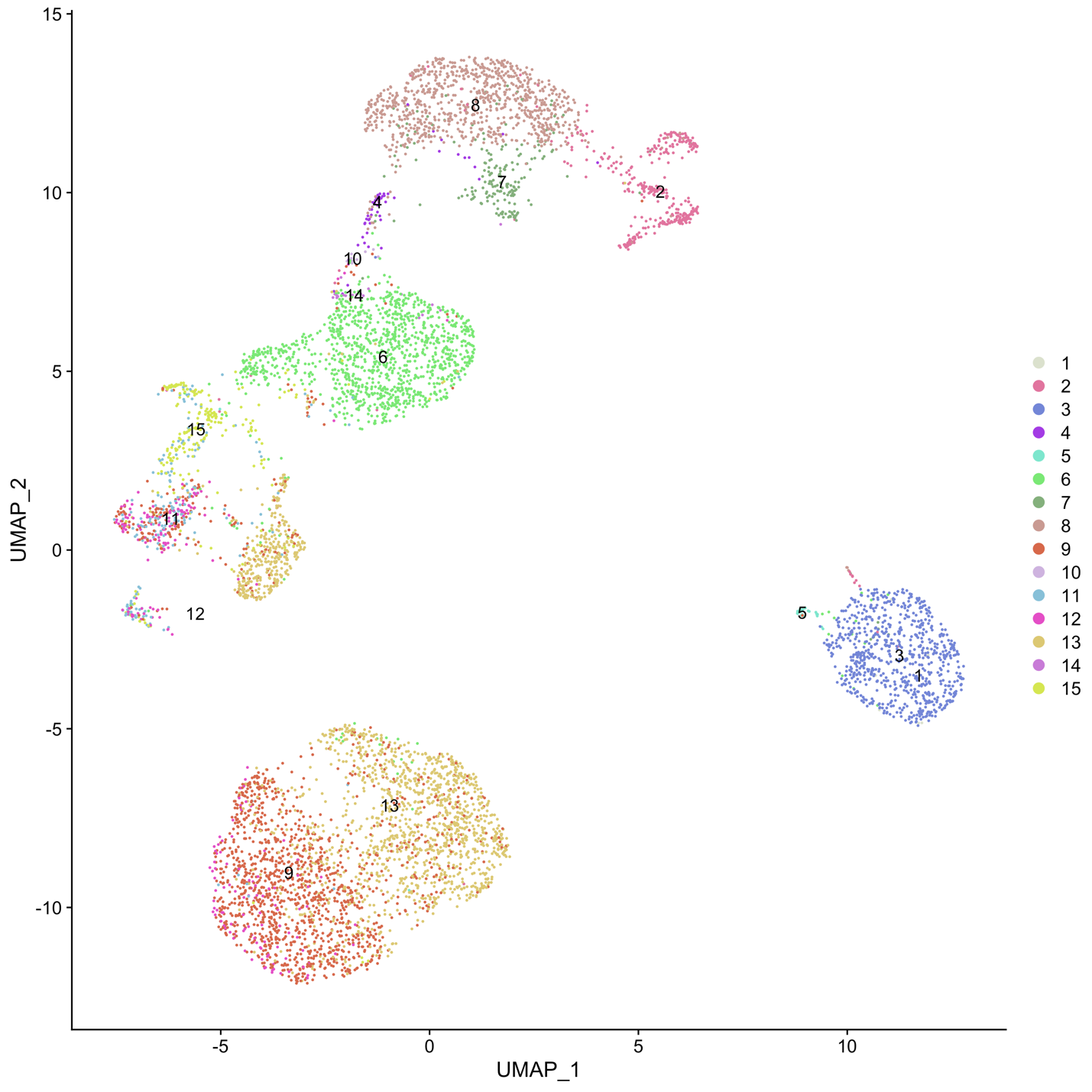

C) Seurat

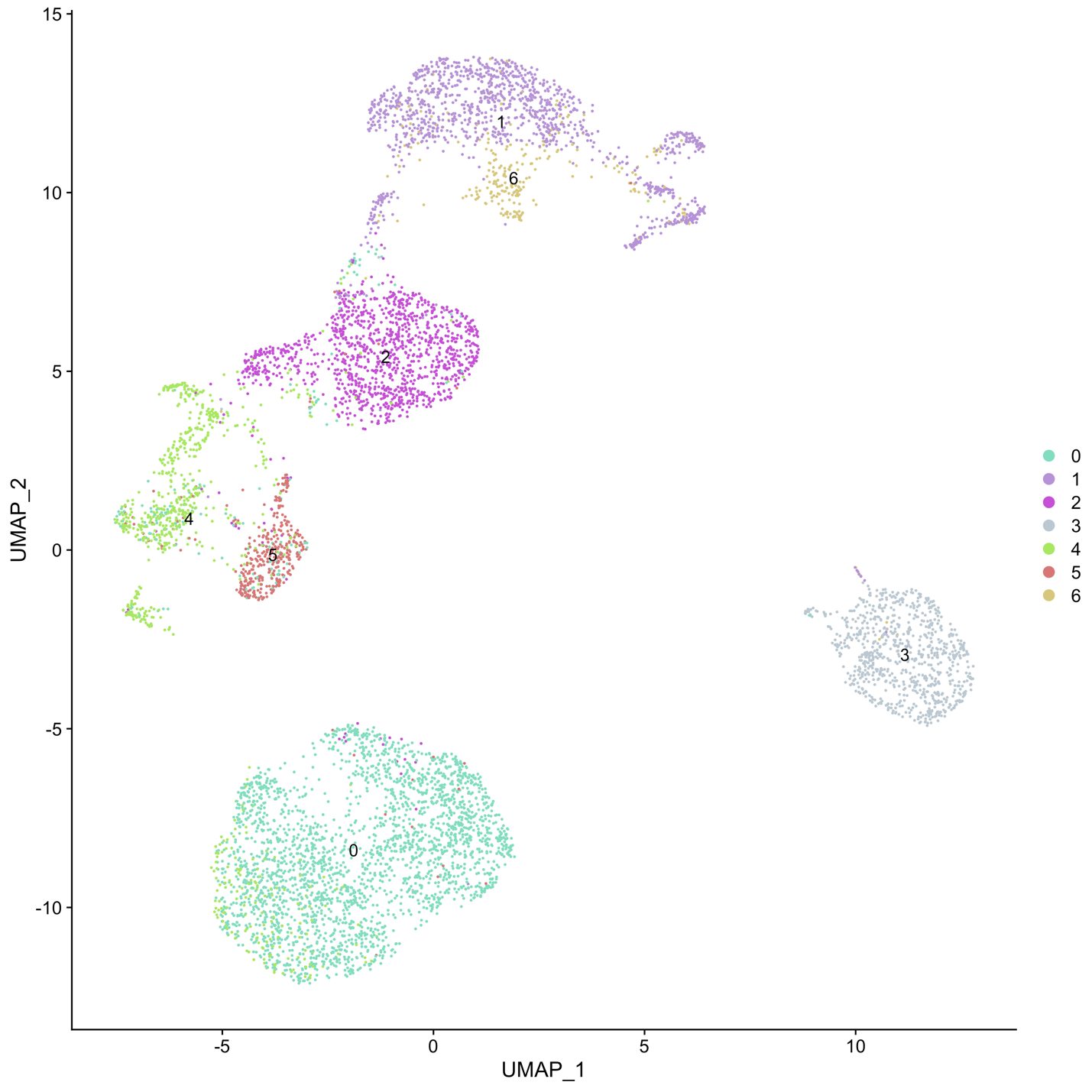

D) RCA

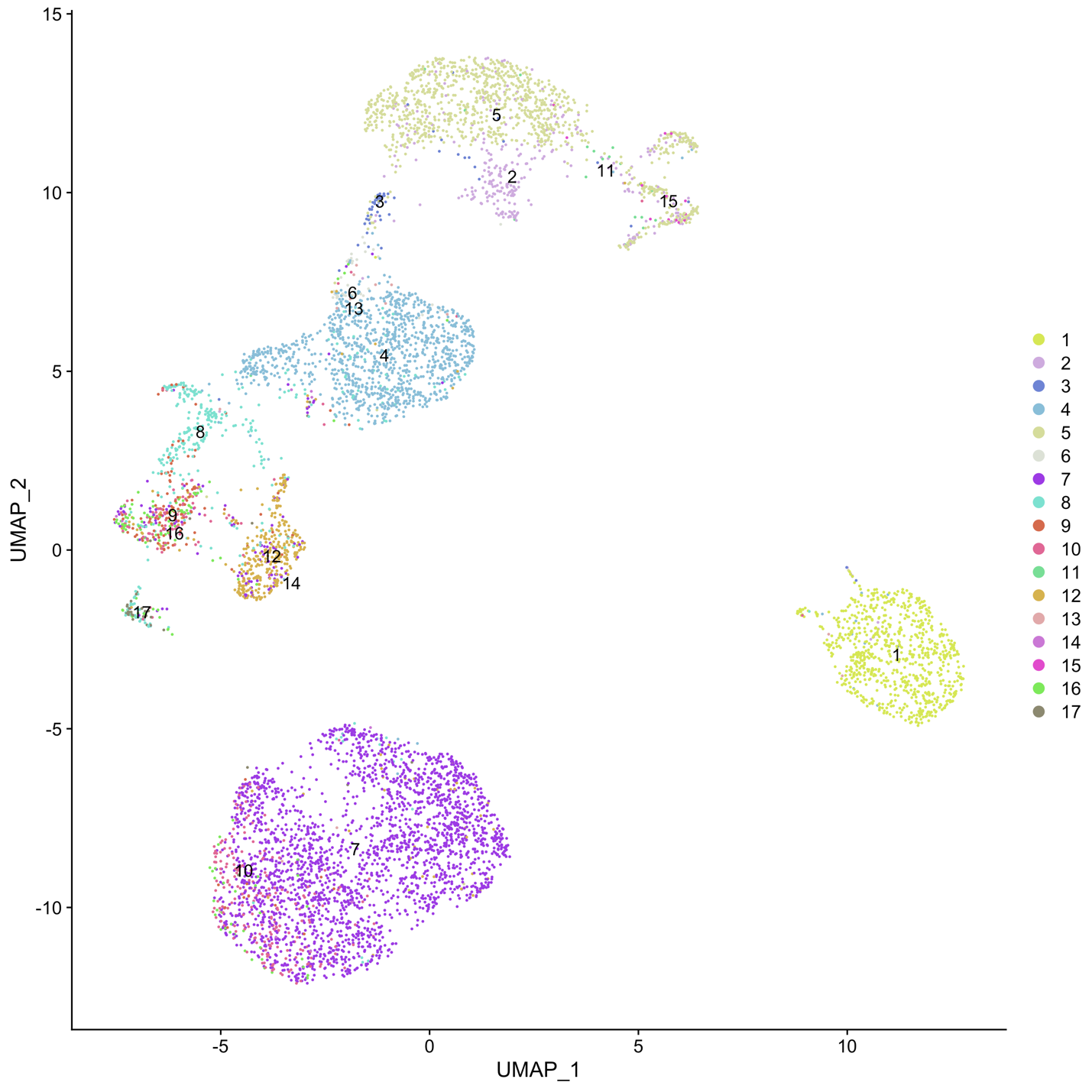

**Supplementary Figure S6:** UMAP visualizations of cells in ADT space colored by A) Antibody-cluster ground truth B) scConsensus C) Seurat and D) RCA in the antibody expression space for the PBMC Drop-Seq dataset

A) Ground Truth

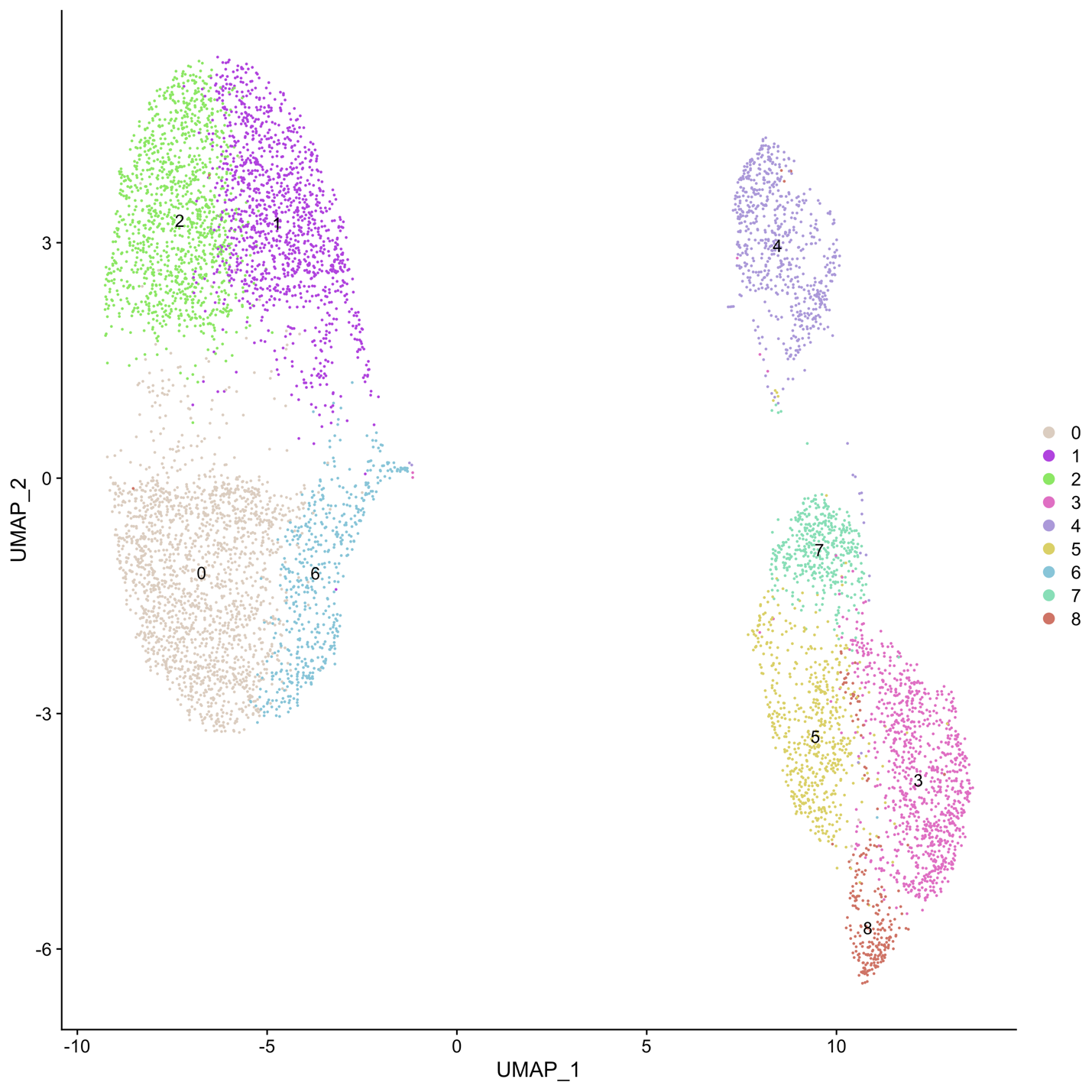

B) scConsensus

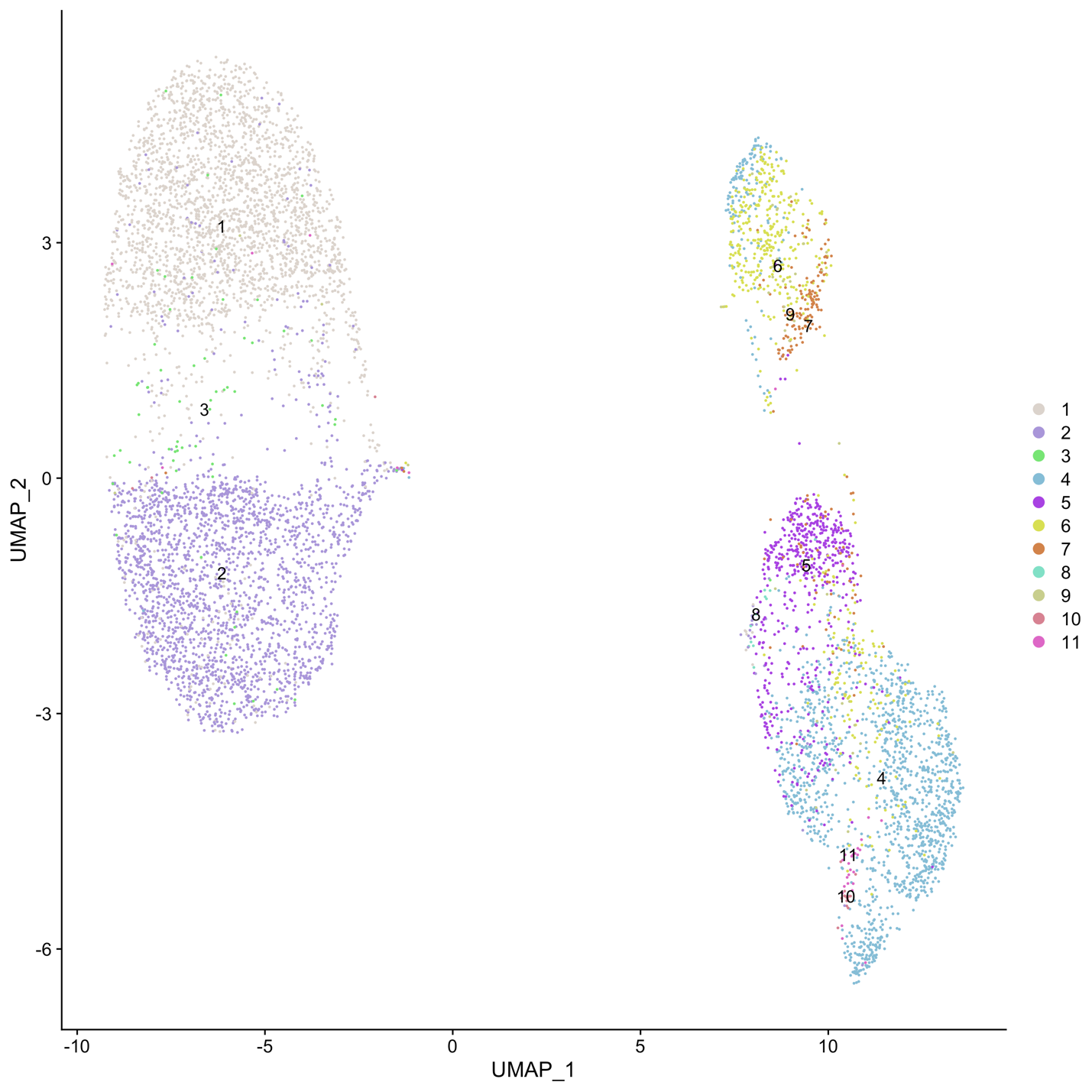

C) Seurat

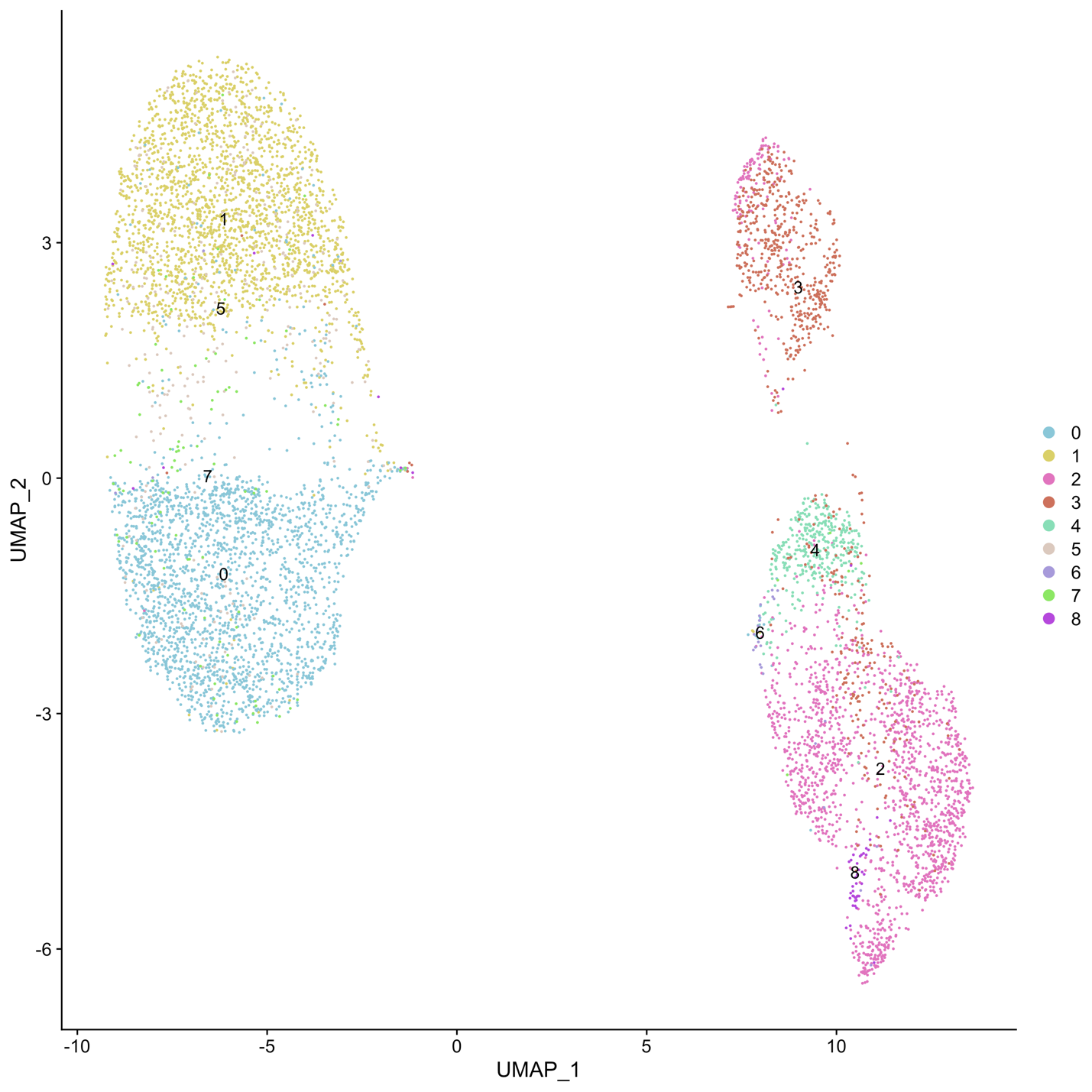

D) RCA

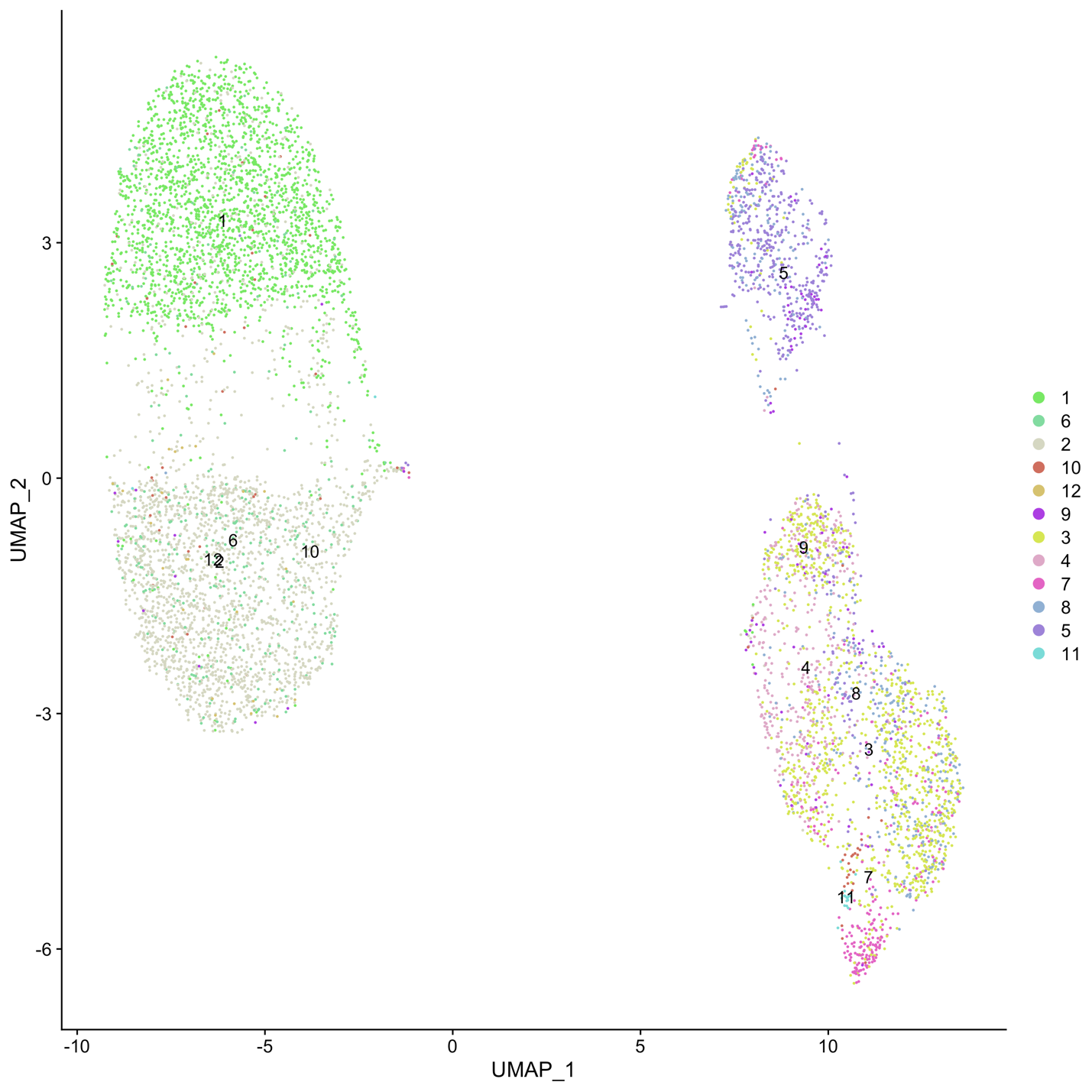

**Supplementary Figure S7:** UMAP visualizations of cells in ADT space colored by A) Antibody-cluster ground truth B) scConsensus C) Seurat and D) RCA in the antibody expression space for the MALT dataset

A) Ground Truth

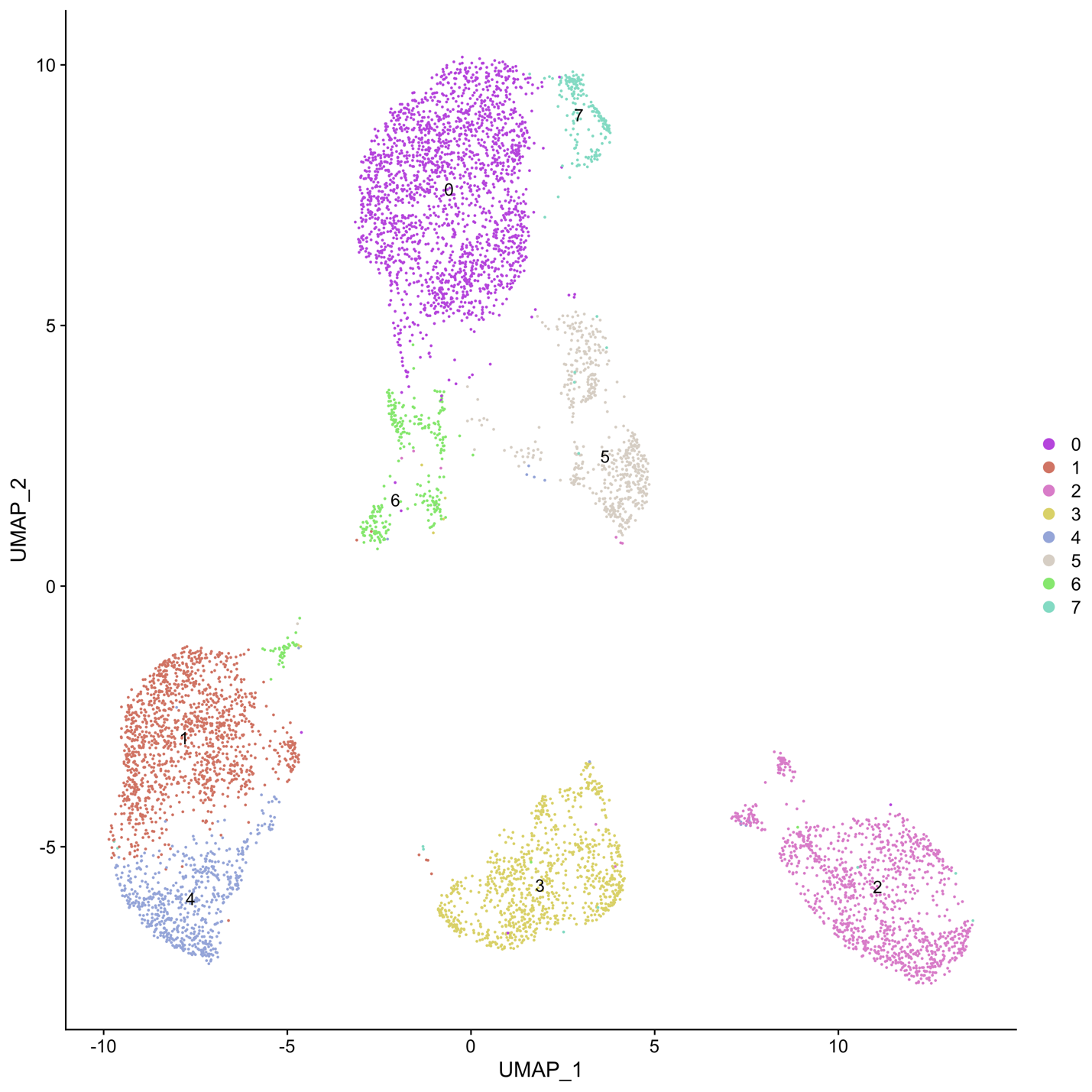

B) scConsensus

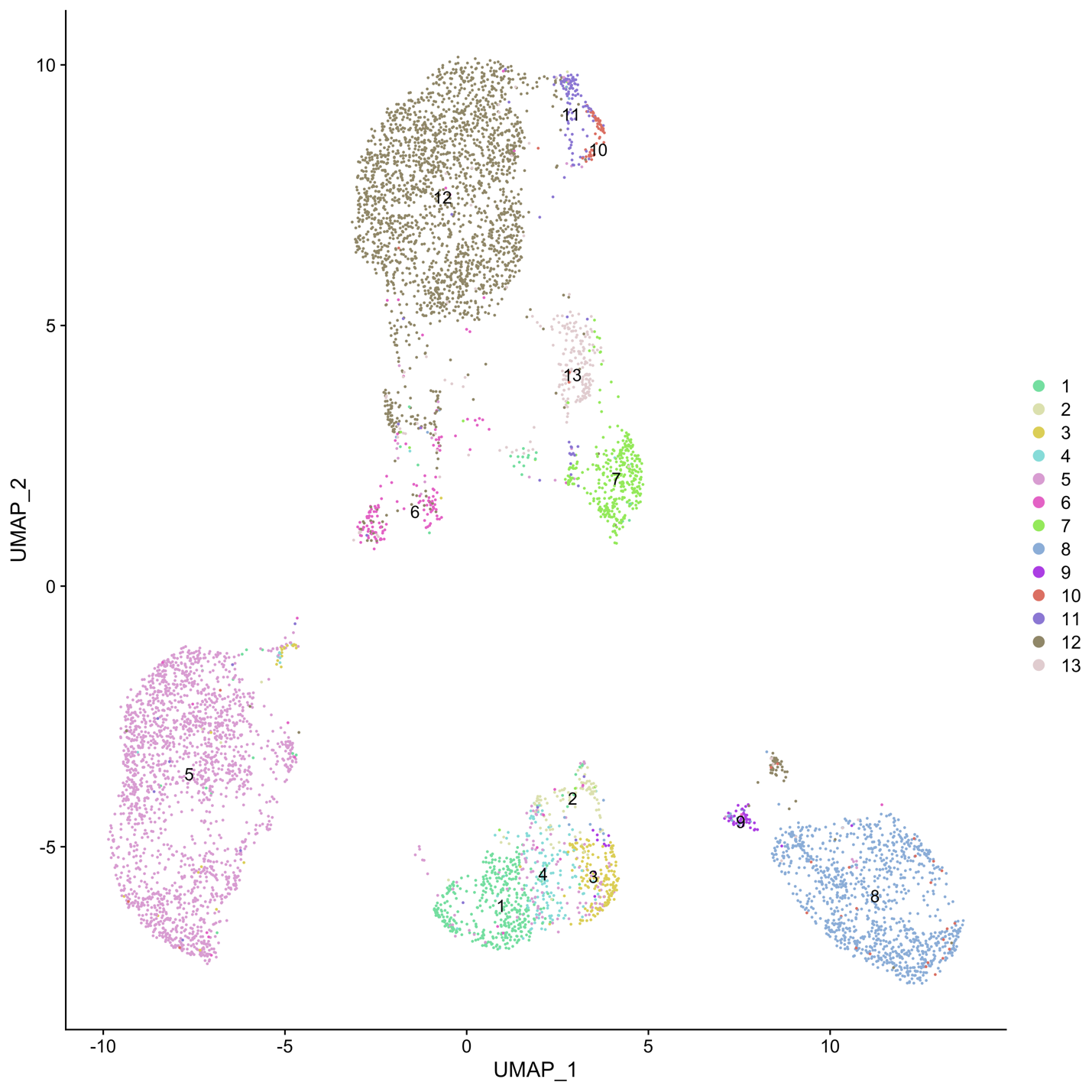

C) Seurat

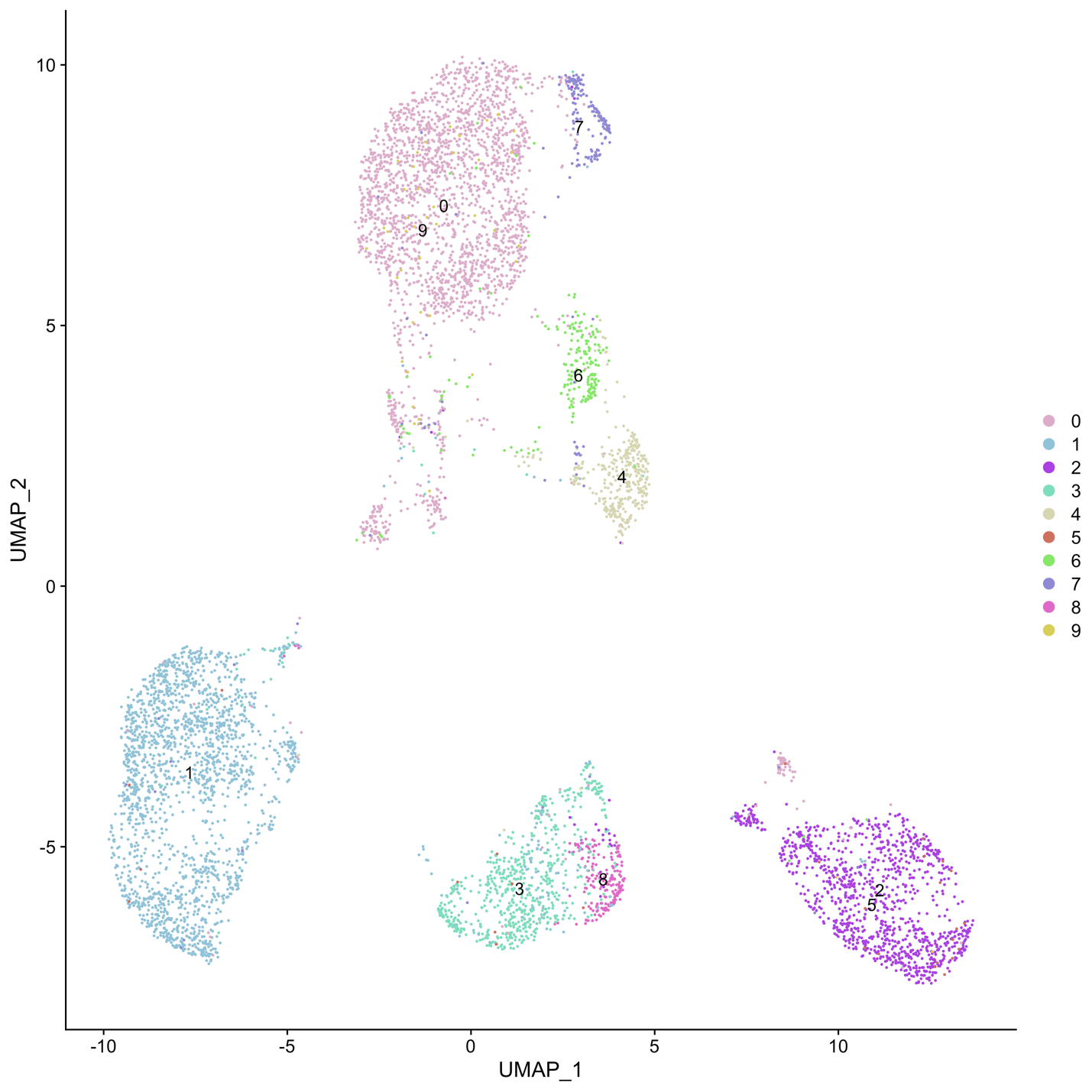

D) RCA

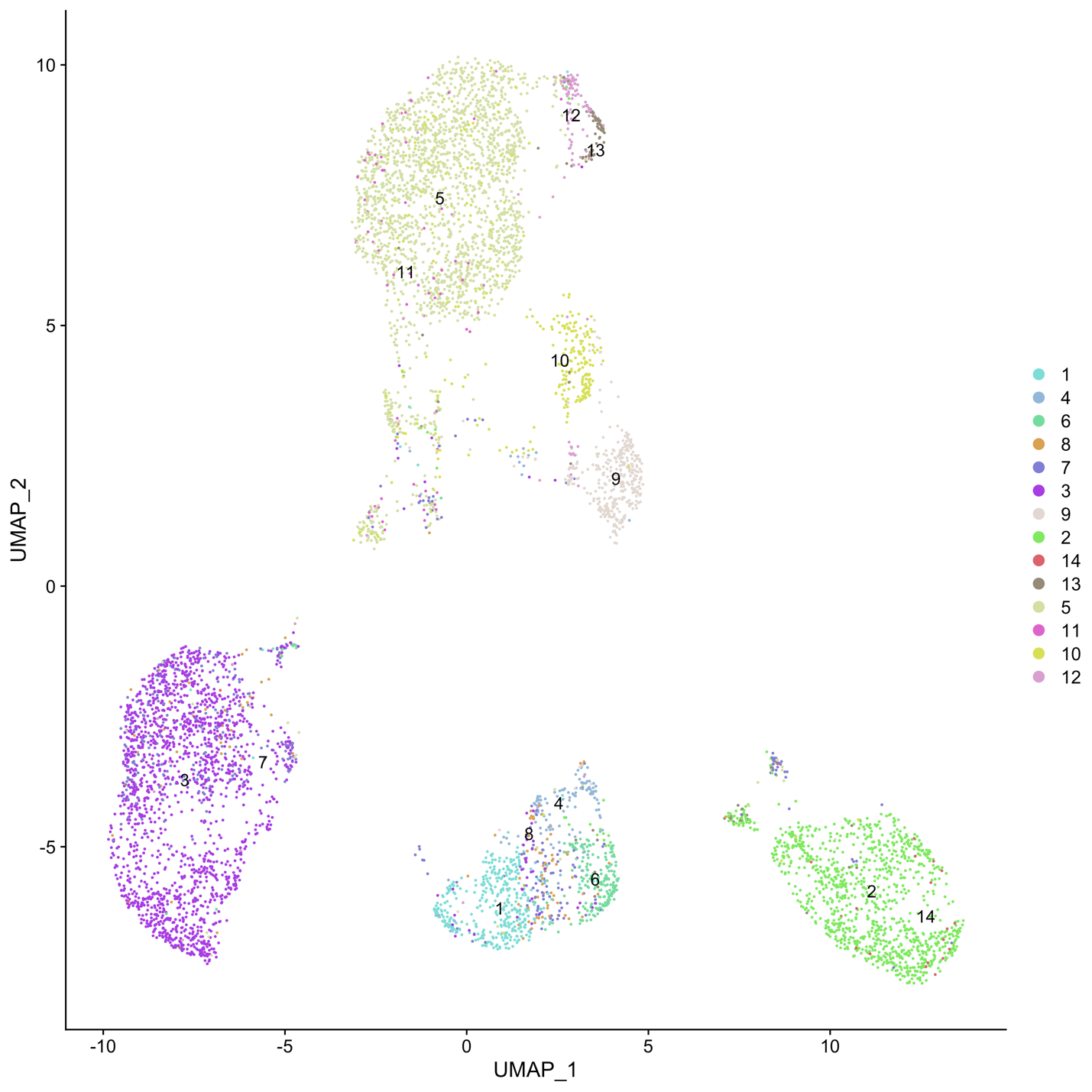

**Supplementary Figure S8:** UMAP visualizations of cells in ADT space colored by A) Antibody-cluster ground truth B) scConsensus C) Seurat and D) RCA in the antibody expression space for the PBMC-VDJ dataset

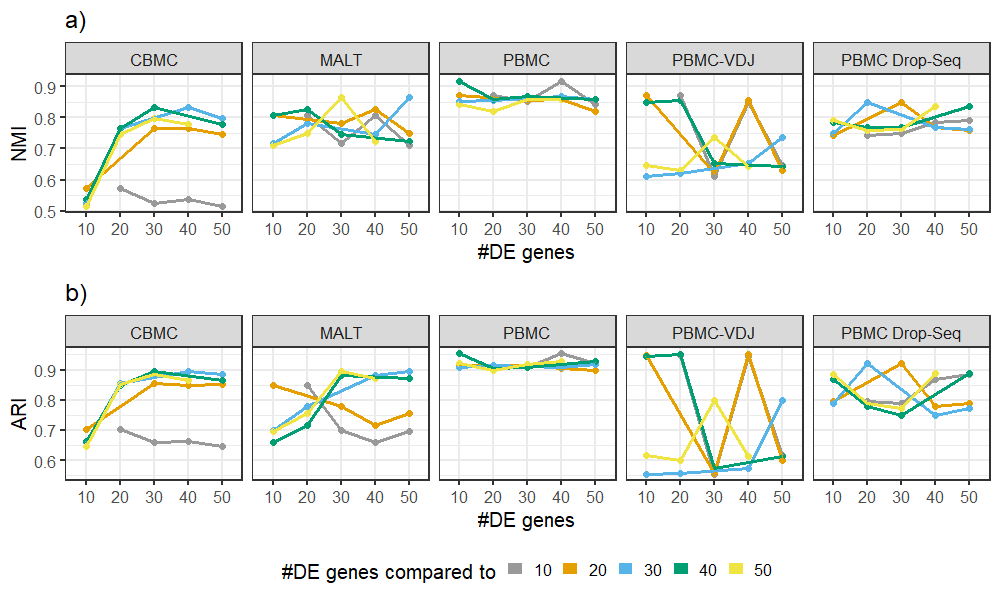

**Supplementary Figure S9:** Using (a) Normalized Mutual Information (NMI) and (b) Adjusted Rand Index (ARI), we measure the agreement between clustering results obtained using either 10, 20, 30, 40 or 50 DE (x-axis) genes compared to each other (color). Note that self-comparisons are omitted for clarity. For the *CBMC* data set, both scores remain constant from 30 DE genes onwards. For the *MALT* data set, we observe that considering 30 DE genes has a good agreement with a clustering based on 50 and 40 DE genes. Considering the *PBMC* dataset, the number of DE genes does not strongly influence the result at all. Interestingly, there is a lot of variation in the agreement between different clustering results in the *PBMC-VDJ* data. Roughly, clusterings using 10,20 and 40 DE genes agree with each other, while cluster results using 30 and 50 DE genes also agree well. For the PBMC Drop-Seq dataset, the variation is less pronounced, generally we observe that clusters with 10, 40 and 50 DE genes agree well with each other, although the highest scores are obtaine using 20 or 30 DE genes.

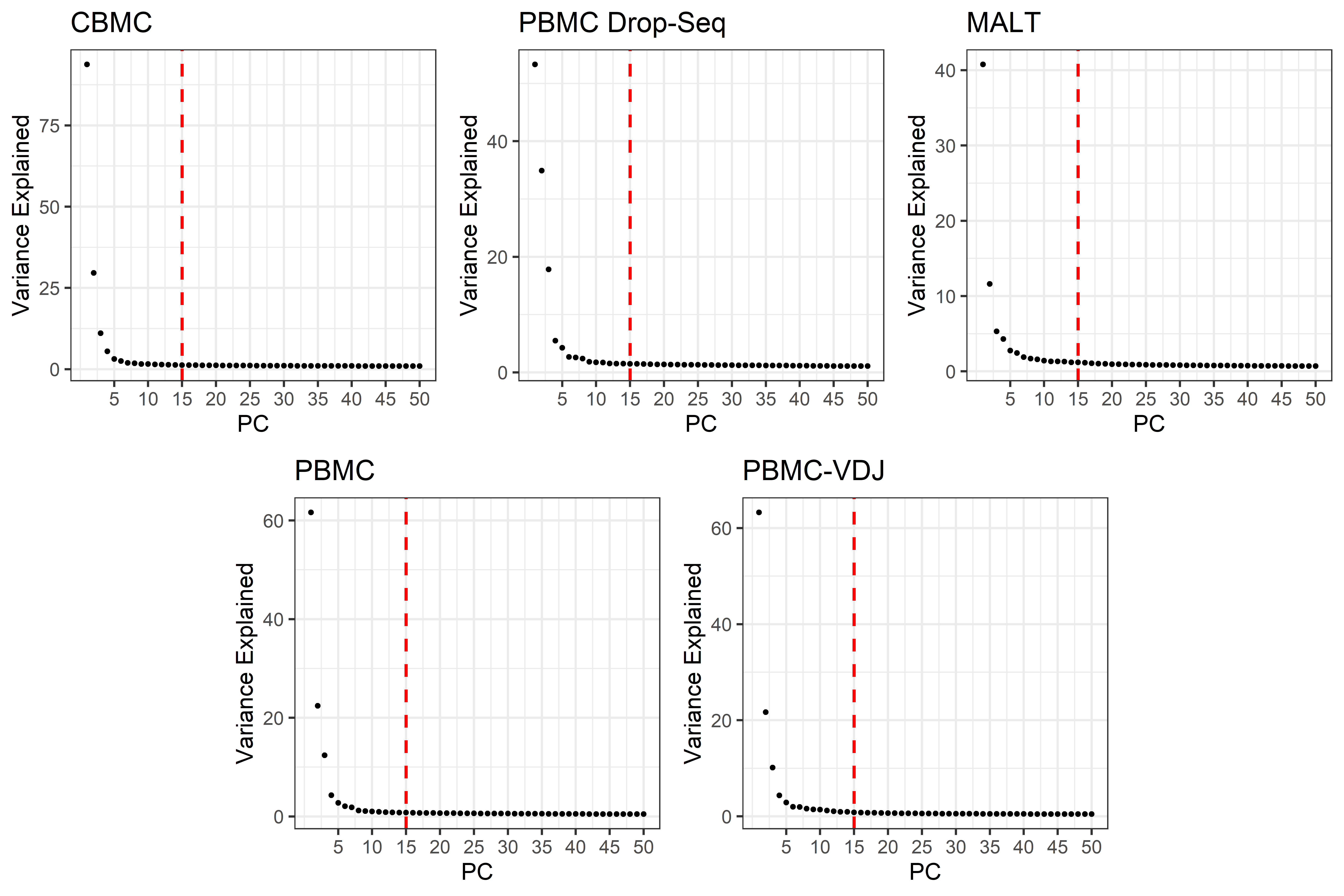

**Supplementary Figure S10:** Elbow plots showing the variance explained by principal components (PCs) for all datasets. Here, the red dashed line indicates the number of PCs that were selected for each dataset (15 PCs in each case). This figure shows that, for all datasets, the curve flattens out before 15 PCs, and there is little change in the variance explained by the subsequent PCs. Thus, we select 15 as a reasonable and consistent number of PCs for analysis of these datasets.

### Supplementary Tables

| **Dataset** | **Data Type** | **# genes** | **# cells** | **Reference** |
| --- | --- | --- | --- | --- |
| Cord Blood 10X  (CBMC) | 10X CITE-Seq UMI + ADT (Cord Blood Mononuclear Cells) | 4341 | 7817 | *Stoeckius, Marlon, et al. "Simultaneous epitope and transcriptome measurement in single cells." Nature methods 14.9 (2017): 865.* |
| Peripheral Blood Drop-Seq  (PBMC Drop-Seq) | 10X CITE-Seq UMI + ADT | 3976 | 7583 | *Stoeckius, Marlon, et al. "Simultaneous epitope and transcriptome measurement in single cells." Nature methods 14.9 (2017): 865.* |
| Mucosa-Associated Lymphoid Tissue 10X  (MALT) | 10X CITE-Seq UMI + ADT | 4713 | 8242 | [*https://support.10xgenomics.com/single-cell-gene-expression/datasets/3.0.0/malt_10k_protein_v3*](https://support.10xgenomics.com/single-cell-gene-expression/datasets/3.0.0/malt_10k_protein_v3) |
| Peripheral Blood 10X  (PBMC) | 10X CITE-Seq UMI + ADT | 5924 | 7750 | [*https://support.10xgenomics.com/single-cell-gene-expression/datasets/3.0.0/pbmc_10k_protein_v3*](https://support.10xgenomics.com/single-cell-gene-expression/datasets/3.0.0/pbmc_10k_protein_v3) |
| Peripheral Blood 10X-VDJ  (PBMC-VDJ) | 10X CITE-Seq UMI + ADT  (5’ Gene Expression) | 5185 | 7627 | [*https://support.10xgenomics.com/single-cell-vdj/datasets/3.0.0/vdj_v1_hs_pbmc2_5gex_protein*](https://support.10xgenomics.com/single-cell-vdj/datasets/3.0.0/vdj_v1_hs_pbmc2_5gex_protein) |
| PBMCs FACS | 10X GemCode Single-Cell 3’ Library | 10098 | 25389 | Zheng et al., “Massively parallel digital transcriptional profiling of single cells”, Nature Communications 14049 (2017) |

**Supplementary Table S1:** Details on published scRNA-seq data sets used in the study.

| **Dataset** | **Minimum NODG** | **Maximum NODG** | **Maximum % mito** | **Minimum number of cells a gene must be expressed in** |
| --- | --- | --- | --- | --- |
| Cord Blood 10X | 500 | 2000 | 8 | 100 |
| Peripheral Blood Drop-Seq | 500 | 2000 | 8 | 100 |
| Lymphoid Tissue 10X | 500 | 3000 | 30 | 100 |
| Peripheral Blood 10X | 500 | 3000 | 15 | 100 |
| Peripheral Blood 10X-VDJ | 500 | 4000 | 15 | 100 |
| PBMCs FACS | 300 | 2000 | 8 | 100 |

**Supplementary Table S2:** QC Metrics for data pre-processing. NODG: Number of Detected Genes

| **Variance threshold** | **CBMC** | **PBMC Drop-Seq** | **MALT** | **PBMC** | **PBMC-VDJ** |
| --- | --- | --- | --- | --- | --- |
| 0.5 | 1239 | 1457 | 826 | 866 | *620* |
| 1 | 348 | 471 | 82 | 75 | *60* |

**Supplementary Table S3:** Number of genes used for bootstrapping depending on the variance threshold.

### Supplementary Notes

#### Supplementary Note 1 – Generation of an Immune Reference Panel

The immune reference panel was generated by downloading the bulk RNA sequencing data from GEO Accession Viewer: GSE107011 (Monaco et al. 2019). Cells derived from PBMCs and CD4_TE s were removed from all donors. In addition, VD2 positive cell from DZQV donor was removed. Quantile normalization was done using the preprocessCore R package. Genes with TPM greater and equal to 10 in at least 3 biological replicates were retained and log normalization with base 10 was performed. Batch correction using COMBAT (SVA R package) was done to remove the Batch effects. Gene expression for each gene was averaged across donors prior to feature gene identification. Lastly, feature genes were selected based on cell type specific expression in the dataset. A gene is included in the feature gene set if the fold change of its expression relative to other cell types is greater or equal to five.

options(stringsAsFactors=FALSE)

args=(commandArgs(TRUE))

optional_args <- c()

for(i in 1:length(args)) {

eval(parse(text=args[[i]]))

arg_str <- gsub("^(\\S+)=\\S+$", "\\1", args[[i]])

if(

arg_str == "with_replicates" ||

arg_str == "is_bulk" ||

arg_str == "detected_gene_cutoff" ||

arg_str == "hq_genes_cutoff" ||

arg_str == "log_transform" ||

arg_str == "log_pseudo" ||

arg_str == "log_base" ||

arg_str == "log_pseudo_replace" ||

arg_str == "normalization_method" ||

arg_str == "fs_fc_cutoff" ||

arg_str == "fs_quantile_probe"

) {

optional_args <- c(optional_args, args[[i]])

}

}

\# args default value

with_replicates <- FALSE

is_bulk <- TRUE

detected_gene_cutoff <- 0

hq_genes_cutoff <- 2

log_transform <- TRUE

log_pseudo <- 1

log_base <- 10

log_pseudo_replace <- FALSE

normalization_method <- "quantile"

fs_fc_cutoff <- 10

fs_quantile_probe <- 0.5

\# parsing optional args

optional_args_str <- paste0(optional_args, collapse=";")

eval(parse(text=optional_args_str))

\# read TPM data

exprs <- readRDS(exprs_file)

######################################################

########## plot violin plot of TPM	#################

######################################################

beforenorm_violin_in <- exprs

if(log_transform) {

if(log_pseudo_replace) {

beforenorm_violin_in[beforenorm_violin_in < log_pseudo] <- log_pseudo

beforenorm_violin_in <- log(beforenorm_violin_in, base=log_base)

} else {

beforenorm_violin_in <- log(beforenorm_violin_in + log_pseudo, base=log_base)

}

}

if(!require("reshape2")) {

install.packages("reshape2", repos="http://cran.stat.nus.edu.sg")

require(reshape2)

}

beforenorm_violin_ggplot_in <- melt(as.data.frame(beforenorm_violin_in))

colnames(beforenorm_violin_ggplot_in) <- c("celltype", "value")

if(!require("ggplot2")) {

install.packages("ggplot2", repos="http://cran.stat.nus.edu.sg")

require("ggplot2")

}

pdf(paste0(output_dir, "/violin_beforenorm.pdf"), width=14, height=7)

ggplot(beforenorm_violin_ggplot_in, aes(x=celltype, y=value)) + geom_violin() + geom_boxplot(width=.1, fill="red") + theme_bw() + theme(axis.text.x = element_text(angle = 90, hjust = 1)) + labs(y="log(expression)")

dev.off()

#######################################################

##########	quantile norm	###########################

#######################################################

if (normalization_method=="quantile") {

\# quantile normalization

if(!require("preprocessCore")) {

source("http://bioconductor.org/biocLite.R")

biocLite("preprocessCore")

require(preprocessCore)

}

quantile_out <- normalize.quantiles.robust(as.matrix(exprs), use.median=TRUE)

rownames(quantile_out) <- rownames(exprs)

colnames(quantile_out) <- colnames(exprs)

} else if (normalization_method=="iQ") {

quantile_out <- iQ_columnsorted_mediantarget_correctassignback(exprs)

} else {

cat("Unknown normalization method, use normalize.quantiles.robust(, use.median=TRUE) from preprocessCore ...\n")

\# quantile normalization

if(!require("preprocessCore")) {

source("http://bioconductor.org/biocLite.R")

biocLite("preprocessCore")

require(preprocessCore)

}

quantile_out <- normalize.quantiles.robust(as.matrix(exprs), use.median=TRUE)

rownames(quantile_out) <- rownames(exprs)

colnames(quantile_out) <- colnames(exprs)

}

#######################################################

##########	violin plot after quantile	###############

#######################################################

afternorm_violin_in <- quantile_out

if(log_transform) {

if(log_pseudo_replace) {

afternorm_violin_in[afternorm_violin_in < log_pseudo] <- log_pseudo

afternorm_violin_in <- log(afternorm_violin_in, base=log_base)

} else {

afternorm_violin_in <- log(afternorm_violin_in + log_pseudo, base=log_base)

}

}

afternorm_violin_ggplot_in <- melt(as.data.frame(afternorm_violin_in))

colnames(afternorm_violin_ggplot_in) <- c("celltype", "value")

pdf(paste0(output_dir, "/violin_afternorm.pdf"), width=14, height=7)

ggplot(afternorm_violin_ggplot_in, aes(x=celltype, y=value)) + geom_violin() + geom_boxplot(width=.1, fill="red") + theme_bw() + theme(axis.text.x = element_text(angle = 90, hjust = 1)) + labs(y="log(expression)")

dev.off()

######################################################

######### plot histogram of median	##########

######### expression of gene across samples ##########

######################################################

median_of_genes<- apply(quantile_out,1,function(x) median(x[x>0]))

\# plot the histogram to check the distribution of genes across

pdf(paste0(output_dir,"/histogram_medianofgeneexpressionacross_samples.pdf"), width=14, height=7)

hist(median_of_genes, breaks=10000, xlim=c(0,100), col="pink")

dev.off()

######################################################

########### find expressed genes	##############

######################################################

print(sample_info_file)

sample_info <- read.table(sample_info_file, sep="\t", header=TRUE, as.is=TRUE, check.names=FALSE)

rownames(sample_info) <- sample_info$sample_id

\# make sure that the gene expressed above TPM > detected_gene_cutoff in any cell type in at least 3 biological replicates

check_reps_gene_exprs<-t(apply(as.matrix(quantile_out), 1,function(v){aggregate(v, by=list(celltype=sample_info[colnames(quantile_out), "celltype"]), function(x){sum(x>=detected_gene_cutoff) >= hq_genes_cutoff})$x}))

colnames(check_reps_gene_exprs) <- aggregate(as.matrix(quantile_out)[1, ], by=list(celltype=sample_info[colnames(quantile_out), "celltype"]), mean)$celltype

\# get number of cell types

cell_types <- list(celltype=sample_info[colnames(quantile_out), "celltype"])$celltype

number_of_cell_types <- length(unique(cell_types))

print(number_of_cell_types)

\# filter those transcripts/genes that fail the above criteria

\# rows that have all FALSE across all cell types

\# expressed_genes are genes that do not have FALSE for all 28 cell types

matrix_expressed <- quantile_out[which(apply(check_reps_gene_exprs, 1,function(x)sum(x==FALSE))!= number_of_cell_types),]

#######################################################

##########	log transformation	#######################

#######################################################

if(log_transform) {

if(log_pseudo_replace) {

matrix_expressed[matrix_expressed < log_pseudo] <- log_pseudo

quantile_log <- log(matrix_expressed, base=log_base)

} else {

quantile_log <- log(matrix_expressed + log_pseudo, base=log_base)

}

}

saveRDS(quantile_log, file=paste0(output_dir, "/After_quantile_normalization.rds"))

#######################################################

##########	COMBAT to correct for batch effects #############

########################################################

library(sva)

batch <- factor(sample_info$sample_type)

celltype<-factor(sample_info$celltype)

modcombat <- model.matrix(~1+celltype, data=sample_info)

combat_edata <- ComBat(dat=quantile_log,batch=batch, mod=modcombat, par.prior = TRUE,prior.plots=FALSE)

saveRDS(combat_edata, file=paste0(output_dir, "/After_combat_expression_data.rds"))

\# plot a dendrogram to see the clsutering of samples after combat

pdf(paste0(output_dir, "/all_expressed_genes_after_combat_before_choosing_featuregenes_hclust.pdf"), height=7, width=14)

plot(hclust(as.dist(1 - cor(combat_edata)), method="average"), xlab="hclust(as.dist(1 - cor(*)), method=\"average\")", main=paste0("Number of genes: ", nrow(combat_edata)))

dev.off()

###################################################################################

##########	merge replicates on feature genes #####################################

###################################################################################

if(with_replicates) {

sample_info <- read.table(sample_info_file, sep="\t", header=TRUE, as.is=TRUE, check.names=FALSE)

rownames(sample_info) <- sample_info$sample_id

expr_gene_repmerge <- t(apply(as.matrix(combat_edata), 1,

function(v){

aggregate(v, by=list(celltype=sample_info[colnames(combat_edata), "celltype"]), mean)$x

}

))

colnames(expr_gene_repmerge) <- aggregate(as.matrix(combat_edata)[1, ], by=list(celltype=sample_info[colnames(combat_edata), "celltype"]), mean)$celltype

saveRDS(expr_gene_repmerge, file=paste0(output_dir, "/expr_gene_repmerge_average.rds"))

}

###################################################################

##########	DE genes selection for ref panel ##################

###################################################################

logFC_thr <- log(fs_fc_cutoff, base=log_base)

tmp_binary <- t(apply(expr_gene_repmerge, 1, function(v){(v-quantile(v, probs=fs_quantile_probe)) > logFC_thr})) + 0

foldchange <- t(apply(expr_gene_repmerge,1,function(v){(v-quantile(v, probs=fs_quantile_probe))}))

saveRDS(foldchange,file=paste0(output_dir, "/fold_change_aftermerge.rds"))

\# plot the violin plot for fold change

foldchange_violin_ggplot_in <- melt(as.data.frame(foldchange))

colnames(foldchange_violin_ggplot_in) <- c("celltype", "value")

pdf(paste0(output_dir, "/violin_foldchange_aftermerge.pdf"), width=14, height=7)

ggplot(foldchange_violin_ggplot_in, aes(x=celltype, y=value)) + geom_violin() + geom_boxplot(width=.1, fill="red") + theme_bw() + theme(axis.text.x = element_text(angle = 90, hjust = 1)) + labs(y="foldchange")

dev.off()

featuregene_binary <- tmp_binary[apply(tmp_binary, 1, function(v){sum(v) > 0}), ]

expr_featuregene <- expr_gene_repmerge[rownames(featuregene_binary), ]

saveRDS(expr_featuregene, file=paste0(output_dir, "/expr_featuregene_aftermerge.RDS"))

saveRDS(featuregene_binary, file=paste0(output_dir, "/featuregene_binary_aftermerge.RDS"))

pdf(paste0(output_dir, "/expr_featuregene_aftermerge_hclust.pdf"), height=7, width=14)

plot(hclust(as.dist(1 - cor(expr_featuregene)), method="average"), xlab="hclust(as.dist(1 - cor(*)), method=\"average\")", main=paste0("Number of feature genes: ", nrow(expr_featuregene)))

dev.off()

library(gplots)

pdf(paste0(output_dir, "/expr_featuregene_aftermerge_selfprojection.pdf"), height=20, width=20)

color_scheme <- colorRampPalette(c("#00007F", "blue", "#007FFF", "cyan", "#7FFF7F", "yellow", "#FF7F00", "red", "#7F0000"))(100)

heatmap_in <- cor(expr_featuregene)

if(with_replicates) {

if (!require("WGCNA")) {

source("http://bioconductor.org/biocLite.R")

biocLite(c("AnnotationDbi", "impute", "GO.db", "preprocessCore"), suppressUpdates=TRUE)

install.packages("WGCNA", repos="http://cran.stat.nus.edu.sg")

require(WGCNA)

}

sample_info <- read.table(sample_info_file, sep="\t", header=TRUE, as.is=TRUE, check.names=FALSE)

rownames(sample_info) <- sample_info$sample_id

heatmap.2(heatmap_in,col=color_scheme,

Colv=as.dendrogram(hclust(as.dist(1 - cor(heatmap_in)), method="average")),

Rowv=as.dendrogram(hclust(as.dist(1 - cor(heatmap_in)), method="average")),

RowSideColors=labels2colors(sample_info[colnames(heatmap_in), "celltype"]),

scale="none", margins=c(5,20),

trace="none",

key = TRUE,

keysize = 0.5,

cexCol = 1,cexRow =1,

labCol = "",

main = paste0("Number of feature genes: ", nrow(expr_featuregene))

)

dev.off()

} else {

heatmap.2(heatmap_in,col=color_scheme,

Colv=as.dendrogram(hclust(as.dist(1 - cor(heatmap_in)), method="average")),

Rowv=as.dendrogram(hclust(as.dist(1 - cor(heatmap_in)), method="average")),

scale="none", margins=c(5,20),

trace="none",

key = TRUE,

keysize = 0.5,

cexCol = 1,cexRow =1,

labCol = "",

main = paste0("Number of feature genes: ", nrow(expr_featuregene))

)

dev.off()

}

library(gplots)

pdf(paste0(output_dir, "/featuregene_binary_aftermerge.pdf"), height=20, width=20)

heatmap.2(featuregene_binary,

Colv=as.dendrogram(hclust(as.dist(1 - cor(expr_featuregene)), method="average")),

scale="none", margins=c(20,20),

trace="none",

dendrogram="column",

key = TRUE,

keysize = 0.5,

cexCol = 1.5,cexRow = 0.1,

lhei=c(0.2,2)

)

dev.off()

#### **Supplementary Note 2** – Preprocessing and scConsensus Execution for CITE-Seq data

Function to compute DE gene refinement

#'

#'@author: ranjanb

#'

#'@description: ComputeDatasetGroundTruth() takes in a dataMatrix and clusterLabels and outputs the DE gene

#'heatmap along with a data object containing the q-values, log2-fold changes, DE genes, union of DE genes, DE gene

#'counts and the up and down regulated DE genes for each pairwise comparison.

#'

#'@inputs:

#'matrix dataMatrix: genes (rows) x cells (columns) log10 transformed gene expression matrix

#'named character vector clusterLabels: vector of cluster labels, named using corresponding cell IDs (ordered as columns of dataMatrix)

#'character method: defines the method to be used to compute DE genes, can take the value of "Wilcoxon", "LimmaVoom" or "edgeR". default = "Wilcoxon".

#'numeric meanScalingFactor: scale of the mean gene expression across the gene expression matrix to set a minimum threshold of average cluster expression for a gene to be considered a DE gene

#'numeric qValThrs: maximum q-value threshold for Wilcoxon Rank Sum Test

#'numeric fcThrs: minimum fold-change threshold for DE gene criterion

#'vector deepSplitValues: vector of WGCNA tree cutting deepsplit parameters

#'#'character minClusterSizeType: specifies the type of minimum cluster factor the user would like to specify: "scaled" for a fraction of the number of cells OR "absolute" for the absolute value of minimum number of cells. default = "scaled"

#'integer minClusterFactor: factor to determine minimum number of cells in each cluster.

#'character dataType: specifies if the type of data is 10x or not, and hence if it requires additional normalization or not

#'character folderOut: specifies the output directory of the results

#'

#'@output:

#'returnObj = list(

#'list qValueList: list of q-values for each pairwise cluster comparison

#'list log2FCList: list of log2-fold-changes for each pairwise cluster comparison

#'list deGeneList: list of de genes for each pairwise cluster comparison

#'character vector deGeneUnion: union of all DE genes

#'numeric matrix deCountMatrix: matrix with number of DE genes for each pairwise cluster comparison

#'list deGeneRegulationList: list of up and down regulated DE genes

#'list upregulatedDEGeneList: list of cluster-specific upregulated DE genes

#'list downregulatedDEGeneList: list of cluster-specific downregulated DE genes

#'list cellTree: dendrogram of cells clustered in deGeneUnion space

#'list dynamicColors: list of cluster labels based on clustering in deGeneUnion space over all deepSplitValues

#')

#'

\# source("R:/CSB/CSB6/Private/Bobby/rna-seq-pipelines/cellTypeDEPlot.R")

\# source("R:/CSB/CSB6/Private/Bobby/rna-seq-pipelines/plotVolcano.R")

ComputeDatasetGroundTruth <- function(dataMatrix,

clusterLabels,

method = "Wilcoxon",

meanScalingFactor = 5,

qValThrs,

fcThrs,

deepSplitValues = 1:4,

minClusterSizeType = "scaled",

minClusterFactor = 100,

dataType = "10x",

folderOut = ".") {

\### Check if package dependencies are available; if not, download from CRAN and require those packages

\# Seurat

if (!require(Seurat))

install.packages("Seurat", repos = "http://cran.us.r-project.org")

require(Seurat)

\# flashClust

if (!require(flashClust))

install.packages("flashClust", repos = "http://cran.us.r-project.org")

require(flashClust)

\# calibrate

if (!require(calibrate))

install.packages("calibrate", repos = "http://cran.us.r-project.org")

require(calibrate)

\# WGCNA

if (!require(WGCNA)) {

source("http://bioconductor.org/biocLite.R")

biocLite(c("impute", "GO.db", "preprocessCore"))

install.packages("WGCNA")

}

require(WGCNA)

\# Limma

if (!require(limma)) {

if (!requireNamespace("BiocManager", quietly = TRUE))

install.packages("BiocManager")

BiocManager::install("limma", version = "3.8")

}

require(limma)

\# edgeR

if (!require(edgeR)) {

if (!requireNamespace("BiocManager", quietly = TRUE))

install.packages("BiocManager")

BiocManager::install("edgeR", version = "3.8")

}

require(edgeR)

\### Initialise data input

dataIn <- as.matrix(dataMatrix)

\### Initialize minimum cluster size

if (minClusterSizeType == "scaled") {

minClusterSize = ceiling(ncol(dataIn) / minClusterFactor)

} else if (minClusterSizeType == "absolute") {

minClusterSize = minClusterFactor

} else {

minClusterSize = 10

}

\### 10x data needs to be log normalized by cell using Seurat's LogNormalize method to reduce ties in data before calling DE genes

if (dataType == "10x") {

\### Log Normalize using Seurat

seuratObj <- CreateSeuratObject(counts = dataIn)

seuratObj <-

NormalizeData(

object = seuratObj,

normalization.method = "LogNormalize",

scale.factor = 10000

)

\### Return normalized data to data matrix

dataIn <- as.matrix(seuratObj@assays$RNA@data)

\### Convert log base from e to 2

dataIn <- dataIn / log(2)

} else {

\### Convert log base from e to 2

dataIn <- dataIn / log(2)

}

\### Initialize mean expression threshold

meanExprsThrs = meanScalingFactor * mean(2^dataIn-1)

\### Use only clusters with number of cells > minimum cluster size for DE gene calling

which(table(clusterLabels) > minClusterSize)

colorCounts <-

table(clusterLabels)

\### Extract unique cluster labels

uniqueClusters <-

names(colorCounts[colorCounts > minClusterSize])

\# ignore the grey cluster because it represents unclustered cells

uniqueClusters <-

uniqueClusters[!grepl("grey", uniqueClusters)]

\### If the cluster label vector is unnamed, name it in the order of the data matrix columns

if (is.null(names(clusterLabels))) {

names(clusterLabels) <- colnames(dataIn)

}

\### Initialize nested lists to store q values, log-normalized fold change values and de genes

qValueList <-

rep(list(list()), length(uniqueClusters))

log2FCList <-

rep(list(list()), length(uniqueClusters))

deGeneList <- rep(list(list()), length(uniqueClusters))

\### Initialise number of comparisons as n(n-1)/2

numComparisons <-

(length(uniqueClusters) * (length(uniqueClusters) - 1)) / 2

\### Conduct pairwise cluster comparison to obtain q-values, log-normalized fold changes and

\### DE genes for each comparison

for (i in 1:(length(uniqueClusters) - 1)) {

for (j in (i + 1):length(uniqueClusters)) {

\### Get the cell names cell data for cluster i

cellNamesi <-

names(clusterLabels)[which(clusterLabels == uniqueClusters[i])]

cellDatai <- dataIn[, cellNamesi]

\### Get the cell names cell data for cluster j

cellNamesj <-

names(clusterLabels)[which(clusterLabels == uniqueClusters[j])]

cellDataj <- dataIn[, cellNamesj]

deCellData <- cbind(cellDatai, cellDataj)

\### Declare variables to store p-values, q-values and log2 fold change values for each pairwise gene comparison

pval <- NA

qval <- NA

log2fc <- NA

if (method == "Wilcoxon") {

\### For each gene, conduct Wilcoxon Rank Sum test to obtain q-value and log2-normalized fold change

for (k in 1:nrow(cellDatai)) {

\# extract data of gene k for cluster i

geneDatai <- cellDatai[k, ]

\# extract data of gene k for cluster j

geneDataj <- cellDataj[k, ]

\# run wilcoxon test

wcTestOut <-

wilcox.test(geneDatai, geneDataj)

\# initialize q value and fold change for each gene

pval[k] <- wcTestOut$p.value

log2fc[k] <-

mean(geneDatai) - mean(geneDataj)

}

\# check if gene satisfies mean expression threshold

meanExprsLogicalVector <-

apply(deCellData, 1, function(row) {

mean(row[cellNamesi]) > log2(meanExprsThrs) |

mean(row[cellNamesj]) > log2(meanExprsThrs)

})

\# adjust p-value using Benjamini-Hochberg adjustment

qval <-

p.adjust(

p = pval,

method = "BH",

n = nrow(cellDatai)

)

} else if (method == "LimmaVoom") {

\# initialise vector of 1s to denote cells of cluster i and -1s to denote cells of cluster j

limmaGroupi <-

rep(uniqueClusters[i], ncol(cellDatai))

limmaGroupj <-

rep(uniqueClusters[j], ncol(cellDataj))

limmaGroup <-

c(limmaGroupi, limmaGroupj)

dge <-

DGEList(deCellData, group = limmaGroup)

dge <- calcNormFactors(dge)

design <- model.matrix(~ limmaGroup)

vm <- voom(dge, design = design, plot = TRUE)

fit <- lmFit(vm, design = design)

fit <- eBayes(fit)

tt <-

topTable(fit = fit,

n = Inf,

adjust.method = "BH")

\# # compute p-value using limma-voom fitting & eBayes method

\# pval <- apply(deCellData, 1, function(v) {

\#	fit <- lmFit(v, limmaGroup)

\#	return(eBayes(fit, trend = TRUE)$p.value)

\# })

\# adjust p-value using Benjamini-Hochberg adjustment

\# qval <-

\#	p.adjust(

\#	p = pval,

\#	method = "BH",

\#	n = numComparisons * nrow(cellDatai)

\#	)

qval <- tt$adj.P.Val

\# compute q-value in log space by finding difference of means

log2fc <-

apply(deCellData, 1, function(row) {

mean(row[cellNamesi]) - mean(row[cellNamesj])

})

\# check if gene satisfies mean expression threshold

meanExprsLogicalVector <-

apply(deCellData, 1, function(row) {

mean(row[cellNamesi]) > log2(meanExprsThrs) |

mean(row[cellNamesj]) > log2(meanExprsThrs)

})

} else if (method == "edgeR") {

\# Create DGE object

dgeObj <- DGEList(counts=deCellData, group = c(rep(1, ncol(cellDatai)), rep(-1, ncol(cellDataj))))

\# Estimate common and tag-wise dispersion for the pair of clusters

dgeObj <- estimateCommonDisp(dgeObj)

dgeObj <- estimateTagwiseDisp(dgeObj)

\# Calculate norm factors for data

dgeObj <- calcNormFactors(dgeObj)

\# Perform exact test

etObj <- exactTest(object = dgeObj)

\# Save p-value results

pval <- etObj$table$PValue

\# adjust p-value using Benjamini-Hochberg adjustment

qval <-

p.adjust(

p = pval,

method = "BH",

n = nrow(cellDatai)

)

log2fc <- etObj$table$logFC

\# check if gene satisfies mean expression threshold

meanExprsLogicalVector <-

apply(deCellData, 1, function(row) {

mean(row[cellNamesi]) > log2(meanExprsThrs) |

mean(row[cellNamesj]) > log2(meanExprsThrs)

})

} else {

\### exit from function if no method is chosen

print("Incorrect method chosen.")

return(NULL)

}

\### determine if a gene is a DE gene based on thresholds

deGeneLogicalVector <-

qval < qValThrs &

abs(log2fc) > log2(fcThrs)

\# filter de genes by mean expression

deGeneLogicalVector <- deGeneLogicalVector & meanExprsLogicalVector

print(paste0(

uniqueClusters[i],

", ",

uniqueClusters[j],

" DE genes: ",

sum(deGeneLogicalVector)

))

\### store q-values, log-normalized fold change and DE genes

qValueList[[i]][[j]] <- qval

log2FCList[[i]][[j]] <- log2fc

deGeneList[[i]][[j]] <-

rownames(cellDatai)[deGeneLogicalVector]

if (sum(deGeneLogicalVector) <= 1)

next

de_genes <- deGeneList[[i]][[j]]

}

}

\### Name the q-value, log2 fold change and DE gene lists by cluster names

names(qValueList) <- uniqueClusters

names(log2FCList) <- uniqueClusters

names(deGeneList) <- uniqueClusters

\### Obtain union of all de genes

\# initialize empty DE gene union vector

deGeneUnion <- c()

\# For each pair of clusters

for (i in 1:(length(uniqueClusters) - 1)) {

for (j in (i + 1):length(uniqueClusters)) {

\# obtain the union of the current DE gene union so far with the next DE gene list

deGeneUnion <-

union(deGeneUnion, deGeneList[[i]][[j]])

\# Rank the DE genes for each comparison by fold change and take the top 20

\# de_log2fc <- log2FCList[[i]][[j]][which(rownames(dataIn) %in% deGeneList[[i]][[j]])]

\# names(de_log2fc) <- rownames(dataIn)[which(rownames(dataIn) %in% deGeneList[[i]][[j]])]

\#

\# sorted_de_log2fc <- sort(x = abs(de_log2fc), decreasing = T)

\#

\# if(length(sorted_de_log2fc) > 20) {

\#	de_genes <- names(sorted_de_log2fc)[1:20]

\# } else {

\#	de_genes <- names(sorted_de_log2fc)

\# }

\#

\# deGeneUnion <-

\#	union(deGeneUnion, de_genes)

}

}

print(str(deGeneUnion))

\### Make DE gene matrix symmetric, as it is currrently an upper triangular matrix

\# First cover every element barring the last row and last column

for (i in 1:(length(uniqueClusters) - 1)) {

for (j in 1:(length(uniqueClusters) - 1)) {

\# clusters do not have DE genes with themselves, so skip to next iteration

if (i == j) {

next

}

\# if an (i,j) pair is empty, fill it with the corresponding (j,i) DE genes

if (is.null(deGeneList[[i]][[j]])) {

deGeneList[[i]][[j]] <- deGeneList[[j]][[i]]

}

}

\# Name the DE gene list with cluster names

names(deGeneList[[i]]) <- uniqueClusters

}

\# For the last row and last column

for (i in 1:(length(uniqueClusters) - 1)) {

\# set the DE genes of (last_row, i) as DE genes (i, last_row)

deGeneList[[length(uniqueClusters)]][[i]] <-

deGeneList[[i]][[length(uniqueClusters)]]

\# name the DE gene list with cluster names

names(deGeneList[[length(uniqueClusters)]])[i] <-

uniqueClusters[i]

}

\## Make the log2 fold change matrix symmetric, as it is currrently an upper triangular matrix

\# First cover every element barring the last row and last column

for (i in 1:(length(uniqueClusters) - 1)) {

for (j in 1:(length(uniqueClusters) - 1)) {

\# clusters do not have DE genes with themselves, so skip to next iteration

if (i == j) {

next

}

\# if an (i,j) pair is empty, fill it with negation of the corresponding (j,i) fold change values

if (is.null(log2FCList[[i]][[j]])) {

log2FCList[[i]][[j]] <- -log2FCList[[j]][[i]]

}

}

\# name the log2 fold change list with cluster names

names(log2FCList[[i]]) <- uniqueClusters

}

\# For the last row and last column

for (i in 1:(length(uniqueClusters) - 1)) {

\# set the log2 fold changes of (last_row, i) as the negation of log2 fold changes of (i, last_row)

log2FCList[[length(uniqueClusters)]][[i]] <-

-log2FCList[[i]][[length(uniqueClusters)]]

\# name the log2 fold change list with cluster names

names(log2FCList[[length(uniqueClusters)]])[i] <-

uniqueClusters[i]

}

\## Make the q values matrix symmetric, as it is currrently an upper triangular matrix

\# First cover every element barring the last row and last column

for (i in 1:(length(uniqueClusters) - 1)) {

for (j in 1:(length(uniqueClusters) - 1)) {

\# clusters do not have DE genes with themselves, so skip to next iteration

if (i == j) {

next

}

\# if an (i,j) pair is empty, fill it with the corresponding (j,i) q-values

if (is.null(qValueList[[i]][[j]])) {

qValueList[[i]][[j]] <- qValueList[[j]][[i]]

}

}

\# name the q-value list with cluster names

names(qValueList[[i]]) <- uniqueClusters

}

\# For the last row and last column

for (i in 1:(length(uniqueClusters) - 1)) {

\# set the q-values of (last_row, i) as the q-values of (i, last_row)

qValueList[[length(uniqueClusters)]][[i]] <-

qValueList[[i]][[length(uniqueClusters)]]

\# name the q-value list with cluster names

names(qValueList[[length(uniqueClusters)]])[i] <-

uniqueClusters[i]

}

\### Construct de gene count matrix

\# initialise zeroes matrix

deCountMatrix <-

matrix(

data = 0,

nrow = length(uniqueClusters),

ncol = length(uniqueClusters)

)

\# name the de gene count matrix with cluster names

rownames(deCountMatrix) <- uniqueClusters

colnames(deCountMatrix) <- uniqueClusters

\# for each pair of clusters

for (i in 1:length(uniqueClusters)) {

for (j in 1:length(uniqueClusters)) {

\# clusters do not have DE genes with themselves, so count = 0

if (i == j) {

deCountMatrix[i, j] <- 0

}

\# number of DE genes in (last_row, j) = number of elements in the (j, last_row)

else if (i == length(uniqueClusters)) {

deCountMatrix[i, j] <- length(deGeneList[[j]][[i]])

}

\# if DE gene list is empty, count = 0

else if (is.null(deGeneList[[i]][[j]])) {

deCountMatrix[i, j] <- 0

}

\# else, DE gene count (i,j) = number of DE genes at (i,j)

else {

deCountMatrix[i, j] <- length(deGeneList[[i]][[j]])

}

}

}

\### Construct de gene list with up/down regulatory information

\# initialize empty list of lists

deGeneRegulationList <-

rep(list(list()), length(uniqueClusters))

\# name DE gene regulation list with cluster names

names(deGeneRegulationList) <- uniqueClusters

\# For each pair of clusters

for (i in 1:length(uniqueClusters)) {

for (j in 1:length(uniqueClusters)) {

\# clusters do not have DE genes with themselves, so skip to next iteration

if (i == j) {

next

}

\# store fold change vector for clusters (i,j)

fcVec <- log2FCList[[i]][[j]]

\# name fold change vector with gene names

names(fcVec) <- rownames(dataIn)

\# extract fold changes for DE genes of (i,j)

fcVec <- fcVec[deGeneList[[i]][[j]]]

\# obtain up and down regulated DE genes from DE gene fold change vector

upDEGenes <- names(which(fcVec > 0))

downDEGenes <- names(which(fcVec < 0))

\# save up and down regulated genes in list

deGeneRegulationList[[i]][[j]] <-

list(

"upregulated_de_genes" = upDEGenes,

"downregulated_de_genes" = downDEGenes

)

\# name DE gene regulation list with cluster names

names(deGeneRegulationList[[i]])[j] <-

uniqueClusters[j]

}

}

\### For each cluster, obtain intersection of upregulated and downregulated DE genes against every other cluster

upregulatedDEGeneList <-

rep(list(), length(deGeneRegulationList))

downregulatedDEGeneList <-

rep(list(), length(deGeneRegulationList))

for (i in 1:length(deGeneRegulationList)) {

\# initialise empty list for each cluster

upregulatedDEGeneList[[names(deGeneRegulationList)[i]]] <-

c()

downregulatedDEGeneList[[names(deGeneRegulationList)[i]]] <-

c()

for (j in 1:length(deGeneRegulationList[[i]])) {

\# clusters do not have DE genes with themselves, so skip to next iteration

if (i == j)

next

\# obtain the union of existing upregulated DE genes for that cluster with upregulated DE genes for cluster pair (i,j)

upregulatedDEGeneList[[names(deGeneRegulationList)[i]]] <-

union(upregulatedDEGeneList[[names(deGeneRegulationList)[i]]],

deGeneRegulationList[[i]][[j]]$upregulated_de_genes)

\# obtain the union of existing upregulated DE genes for that cluster with upregulated DE genes for cluster pair (i,j)

downregulatedDEGeneList[[names(deGeneRegulationList)[i]]] <-

union(

downregulatedDEGeneList[[names(deGeneRegulationList)[i]]],

deGeneRegulationList[[i]][[j]]$downregulated_de_genes

)

}

for (j in 1:length(deGeneRegulationList[[i]])) {

\# clusters do not have DE genes with themselves, so skip to next iteration

if (i == j)

next

\# obtain the intersection of existing upregulated DE genes for that cluster with upregulated DE genes for cluster pair (i,j)

upregulatedDEGeneList[[names(deGeneRegulationList)[i]]] <-

intersect(upregulatedDEGeneList[[names(deGeneRegulationList)[i]]],

deGeneRegulationList[[i]][[j]]$upregulated_de_genes)

\# obtain the intersection of existing upregulated DE genes for that cluster with upregulated DE genes for cluster pair (i,j)

downregulatedDEGeneList[[names(deGeneRegulationList)[i]]] <-

intersect(

downregulatedDEGeneList[[names(deGeneRegulationList)[i]]],

deGeneRegulationList[[i]][[j]]$downregulated_de_genes

)

}

\# name upregulated DE gene list with cluster names

upregulatedDEGeneList[[names(deGeneRegulationList)[i]]] <-

unique(upregulatedDEGeneList[[names(deGeneRegulationList)[i]]])

\# name upregulated DE gene list with cluster names

downregulatedDEGeneList[[names(deGeneRegulationList)[i]]] <-

unique(downregulatedDEGeneList[[names(deGeneRegulationList)[i]]])

}

\### Create and initialise object to return to function caller

returnObj = list(

"qValueList" = qValueList,

"log2FCList" = log2FCList,

"deGeneList" = deGeneList,

"deGeneUnion" = deGeneUnion,

"deCountMatrix" = deCountMatrix,

"deGeneRegulationList" = deGeneRegulationList,

"upregulatedDEGeneList" = upregulatedDEGeneList,

"downregulatedDEGeneList" = downregulatedDEGeneList

\# "cellTree" = cellTree,

\# "dynamicColors" = dynamicColorsList

)

save(returnObj, file = paste0(folderOut, "/", "de_gene_object.rds"))

\### Return object

return(returnObj)

}

\# Redefine groundtruth

\# Seurat 3.1

setwd("R:/CSB/CSB6/Private/Bobby/rna-seq-pipelines/")

library(stringr)

library(gplots)

library(RColorBrewer)

library(irlba)

library(Seurat)

library(RCAv2)

library(dplyr)

library(mclust)

###################################################CBMC############################################################################

\# load the original data

data_original = read.csv("../feature_selection/raw_datasets/GSE100866_CBMC_8K_13AB_10X-RNA_umi.csv.gz",sep=",",header=T,row.names=1)

cbmc.rna.collapsed <- CollapseSpeciesExpressionMatrix(data_original)

cbmc <- CreateSeuratObject(cbmc.rna.collapsed, min.cells = 100, min.features = 500, project = "CBMC")

mito.features <- grep(pattern = "^MT-", x = rownames(x = cbmc), value = TRUE)

percent.mito <- Matrix::colSums(x = GetAssayData(object = cbmc, slot = 'counts')[mito.features, ]) / Matrix::colSums(x = GetAssayData(object = cbmc, slot = 'counts'))

cbmc[['percent.mito']] <- percent.mito

VlnPlot(object = cbmc, features = c("nFeature_RNA", "nCount_RNA", "percent.mito"), ncol = 3)

cbmc <- subset(x = cbmc, subset = nFeature_RNA > 500 & nFeature_RNA < 2000 & percent.mito < 0.08)

cbmc <- NormalizeData(cbmc)

cbmc <- FindVariableFeatures(cbmc,selection.method = "vst",nfeatures = 1000)

cbmc <- ScaleData(cbmc, display.progress = FALSE)

cbmc <- RunPCA(cbmc,features = VariableFeatures(object = cbmc))

ElbowPlot(cbmc)

cbmc <- FindNeighbors(object = cbmc, reduction = "pca", dims = 1:15)

cbmc <- FindClusters(cbmc, resolution = 0.2)

cbmc <- RunTSNE(cbmc, dims = 1:15)

cbmc.markers <- FindAllMarkers(object = cbmc, only.pos = TRUE, min.pct = 0.25, logfc.threshold = 0.25)

cbmc.markers %>% group_by(cluster) %>% top_n(n = 2, wt = avg_logFC) %>% print(n=26)

new.cluster.ids <- c("CD4 T", "CD14+ Mono", "NK", "Mouse","CD8 T", "FCGR3A+ Mono","B",

"Mk", "Precursors", "DC","Eryth")

names(x = new.cluster.ids) <- levels(x = cbmc)

cbmc <- RenameIdents(object = cbmc, new.cluster.ids)

DimPlot(object = cbmc, reduction = 'tsne', label = TRUE, pt.size = 0.5)

\# DoHeatmap(object = cbmc, features = cbmc.markers$gene, slot = "data", size = 3)

table(cbmc@active.ident)

cbmc[["rna_11clusters"]] <- Idents(object = cbmc)

\# load the ADT UMI data

cbmc.adt <- read.csv("../feature_selection/raw_datasets/GSE100866_CBMC_8K_13AB_10X-ADT_umi.csv",sep = ",", header = TRUE, row.names = 1)

cbmc.adt = cbmc.adt[,names(cbmc@active.ident)]

cbmc.adt = cbmc.adt[setdiff(rownames(cbmc.adt),c("CCR5","CCR7","CD10")),]

cbmc.cite <- CreateSeuratObject(counts = cbmc.adt)

cbmc.cite = NormalizeData(object = cbmc.cite, method = "CLR")

cbmc.cite <- ScaleData(cbmc.cite, features = NULL, display.progress = FALSE)

\###### cluster based on ADT data

\# Cluster based on PCA

cbmc.cite <- RunPCA(cbmc.cite, features = rownames(cbmc.cite@assays$RNA))

cbmc.cite <- FindNeighbors(object = cbmc.cite, reduction = "pca", dims = 1:9)

cbmc.cite <- FindClusters(cbmc.cite, resolution = 0.3, Reduction.type = "pca")

cbmc.cite[["rna_11clusters"]] <- Idents(object = cbmc)

adt_pca_labels = cbmc.cite@active.ident

cbmc.cite <- RunTSNE(cbmc.cite, dims = 1:9)

DimPlot(object = cbmc.cite, reduction = 'tsne', label = TRUE, pt.size = 0.5)

cbmc.small <- SubsetData(cbmc.cite, max.cells.per.ident = 300)

adt.markers <- FindAllMarkers(cbmc.small, only.pos = TRUE)

DoHeatmap(cbmc.small, features = unique(adt.markers$gene),size=3)

DoHeatmap(cbmc.small, features = unique(adt.markers$gene),size=3,group.by = "rna_11clusters")

\# Cluser based on UMAP

cbmc.cite <- RunUMAP(cbmc.cite,features = rownames(cbmc.cite@assays$RNA))

cbmc.cite <- FindNeighbors(object = cbmc.cite, reduction = "umap", dims = 1:2)

cbmc.cite <- FindClusters(cbmc.cite, resolution = 0.05, Reduction.type = "umap")

DimPlot(object = cbmc.cite, reduction = 'umap', label = TRUE, pt.size = 0.5)

cbmc.small <- SubsetData(cbmc.cite, max.cells.per.ident = 300)

adt.markers <- FindAllMarkers(cbmc.small, only.pos = TRUE)

DoHeatmap(cbmc.small, features = unique(adt.markers$gene),size=3)

\# load data after filtering

data=benchmarkDataList$CBMC # in normal domain after filtering and normalization

\# Intersection of data and the original data

data = data[, colnames(data) %in% colnames(cbmc@assays$RNA@data)]

\# Run RCA global panel to label cells

\# data_obj = dataConstruct(as.matrix(data))

\# data_obj = geneFilt(obj_in = data_obj)

\# data_obj = cellNormalize(data_obj)

\# data_obj = dataTransform(data_obj)

\# data_obj = featureConstruct(data_obj,method = "GlobalPanel",power = 4)

\#

\# data_obj = cellClust(data_obj,deepSplit_wgcna=0)

\# RCAPlot(data_obj)

\# Run RCA Monaco panel to label cells

data_obj = createRCAObject(rawData = log(1+data))

data_obj = dataProject(rca.obj = data_obj, method = "Custom", customPath = "../immune panels/Custom_panel_Monaco_TPM10_absolutefc5_20190320/global_custom/global_monaco_panel.rds")

data_obj = dataClust(rca.obj = data_obj, deepSplitValues = 1:4, minClustSize = 10)

plotRCAHeatmap(data_obj, folderpath = "../CSHL Single Cell Analyses/CBMC/")

plotRCAUMAP(data_obj, folderpath = "../CSHL Single Cell Analyses/CBMC/")

#get the seurat label for filtered data

groundtruth = data.frame(RCA_labels = data_obj$clustering.out$dynamicColorsList$`deepSplit 2`)

groundtruth$seurat_labels = cbmc@active.ident[colnames(data)]

groundtruth$adt_umap_labels = cbmc.cite@active.ident[colnames(data)]

groundtruth$adt_pca_labels = adt_pca_labels[colnames(data)]

#Umap using RCA global panel clustering

seed = 42

\# seurat_obj = CreateSeuratObject(data_obj$fpkm_for_clust)

\# seurat_obj <- ScaleData(object = seurat_obj, display.progress = FALSE)

\#

\# seurat_obj = RunUMAP(obj=seurat_obj,features = rownames(seurat_obj@assays$RNA@data))

\# seurat_obj@active.ident = as.factor(data_obj$group_labels_color$groupLabel)

\# names(seurat_obj@active.ident) = colnames(data)

\#

\# colors = as.character(unique(data_obj$group_labels_color$dynamicColors))

\# labels = unique(data_obj$group_labels_color$groupLabel)

\# colors_sorted = colors[sort.int(labels,index.return = T)$ix]

\# DimPlot(object = seurat_obj, reduction = 'umap',group.by = "ident",cols = colors_sorted)

\# label using seurat labels

seed = 42

seurat_obj@active.ident = as.factor(groundtruth$seurat_labels)

names(seurat_obj@active.ident) = colnames(data)

colors = unique(labels2colors(groundtruth$seurat_labels))

DimPlot(object = seurat_obj, reduction = 'umap',group.by = "ident",cols = colors)

\# label using ADT labels

seurat_obj@active.ident = as.factor(groundtruth$adt_umap_labels)

names(seurat_obj@active.ident) = colnames(data)

colors = unique(labels2colors(groundtruth$adt_umap_labels))

DimPlot(object = seurat_obj, reduction = 'umap',group.by = "ident",cols = colors)

\# source('/mnt/volume1/project/plotHeatmap.R')

\# cellTypeDEPlot(data_obj$fpkm_for_clust,cellTree = data_obj$cellTree,clusterLabels = data_obj$group_labels_color$groupLabel,dynamicColors = data_obj$dynamicColors,filename = "heatmap_RCA_Monacopanel_deepsplit2")

\# Find the consensus groundtruth based on RCA and Seurat clustering

groundtruth = as.data.frame(data_obj$clustering.out$dynamicColorsList$`deepSplit 2`)

rownames(groundtruth) = colnames(data)

colnames(groundtruth) = "RCA"

groundtruth$RNA = data_obj$seurat_labels

groundtruth$ADT_Umap = data_obj$adt_umap_labels

groundtruth$ADT_pca = data_obj$adt_pca_labels

groundtruth$consensus = 0

groundtruth[groundtruth$RCA == 1,]$consensus = "T"

groundtruth[groundtruth$ADT_pca==0 & groundtruth$RCA == 1,]$consensus = "CD4 T Naive"

groundtruth[groundtruth$ADT_pca==3 & groundtruth$RCA == 1,]$consensus = "CD4 T Memory"

groundtruth[groundtruth$ADT_pca==6 & groundtruth$RCA == 1,]$consensus = "CD8 T"

groundtruth[groundtruth$RCA == 2,]$consensus = "Mono"

groundtruth[groundtruth$ADT_pca==1 & groundtruth$RCA == 2,]$consensus = "Mono"

groundtruth[groundtruth$ADT_pca==8 & groundtruth$RCA == 2,]$consensus = "CD16 Mono"

groundtruth[groundtruth$ADT_pca==9 & groundtruth$RCA == 2,]$consensus = "CD14 Mono"

groundtruth[groundtruth$RCA == 3,]$consensus = "NK"

groundtruth[groundtruth$ADT_Umap==2 & groundtruth$RCA == 3,]$consensus = "NK"

groundtruth[groundtruth$ADT_Umap==11 & groundtruth$RCA == 3,]$consensus = "NK T"

groundtruth[groundtruth$RCA == 4,]$consensus = "B"

groundtruth[groundtruth$ADT_pca==7 & groundtruth$RCA == 4,]$consensus = "B"

groundtruth[groundtruth$ADT_pca==11 & groundtruth$RCA == 4,]$consensus = "CD3+ B"

groundtruth[groundtruth$RCA == 5,]$consensus = "Precursors"

groundtruth[groundtruth$ADT_pca==10 & groundtruth$RCA == 5,]$consensus = "Precursors"

groundtruth[groundtruth$RCA == 6,]$consensus = "DC"

groundtruth[groundtruth$RCA == 7,]$consensus = "Myeloid-like NK"

groundtruth[groundtruth$ADT_Umap==8 & groundtruth$RCA == 7,]$consensus = "Myeloid-like NK"

groundtruth[groundtruth$RCA == 8,]$consensus = "Eryth"

groundtruth[groundtruth$RCA == 10,]$consensus = "Mk"

dim(groundtruth[groundtruth$consensus==0,])

groundtruthDataList = list()

groundtruthDataList[["CBMC"]] = data[,rownames(groundtruth[groundtruth$consensus!=0,])]

groundtruthGroupList = list()

groundtruthGroupList[["CBMC"]] = groundtruth[groundtruth$consensus!=0,]$consensus

save(groundtruthDataList,groundtruthGroupList,file="groundtruth_CBMC.RData")

\# Find the consensus groundtruth based on RCA and Seurat clustering

groundtruth = as.data.frame(data_obj$group_labels_color$groupLabel)

rownames(groundtruth) = colnames(data)

colnames(groundtruth) = "RCA"

groundtruth$RNA = data_obj$seurat_labels

groundtruth$ADT_Umap = data_obj$adt_umap_labels

groundtruth$ADT_pca = data_obj$adt_pca_labels

groundtruth$consensus = 0

groundtruth[groundtruth$RCA == 1,]$consensus = "T1"

groundtruth[groundtruth$ADT_pca==0 & groundtruth$RCA == 1,]$consensus = "CD4 T Naive"

groundtruth[groundtruth$ADT_pca==3 & groundtruth$RCA == 1,]$consensus = "CD4 T Memory"

groundtruth[groundtruth$ADT_pca==6 & groundtruth$RCA == 1,]$consensus = "CD8 T"

groundtruth[groundtruth$RCA == 2,]$consensus = "Mono"

groundtruth[groundtruth$ADT_pca==1 & groundtruth$RCA == 2,]$consensus = "Mono"

groundtruth[groundtruth$ADT_pca==8 & groundtruth$RCA == 2,]$consensus = "CD16 Mono"

groundtruth[groundtruth$ADT_pca==9 & groundtruth$RCA == 2,]$consensus = "CD14 Mono"

groundtruth[groundtruth$RCA == 3,]$consensus = "NK"

groundtruth[groundtruth$ADT_Umap==2 & groundtruth$RCA == 3,]$consensus = "NK"

groundtruth[groundtruth$ADT_Umap==11 & groundtruth$RCA == 3,]$consensus = "NK T"

groundtruth[groundtruth$RCA == 4,]$consensus = "B"

groundtruth[groundtruth$ADT_pca==7 & groundtruth$RCA == 4,]$consensus = "B"

groundtruth[groundtruth$ADT_pca==11 & groundtruth$RCA == 4,]$consensus = "CD3+ B"

groundtruth[groundtruth$RCA == 5 | groundtruth$RCA == 15,]$consensus = "CD16 Mono"

groundtruth[groundtruth$ADT_pca==8 & groundtruth$consensus == "CD16 Mono",]$consensus = "CD16 Mono"

groundtruth[groundtruth$ADT_pca==4 & groundtruth$consensus == "CD16 Mono",]$consensus = "Myeloid-like NK"

groundtruth[groundtruth$RCA == 6,]$consensus = "Progenitor"

groundtruth[groundtruth$RCA == 7,]$consensus = "mDC"

groundtruth[groundtruth$RCA == 8,]$consensus = "pDC"

groundtruth[groundtruth$RCA == 9,]$consensus = "T2"

groundtruth[groundtruth$RCA == 10,]$consensus = "Neutrophils"

groundtruth[groundtruth$RCA == 11 | groundtruth$RCA == 12,]$consensus = "Myeloid-like NK"

groundtruth[groundtruth$RCA == 13,]$consensus = "Progenitor_B"

groundtruth[groundtruth$RCA == 14,]$consensus = "CD3+ B"

dim(groundtruth[groundtruth$consensus==0,])

sort(table(groundtruth$consensus))

groundtruthDataList = list()

groundtruthDataList[["CBMC"]] = data[,rownames(groundtruth[groundtruth$consensus!=0,])]

groundtruthGroupList = list()

groundtruthGroupList[["CBMC"]] = groundtruth[groundtruth$consensus!=0,]$consensus

save(groundtruthDataList,groundtruthGroupList,file="groundtruth_CBMC_Monaco.RData")

#################################################PBMC Dropseq###########################################################

\# load the original data

data_original = read.csv("../feature_selection/raw_datasets/GSE100866_PBMC_vs_flow_10X-RNA_umi.csv.gz",sep=",",header=T,row.names=1)

pbmc.rna.collapsed <- CollapseSpeciesExpressionMatrix(data_original)

pbmc <- CreateSeuratObject(pbmc.rna.collapsed,min.cells = 100, min.features = 500, project = "pbmc")

mito.features <- grep(pattern = "^MT-", x = rownames(x = pbmc), value = TRUE)

percent.mito <- Matrix::colSums(x = GetAssayData(object = pbmc, slot = 'counts')[mito.features, ]) / Matrix::colSums(x = GetAssayData(object = pbmc, slot = 'counts'))

pbmc[['percent.mito']] <- percent.mito

VlnPlot(object = pbmc, features = c("nFeature_RNA", "nCount_RNA", "percent.mito"), ncol = 3)

pbmc <- subset(x = pbmc, subset = nFeature_RNA > 500 & nFeature_RNA < 2000 & percent.mito < 0.08)

pbmc <- NormalizeData(pbmc)

pbmc <- FindVariableFeatures(pbmc,selection.method = "vst",nfeatures = 1000)

pbmc <- ScaleData(pbmc, display.progress = FALSE)

pbmc <- RunPCA(pbmc,features = VariableFeatures(object = pbmc))

ElbowPlot(pbmc)

pbmc <- FindNeighbors(object = pbmc, reduction = "pca", dims = 1:15)

pbmc <- FindClusters(pbmc, resolution = 0.2)

pbmc <- RunTSNE(pbmc, dims = 1:15)

pbmc.markers <- FindAllMarkers(object = pbmc, only.pos = TRUE, min.pct = 0.25, logfc.threshold = 0.25)

pbmc.markers %>% group_by(cluster) %>% top_n(n = 5, wt = avg_logFC) %>% print(n=26)

new.cluster.ids <- c("CD4 T", "CD14+ Mono", "NK", "B","DC", "CD8 T","FCGR3A+ Mono", "Mouse")

names(x = new.cluster.ids) <- levels(x = pbmc)

pbmc <- RenameIdents(object = pbmc, new.cluster.ids)

DimPlot(object = pbmc, reduction = 'tsne', label = TRUE, pt.size = 0.5)

table(pbmc@active.ident)

pbmc[["rna_8clusters"]] <- Idents(object = pbmc)

\# load the ADT UMI data

pbmc.adt <- read.csv("/mnt/volume1/project/pbmc_10x_8k/GSE100866_PBMC_vs_flow_10X-ADT_umi.csv",sep = ",", header = TRUE, row.names = 1)

pbmc.adt = pbmc.adt[,names(pbmc@active.ident)]

pbmc.cite <- CreateSeuratObject(counts = pbmc.adt)

pbmc.cite = NormalizeData(object = pbmc.cite, method = "CLR")

pbmc.cite <- ScaleData(pbmc.cite, features = NULL, display.progress = FALSE)

\###### cluster based on ADT data

\# Cluster based on PCA

pbmc.cite <- RunPCA(pbmc.cite, features = rownames(pbmc.cite@assays$RNA))

pbmc.cite <- FindNeighbors(object = pbmc.cite, reduction = "pca", dims = 1:9)

pbmc.cite <- FindClusters(pbmc.cite, resolution = 0.5, Reduction.type = "pca")

pbmc.cite[["rna_8clusters"]] <- Idents(object = pbmc)

adt_pca_labels = pbmc.cite@active.ident

pbmc.cite <- RunTSNE(pbmc.cite, dims = 1:9)

DimPlot(object = pbmc.cite, reduction = 'tsne', label = TRUE, pt.size = 0.5)

pbmc.small <- SubsetData(pbmc.cite, max.cells.per.ident = 300)

adt.markers <- FindAllMarkers(pbmc.small, only.pos = TRUE)

DoHeatmap(pbmc.small, features = unique(adt.markers$gene),size=3)

DoHeatmap(pbmc.small, features = unique(adt.markers$gene),size=3,group.by = "rna_8clusters")

\# Cluser based on UMAP

pbmc.cite <- RunUMAP(pbmc.cite,features = rownames(pbmc.cite@assays$RNA))

pbmc.cite <- FindNeighbors(object = pbmc.cite, reduction = "umap", dims = 1:2)

pbmc.cite <- FindClusters(pbmc.cite, resolution = 0.16, Reduction.type = "umap")

DimPlot(object = pbmc.cite, reduction = 'umap', label = TRUE, pt.size = 0.5)

pbmc.small <- SubsetData(pbmc.cite, max.cells.per.ident = 300)

adt.markers <- FindAllMarkers(pbmc.small, only.pos = TRUE)

DoHeatmap(pbmc.small, features = unique(adt.markers$gene),size=3)

\# load data after filtering

data=benchmarkDataList$`PBMC 10X` # in normal domain after filtering

\# Intersection of data and the original data

data = data[, colnames(data) %in% colnames(pbmc@assays$RNA@data)]

\# Run RCA global panel to label cells

data_obj = dataConstruct(as.matrix(data))

data_obj = geneFilt(obj_in = data_obj)

data_obj = cellNormalize(data_obj)

data_obj = dataTransform(data_obj)

data_obj = featureConstruct(data_obj,method = "GlobalPanel",power = 4)

data_obj = cellClust(data_obj,deepSplit_wgcna=0)

RCAPlot(data_obj)

\# Run RCA Monaco panel to label cells

data_obj = dataConstruct(as.matrix(data))

data_obj = geneFilt(obj_in = data_obj)

data_obj = cellNormalize(data_obj)

data_obj = dataTransform(data_obj)

source('/mnt/volume1/project/code/featureConstructMonaco.R')

data_obj = featureConstructMonaco(data_obj,method = "MonacoPanel",power = 4)

data_obj = cellClust(data_obj,deepSplit_wgcna=3)

RCAPlot(data_obj)

#get the seurat label for filtered data

data_obj$seurat_labels = pbmc@active.ident[colnames(data)]

data_obj$adt_umap_labels = pbmc.cite@active.ident[colnames(data)]

data_obj$adt_pca_labels = adt_pca_labels[colnames(data)]

#Umap using RCA global panel clustering

seed = 42

seurat_obj = CreateSeuratObject(data_obj$fpkm_for_clust)

seurat_obj <- ScaleData(object = seurat_obj, display.progress = FALSE)

seurat_obj = RunUMAP(obj=seurat_obj,features = rownames(seurat_obj@assays$RNA@data))

seurat_obj@active.ident = as.factor(data_obj$group_labels_color$groupLabel)

names(seurat_obj@active.ident) = colnames(data)

colors = as.character(unique(data_obj$group_labels_color$dynamicColors))

labels = unique(data_obj$group_labels_color$groupLabel)

colors_sorted = colors[sort.int(labels,index.return = T)$ix]

DimPlot(object = seurat_obj, reduction = 'umap',group.by = "ident",cols = colors_sorted)

\# label using seurat labels

seurat_obj@active.ident = as.factor(data_obj$seurat_labels)

names(seurat_obj@active.ident) = colnames(data)

colors = unique(labels2colors(data_obj$seurat_labels))

DimPlot(object = seurat_obj, reduction = 'umap',group.by = "ident",cols = colors)

\# label using ADT labels

seurat_obj@active.ident = as.factor(data_obj$adt_pca_labels)

names(seurat_obj@active.ident) = colnames(data)

colors = unique(labels2colors(data_obj$adt_pca_labels))

DimPlot(object = seurat_obj, reduction = 'umap',group.by = "ident",cols = colors)

source('/mnt/volume1/project/plotHeatmap.R')

cellTypeDEPlot(data_obj$fpkm_for_clust,cellTree = data_obj$cellTree,clusterLabels = data_obj$group_labels_color$groupLabel,dynamicColors = data_obj$dynamicColors,filename = "heatmap_RCA_Monacopanel_deepsplit3")

\# Find the consensus groundtruth based on RCA and Seurat clustering

groundtruth = as.data.frame(data_obj$group_labels_color$groupLabel)

rownames(groundtruth) = colnames(data)

colnames(groundtruth) = "RCA"

groundtruth$RNA = data_obj$seurat_labels

groundtruth$ADT_Umap = data_obj$adt_umap_labels

groundtruth$ADT_pca = data_obj$adt_pca_labels

groundtruth$consensus = 0

groundtruth[groundtruth$RCA == 1,]$consensus = "T"

groundtruth[groundtruth$ADT_pca==0 & groundtruth$RCA == 1,]$consensus = "CD4 T Memory"

groundtruth[groundtruth$ADT_pca==1 & groundtruth$RCA == 1,]$consensus = "CD4 T Naive"

groundtruth[groundtruth$ADT_pca==5 & groundtruth$RCA == 1,]$consensus = "CD8 T Naive"

groundtruth[groundtruth$ADT_pca==6 & groundtruth$RCA == 1,]$consensus = "CD8 T Memory"

groundtruth[groundtruth$ADT_pca==8 & groundtruth$RCA == 1,]$consensus = "NK T"

groundtruth[groundtruth$RCA == 2,]$consensus = "Mono"

groundtruth[groundtruth$ADT_pca==3 & groundtruth$RCA == 2,]$consensus = "Mono"

groundtruth[groundtruth$ADT_pca==7 & groundtruth$RCA == 2,]$consensus = "CD14 Mono"

groundtruth[groundtruth$ADT_pca==9 & groundtruth$RCA == 2,]$consensus = "CD14 Mono"

groundtruth[groundtruth$RCA == 3,]$consensus = "NK"

groundtruth[groundtruth$ADT_pca==2 & groundtruth$RCA == 3,]$consensus = "NK"

groundtruth[groundtruth$ADT_pca==5 & groundtruth$RCA == 3,]$consensus = "CD8 T Naive"

groundtruth[groundtruth$ADT_pca==6 & groundtruth$RCA == 3,]$consensus = "CD8 T Memory"

groundtruth[groundtruth$ADT_pca==8 & groundtruth$RCA == 3,]$consensus = "NK T"

groundtruth[groundtruth$RCA == 4,]$consensus = "B"

groundtruth[groundtruth$ADT_pca==4 & groundtruth$RCA == 4,]$consensus = "B"

groundtruth[groundtruth$ADT_pca==11 & groundtruth$RCA == 4,]$consensus = "CD3+ B"

groundtruth[groundtruth$RCA == 5,]$consensus = "DC"

groundtruth[groundtruth$RCA == 6,]$consensus = "Myeloid-like NK"

groundtruth[groundtruth$RCA == 7,]$consensus = "Precursors"

groundtruth[groundtruth$RCA == 8,]$consensus = "Plasma"

dim(groundtruth[groundtruth$consensus==0,])

groundtruthDataList = list()

groundtruthDataList[["pbmc_dropseq"]] = data[,rownames(groundtruth[groundtruth$consensus!=0,])]

groundtruthGroupList = list()

groundtruthGroupList[["pbmc_dropseq"]] = groundtruth[groundtruth$consensus!=0,]$consensus

save(groundtruthDataList,groundtruthGroupList,file="groundtruth_PBMC_dropseq.RData")

\# Find the consensus groundtruth based on RCA and Seurat clustering

groundtruth = as.data.frame(data_obj$group_labels_color$groupLabel)

rownames(groundtruth) = colnames(data)

colnames(groundtruth) = "RCA"

groundtruth$RNA = data_obj$seurat_labels

groundtruth$ADT_Umap = data_obj$adt_umap_labels

groundtruth$ADT_pca = data_obj$adt_pca_labels

groundtruth$consensus = 0

groundtruth[groundtruth$RCA == 1,]$consensus = "T"

groundtruth[groundtruth$ADT_pca==0 & groundtruth$RCA == 1,]$consensus = "CD4 T Memory"

groundtruth[groundtruth$ADT_pca==1 & groundtruth$RCA == 1,]$consensus = "CD4 T Naive"

groundtruth[groundtruth$ADT_pca==5 & groundtruth$RCA == 1,]$consensus = "CD8 T Naive"

groundtruth[groundtruth$ADT_pca==6 & groundtruth$RCA == 1,]$consensus = "CD8 T Memory"

groundtruth[groundtruth$ADT_pca==8 & groundtruth$RCA == 1,]$consensus = "NK T"

groundtruth[groundtruth$RCA == 2,]$consensus = "NK"

groundtruth[groundtruth$ADT_pca==2 & groundtruth$RCA == 2,]$consensus = "NK"

groundtruth[groundtruth$ADT_pca==5 & groundtruth$RCA == 2,]$consensus = "CD8 T Naive"

groundtruth[groundtruth$ADT_pca==6 & groundtruth$RCA == 2,]$consensus = "CD8 T Memory"

groundtruth[groundtruth$ADT_pca==1 & groundtruth$RCA == 2,]$consensus = "CD4 T Naive"

groundtruth[groundtruth$ADT_pca==8 & groundtruth$RCA == 2,]$consensus = "NK T"

groundtruth[groundtruth$RCA == 3 | groundtruth$RCA == 17,]$consensus = "C_Mono"

groundtruth[groundtruth$RCA == 4 | groundtruth$RCA == 26,]$consensus = "CD4 T_1"

groundtruth[groundtruth$ADT_pca==0 & groundtruth$consensus == "CD4 T_1",]$consensus = "CD4 T Memory"

groundtruth[groundtruth$ADT_pca==1 & groundtruth$consensus == "CD4 T_1",]$consensus = "CD4 T Naive"

groundtruth[groundtruth$ADT_pca==6 & groundtruth$consensus == "CD4 T_1",]$consensus = "CD8 T Memory"

groundtruth[groundtruth$ADT_pca==8 & groundtruth$consensus == "CD4 T_1",]$consensus = "NK T"

groundtruth[groundtruth$RCA == 5 | groundtruth$RCA == 25 | groundtruth$RCA == 28,]$consensus = "B"

groundtruth[groundtruth$ADT_pca==11 & groundtruth$consensus == "B",]$consensus = "CD3+ B"

groundtruth[groundtruth$RCA == 6 | groundtruth$RCA == 27,]$consensus = "I_Mono"

groundtruth[groundtruth$RCA == 7,]$consensus = "CD8 T_1"

groundtruth[groundtruth$ADT_pca==6 & groundtruth$consensus == "CD8 T_1",]$consensus = "CD8 T Memory"

groundtruth[groundtruth$ADT_pca==8 & groundtruth$consensus == "CD8 T_1",]$consensus = "NK T"

groundtruth[groundtruth$ADT_pca==5 & groundtruth$consensus == "CD8 T_1",]$consensus = "CD8 T Naive"

groundtruth[groundtruth$RCA == 8 | groundtruth$RCA == 22,]$consensus = "CD4 T_2"

groundtruth[groundtruth$ADT_pca==0 & groundtruth$consensus == "CD4 T_2",]$consensus = "CD4 T Memory"

groundtruth[groundtruth$ADT_pca==6 & groundtruth$consensus == "CD4 T_2",]$consensus = "CD8 T Memory"

groundtruth[groundtruth$ADT_pca==8 & groundtruth$consensus == "CD4 T_2",]$consensus = "NK T"

groundtruth[groundtruth$ADT_pca==5 & groundtruth$consensus == "CD4 T_2",]$consensus = "CD8 T Naive"

groundtruth[groundtruth$ADT_pca==1 & groundtruth$consensus == "CD4 T_2",]$consensus = "CD4 T Naive"

groundtruth[groundtruth$RCA == 9,]$consensus = "CD8 T Naive"

groundtruth[groundtruth$ADT_pca==1 & groundtruth$consensus == "CD8 T Naive",]$consensus = "CD4 T Naive"

groundtruth[groundtruth$ADT_pca==6 & groundtruth$consensus == "CD8 T Naive",]$consensus = "CD8 T Memory"

groundtruth[groundtruth$RCA == 10,]$consensus = "MAIT"

groundtruth[groundtruth$RCA == 11 | groundtruth$RCA ==15,]$consensus = "CD8 T_2"

groundtruth[groundtruth$ADT_pca==6 & groundtruth$consensus == "CD8 T_2",]$consensus = "CD8 T Memory"

groundtruth[groundtruth$ADT_pca==5 & groundtruth$consensus == "CD8 T_2",]$consensus = "CD8 T Naive"

groundtruth[groundtruth$ADT_pca==8 & groundtruth$consensus == "CD8 T_2",]$consensus = "NK T"

groundtruth[groundtruth$ADT_pca==0 & groundtruth$consensus == "CD8 T_2",]$consensus = "CD4 T Memory"

groundtruth[groundtruth$RCA == 12 | groundtruth$RCA ==21,]$consensus = "mDC"

groundtruth[groundtruth$RCA == 13,]$consensus = "CD8 T_3"

groundtruth[groundtruth$ADT_pca==8 & groundtruth$consensus == "CD8 T_3",]$consensus = "NK T"

groundtruth[groundtruth$ADT_pca==1 & groundtruth$consensus == "CD8 T_3",]$consensus = "CD4 T Naive"

groundtruth[groundtruth$ADT_pca==0 & groundtruth$consensus == "CD8 T_3",]$consensus = "CD4 T Memory"

groundtruth[groundtruth$ADT_pca==5 & groundtruth$consensus == "CD8 T_3",]$consensus = "CD8 T Naive"

groundtruth[groundtruth$ADT_pca==6 & groundtruth$consensus == "CD8 T_3",]$consensus = "CD8 T Memory"

groundtruth[groundtruth$RCA == 14 | groundtruth$RCA == 23,]$consensus = "CD8 T_4"

groundtruth[groundtruth$ADT_pca==5 & groundtruth$consensus == "CD8 T_4",]$consensus = "CD8 T Naive"

groundtruth[groundtruth$ADT_pca==6 & groundtruth$consensus == "CD8 T_4",]$consensus = "CD8 T Memory"

groundtruth[groundtruth$ADT_pca==8 & groundtruth$consensus == "CD8 T_4",]$consensus = "NK T"

groundtruth[groundtruth$ADT_pca==1 & groundtruth$consensus == "CD8 T_4",]$consensus = "CD4 T Naive"

groundtruth[groundtruth$ADT_pca==0 & groundtruth$consensus == "CD8 T_4",]$consensus = "CD4 T Memory"

groundtruth[groundtruth$RCA == 16,]$consensus = "pDC"

groundtruth[groundtruth$RCA == 18,]$consensus = "Neutrophils"

groundtruth[groundtruth$RCA == 19,]$consensus = "Myeloid-like NK"

groundtruth[groundtruth$RCA == 20,]$consensus = "Progenitor"

groundtruth[groundtruth$RCA == 24,]$consensus = "Plasma"

groundtruth[groundtruth$consensus == "CD8 T_3" | groundtruth$consensus == "T" | groundtruth$consensus == "CD4 T_1"|

groundtruth$consensus == "CD4 T_2" | groundtruth$consensus == "CD8 T_1",]$consensus = "T"

dim(groundtruth[groundtruth$consensus==0,])

sort(table(groundtruth$consensus))

groundtruthDataList = list()

groundtruthDataList[["pbmc_dropseq"]] = data[,rownames(groundtruth[groundtruth$consensus!=0,])]

groundtruthGroupList = list()

groundtruthGroupList[["pbmc_dropseq"]] = groundtruth[groundtruth$consensus!=0,]$consensus

save(groundtruthDataList,groundtruthGroupList,file="groundtruth_PBMC_dropseq_Monaco.RData")

#######################MALT#################################################

\# load the original data

data_original = readMM("/mnt/volume1/project/malt_10k/matrix.mtx.gz")

data_genes = read.csv("/mnt/volume1/project/malt_10k/features.tsv.gz",header=F,sep="\t",stringsAsFactors = F)

data_cells = read.csv("/mnt/volume1/project/malt_10k/barcodes.tsv.gz",header=F,sep="\t",stringsAsFactors = F)

data_original@Dimnames[[1]] = data_genes$V2

data_original@Dimnames[[2]] = data_cells$V1

malt.rna.collapsed <- CollapseSpeciesExpressionMatrix(data_original[1:33538,])

malt <- CreateSeuratObject(malt.rna.collapsed,min.cells = 100, min.features = 500, project = "malt")

mito.features <- grep(pattern = "^MT-", x = rownames(x = malt), value = TRUE)

percent.mito <- Matrix::colSums(x = GetAssayData(object = malt, slot = 'counts')[mito.features, ]) / Matrix::colSums(x = GetAssayData(object = malt, slot = 'counts'))

malt[['percent.mito']] <- percent.mito

VlnPlot(object = malt, features = c("nFeature_RNA", "nCount_RNA", "percent.mito"), ncol = 3)

malt <- subset(x = malt, subset = nFeature_RNA > 500 & nFeature_RNA < 3000 & percent.mito < 0.3)

malt <- NormalizeData(malt)

malt <- FindVariableFeatures(malt,selection.method = "vst",nfeatures = 1000)

malt <- ScaleData(malt, display.progress = FALSE)

malt <- RunPCA(malt,features = VariableFeatures(object = malt))

ElbowPlot(malt)

malt <- FindNeighbors(object = malt, reduction = "pca", dims = 1:15)

malt <- FindClusters(malt, resolution = 0.2)

malt <- RunTSNE(malt, dims = 1:15)

malt.markers <- FindAllMarkers(object = malt, only.pos = TRUE, min.pct = 0.25, logfc.threshold = 0.25)

malt.markers %>% group_by(cluster) %>% top_n(n = 5, wt = avg_logFC) %>% print(n=50)

filter(malt.markers, cluster == "0") %>% top_n(n=20,wt=avg_logFC)

new.cluster.ids <- c("B", "Follicular B", "CD4 T", "CD8 T","TIGIT+ T", "B1","T", "Plasma","Macro")

names(x = new.cluster.ids) <- levels(x = malt)

malt <- RenameIdents(object = malt, new.cluster.ids)

DimPlot(object = malt, reduction = 'tsne', label = TRUE, pt.size = 0.5,label.size = 8)+NoLegend()

table(malt@active.ident)

malt[["rna_9clusters"]] <- Idents(object = malt)

\# load the ADT UMI data

malt.adt <- as.matrix(data_original[33539:33554,])

malt.adt = malt.adt[,names(malt@active.ident)]

malt.cite <- CreateSeuratObject(counts = malt.adt)

malt.cite = NormalizeData(object = malt.cite, method = "CLR")

malt.cite <- ScaleData(malt.cite, features = NULL, display.progress = FALSE)

\###### cluster based on ADT data

\# Cluster based on PCA

malt.cite <- RunPCA(malt.cite, features = rownames(malt.cite@assays$RNA))

malt.cite <- FindNeighbors(object = malt.cite, reduction = "pca", dims = 1:15)

malt.cite <- FindClusters(malt.cite, resolution = 0.65, Reduction.type = "pca")

malt.cite[["rna_9clusters"]] <- Idents(object = malt)

adt_pca_labels = malt.cite@active.ident

malt.cite <- RunTSNE(malt.cite, dims = 1:15)

DimPlot(object = malt.cite, reduction = 'tsne', label = TRUE, pt.size = 0.5)

malt.small <- SubsetData(malt.cite, max.cells.per.ident = 300)

adt.markers <- FindAllMarkers(malt.small, only.pos = TRUE)

DoHeatmap(malt.small, features = unique(adt.markers$gene),size=3)

DoHeatmap(malt.small, features = unique(adt.markers$gene),size=3,group.by = "rna_9clusters")

\# Cluser based on UMAP

malt.cite <- RunUMAP(malt.cite,features = rownames(malt.cite@assays$RNA))

malt.cite <- FindNeighbors(object = malt.cite, reduction = "umap", dims = 1:2)

malt.cite <- FindClusters(malt.cite, resolution = 0.14, Reduction.type = "umap")

DimPlot(object = malt.cite, reduction = 'umap', label = TRUE, pt.size = 0.5)

malt.small <- SubsetData(malt.cite, max.cells.per.ident = 300)

adt.markers <- FindAllMarkers(malt.small, only.pos = TRUE)

DoHeatmap(malt.small, features = unique(adt.markers$gene),size=3)

\# load data after filtering

data=benchmarkDataList$MALT # in normal domain after filtering

colnames(data) <- paste0(colnames(data),"-1")

\# Intersection of data and the original data

data = data[, colnames(data) %in% colnames(malt@assays$RNA@data)]

\# Run RCA global panel to label cells

data_obj = dataConstruct(as.matrix(data))

data_obj = geneFilt(obj_in = data_obj)

data_obj = cellNormalize(data_obj)

data_obj = dataTransform(data_obj)

data_obj = featureConstruct(data_obj,method = "GlobalPanel",power = 4)

data_obj = cellClust(data_obj,deepSplit_wgcna=0)

RCAPlot(data_obj)

\# Run RCA Monaco panel to label cells

data_obj = dataConstruct(as.matrix(data))

data_obj = geneFilt(obj_in = data_obj)

data_obj = cellNormalize(data_obj)

data_obj = dataTransform(data_obj)

source('/mnt/volume1/project/code/featureConstructMonaco.R')

data_obj = featureConstructMonaco(data_obj,method = "MonacoPanel",power = 4)

data_obj = cellClust(data_obj,deepSplit_wgcna=3)

RCAPlot(data_obj)

source('/mnt/volume1/project/plotHeatmap.R')

cellTypeDEPlot(data_obj$fpkm_for_clust,cellTree = data_obj$cellTree,clusterLabels = data_obj$group_labels_color$groupLabel,dynamicColors = data_obj$dynamicColors,filename = "heatmap_RCA_Monacopanel_deepsplit3")

\# cluster based on ADT data and label using RCA Monaco panel

malt.adt <- as.matrix(data_original[33539:33554,])

\# Intersection of data and the original data

malt.adt = malt.adt[, colnames(malt.adt) %in% colnames(data)]

malt.cite <- CreateSeuratObject(counts = malt.adt)

malt.cite = NormalizeData(object = malt.cite, method = "CLR")

malt.cite <- ScaleData(malt.cite, features = NULL, display.progress = FALSE)

\###### cluster based on ADT data

\# Cluster based on PCA

malt.cite <- RunPCA(malt.cite, features = rownames(malt.cite@assays$RNA))

malt.cite <- FindNeighbors(object = malt.cite, reduction = "pca", dims = 1:15)

malt.cite <- FindClusters(malt.cite, resolution = 0.65, Reduction.type = "pca")

malt.cite[["rca_monaco"]] <- as.factor(data_obj$group_labels_color$groupLabel)

adt_pca_labels = malt.cite@active.ident

malt.cite <- RunTSNE(malt.cite, dims = 1:15)

DimPlot(object = malt.cite, reduction = 'tsne', label = TRUE, pt.size = 0.5)

malt.small <- SubsetData(malt.cite, max.cells.per.ident = 300)

adt.markers <- FindAllMarkers(malt.small, only.pos = TRUE)

DoHeatmap(malt.small, features = unique(adt.markers$gene),size=3)

DoHeatmap(malt.small, features = unique(adt.markers$gene),size=3,group.by = "rca_monaco")

#get the seurat label for filtered data

data_obj$seurat_labels = malt@active.ident[colnames(data)]

data_obj$adt_umap_labels = malt.cite@active.ident[colnames(data)]

data_obj$adt_pca_labels = adt_pca_labels[colnames(data)]

#Umap using RCA global panel clustering

seed = 42

seurat_obj = CreateSeuratObject(data_obj$fpkm_for_clust)

seurat_obj <- ScaleData(object = seurat_obj, display.progress = FALSE)

seurat_obj = RunUMAP(obj=seurat_obj,features = rownames(seurat_obj@assays$RNA@data))

seurat_obj@active.ident = as.factor(data_obj$group_labels_color$groupLabel)

names(seurat_obj@active.ident) = colnames(data)

colors = as.character(unique(data_obj$group_labels_color$dynamicColors))

labels = unique(data_obj$group_labels_color$groupLabel)

colors_sorted = colors[sort.int(labels,index.return = T)$ix]

DimPlot(object = seurat_obj, reduction = 'umap',group.by = "ident",cols = colors_sorted)

\# label using seurat labels

seurat_obj@active.ident = as.factor(data_obj$seurat_labels)

names(seurat_obj@active.ident) = colnames(data)

colors = unique(labels2colors(data_obj$seurat_labels))

colors[6] = "pink"

colors[8] = "blue"

DimPlot(object = seurat_obj, reduction = 'umap',group.by = "ident",cols = colors)

\# label using ADT labels

seurat_obj@active.ident = as.factor(data_obj$adt_pca_labels)

names(seurat_obj@active.ident) = colnames(data)

\# highlight the marker genes for plasma cells

cells_seurat = colnames(data_obj$fpkm_for_clust[data_obj$seurat_labels == "Plasma"])

cells_rca = colnames(data_obj$fpkm_for_clust[which(data_obj$group_labels_color$groupLabel == 10 | data_obj$group_labels_color$groupLabel == 12)])

FeaturePlot(malt,cols = c("lightgrey","darkblue"), features = c("CD27"), cells = cells_seurat)

FeaturePlot(malt,cols = c("lightgrey","darkblue"), features = c("CD27"), cells = cells_rca)

FeaturePlot(malt,cols = c("lightgrey","darkblue"), features = c("PRDM1"), cells = cells_seurat)

FeaturePlot(malt,cols = c("lightgrey","darkblue"), features = c("PRDM1"), cells = cells_rca)

FeaturePlot(malt,cols = c("lightgrey","darkblue"), features = c("DERL3"), cells = cells_seurat)

FeaturePlot(malt,cols = c("lightgrey","darkblue"), features = c("DERL3"), cells = cells_rca)

library(ggplot2)

colors = c("lightgrey","lightgrey","lightgrey","lightgrey","lightgrey","lightgrey","lightgrey","lightgrey","lightgrey")

plot<-DimPlot(object = seurat_obj, reduction = 'umap',group.by = "ident", cols = colors, do.return=T)

\# plot <- plot+geom_point(data=as.data.frame(data=malt@assays$RNA@counts["CD27",cells_seurat]),

\# mapping = aes(x = seurat_obj@reductions$umap[[,1]], y = seurat_obj@reductions$umap[[,2]],color=))+scale_color_gradient(low="lightgrey", high="darkblue")

data_df <- as.data.frame(seurat_obj@reductions$)

data_df <- cbind(data_df, data.frame(cd27 = malt@assays$RNA@scale.data["CD27",rownames(data_df)]))

ggplot(data=data_df,

mapping = aes(x = UMAP_1, y = UMAP_2, color=cd27))+scale_color_gradient(low="lightgrey", high="darkblue") + geom_point()

data_df <- cbind(data_df, data.frame(PRDM1 = malt@assays$RNA@scale.data["PRDM1",rownames(data_df)]))

ggplot(data=data_df,

mapping = aes(x = UMAP_1, y = UMAP_2, color=PRDM1))+scale_color_gradient2(low="lightgrey", mid = "darkblue", high="black",midpoint = 3) + geom_point()

data_df <- cbind(data_df, data.frame(DERL3 = malt@assays$RNA@scale.data["DERL3",rownames(data_df)]))

ggplot(data=data_df,

mapping = aes(x = UMAP_1, y = UMAP_2, color=DERL3))+scale_color_gradient2(low="lightgrey", mid = "darkblue", high="black",midpoint = 3) + geom_point()

data_df <- cbind(data_df, data.frame(BCL11A = malt@assays$RNA@scale.data["BCL11A",rownames(data_df)]))

ggplot(data=data_df,

mapping = aes(x = UMAP_1, y = UMAP_2, color=BCL11A))+scale_color_gradient2(low="lightgrey", mid = "darkblue", high="black",midpoint = 3) + geom_point()

data_df <- cbind(data_df, data.frame(FCER2 = malt@assays$RNA@scale.data["FCER2",rownames(data_df)]))

ggplot(data=data_df,

mapping = aes(x = UMAP_1, y = UMAP_2, color=FCER2))+scale_color_gradient2(low="lightgrey", mid = "darkblue", high="black",midpoint = 3) + geom_point()

genes = c("IGHG3", "SSPN", "TNFRSF13B", "SLCO4A1", "RGS16", "IGLC3", "IGHG1", "DERL3", "PNOC")

data_df <- cbind(data_df, data.frame(Markers = colMeans(malt@assays$RNA@scale.data[genes[which(genes %in% rownames(malt@assays$RNA@scale.data))] ,rownames(data_df)])))

ggplot(data=data_df,

mapping = aes(x = UMAP_1, y = UMAP_2, color=Markers))+scale_color_gradient2(low="lightgrey", mid = "darkblue", high="black",midpoint = 3) + geom_point()

source('/mnt/volume1/project/plotHeatmap.R')

cellTypeDEPlot(data_obj$fpkm_for_clust,cellTree = data_obj$cellTree,clusterLabels = data_obj$group_labels_color$groupLabel,dynamicColors = data_obj$dynamicColors,filename = "heatmap_RCA_Monacopanel_deepsplit3")

\# Find the consensus groundtruth based on RCA and Seurat clustering

groundtruth = as.data.frame(data_obj$group_labels_color$groupLabel)

rownames(groundtruth) = colnames(data)

colnames(groundtruth) = "RCA"

groundtruth$RNA = data_obj$seurat_labels

groundtruth$ADT_Umap = data_obj$adt_umap_labels

groundtruth$ADT_pca = data_obj$adt_pca_labels

groundtruth$consensus = 0

groundtruth[groundtruth$RCA == 1,]$consensus = "B"

groundtruth[groundtruth$ADT_pca==0 & groundtruth$RCA == 1,]$consensus = "CD14- B"

groundtruth[groundtruth$ADT_pca==1 & groundtruth$RCA == 1,]$consensus = "Activated B"

groundtruth[groundtruth$ADT_pca==3 & groundtruth$RCA == 1,]$consensus = "CD14- activated B"

groundtruth[groundtruth$RCA == 2,]$consensus = "T"

groundtruth[groundtruth$ADT_pca==4 & groundtruth$RCA == 2,]$consensus = "CD127+ Memory CD4 T"

groundtruth[groundtruth$ADT_pca==5 & groundtruth$RCA == 2,]$consensus = "CD8 T Memory"

groundtruth[groundtruth$ADT_pca==6 & groundtruth$RCA == 2,]$consensus = "TIGIT+ CD4 T Memory"

groundtruth[groundtruth$ADT_pca==7 & groundtruth$RCA == 2,]$consensus = "PD1+ CD4 T Memory"

groundtruth[groundtruth$ADT_pca==8 & groundtruth$RCA == 2,]$consensus = "CD4 T Memory"

groundtruth[groundtruth$ADT_pca==9 & groundtruth$RCA == 2,]$consensus = "CD127+ CD8 T"

groundtruth[groundtruth$ADT_pca==10 & groundtruth$RCA == 2,]$consensus = "CD127+ CD4 T"

groundtruth[groundtruth$RCA == 3,]$consensus = "Lymphnode"

groundtruth[groundtruth$ADT_pca==6 & groundtruth$RCA == 3,]$consensus = "TIGIT+ CD4 T Memory"

groundtruth[groundtruth$RCA == 4,]$consensus = "Lymphoma B"

groundtruth[groundtruth$RCA == 5,]$consensus = "Plasma"

groundtruth[groundtruth$RCA == 6,]$consensus = "Macro"

groundtruth[groundtruth$RCA == 7,]$consensus = "DC"

groundtruth[groundtruth$RCA == 8,]$consensus = "NK"

groundtruth[groundtruth$RCA == 9,]$consensus = "Thymus"

dim(groundtruth[groundtruth$consensus==0,])

table(groundtruth$consensus)

groundtruthDataList = list()

groundtruthDataList[["malt"]] = data[,rownames(groundtruth[groundtruth$consensus!=0,])]

groundtruthGroupList = list()

groundtruthGroupList[["malt"]] = groundtruth[groundtruth$consensus!=0,]$consensus

save(groundtruthDataList,groundtruthGroupList,file="groundtruth_malt.RData")

\# Find the consensus groundtruth based on RCA and Seurat clustering

groundtruth = as.data.frame(data_obj$group_labels_color$groupLabel)

rownames(groundtruth) = colnames(data)

colnames(groundtruth) = "RCA"

groundtruth$RNA = data_obj$seurat_labels

groundtruth$ADT_Umap = data_obj$adt_umap_labels

groundtruth$ADT_pca = data_obj$adt_pca_labels

groundtruth$consensus = 0

groundtruth[groundtruth$RCA == 1,]$consensus = "B1"

groundtruth[groundtruth$ADT_pca==0 & groundtruth$consensus == "B1",]$consensus = "B"

groundtruth[groundtruth$ADT_pca==1 & groundtruth$consensus == "B1",]$consensus = "Activated B"

groundtruth[groundtruth$ADT_pca==2 & groundtruth$consensus == "B1",]$consensus = "CD14- activated B"

groundtruth[groundtruth$ADT_pca==3 & groundtruth$consensus == "B1",]$consensus = "CD14- B"

groundtruth[groundtruth$RCA == 2,]$consensus = "B2"

groundtruth[groundtruth$ADT_pca==0 & groundtruth$consensus == "B2",]$consensus = "B"

groundtruth[groundtruth$ADT_pca==1 & groundtruth$consensus == "B2",]$consensus = "Activated B"

groundtruth[groundtruth$ADT_pca==2 & groundtruth$consensus == "B2",]$consensus = "CD14- activated B"

groundtruth[groundtruth$ADT_pca==3 & groundtruth$consensus == "B2",]$consensus = "CD14- B"

groundtruth[groundtruth$RCA == 3 | groundtruth$RCA == 22,]$consensus = "T1"

groundtruth[groundtruth$ADT_pca==4 & groundtruth$consensus == "T1",]$consensus = "CD127+ Memory CD4 T"

groundtruth[groundtruth$ADT_pca==5 & groundtruth$consensus == "T1",]$consensus = "TIGIT+ CD4 T Memory"

groundtruth[groundtruth$ADT_pca==6 & groundtruth$consensus == "T1",]$consensus = "CD8 T Memory"

groundtruth[groundtruth$ADT_pca==7 & groundtruth$consensus == "T1",]$consensus = "PD1+ CD4 T Memory"

groundtruth[groundtruth$ADT_pca==8 & groundtruth$consensus == "T1",]$consensus = "CD127+ CD8 T"

########################### stop here, do not do Monaco panel for MALT

groundtruth[groundtruth$RCA == 4,]$consensus = "Lymphoma B"

groundtruth[groundtruth$RCA == 5,]$consensus = "Plasma"

groundtruth[groundtruth$RCA == 6,]$consensus = "Macro"

groundtruth[groundtruth$RCA == 7,]$consensus = "DC"

groundtruth[groundtruth$RCA == 8,]$consensus = "NK"

groundtruth[groundtruth$RCA == 9,]$consensus = "Thymus"

dim(groundtruth[groundtruth$consensus==0,])

table(groundtruth$consensus)

groundtruthDataList = list()

groundtruthDataList[["malt"]] = data[,rownames(groundtruth[groundtruth$consensus!=0,])]

groundtruthGroupList = list()

groundtruthGroupList[["malt"]] = groundtruth[groundtruth$consensus!=0,]$consensus

save(groundtruthDataList,groundtruthGroupList,file="groundtruth_malt.RData")

###############################PBMC###############################################

\# load the original data

data_original = readMM("/mnt/volume1/project/pbmc_10k/matrix.mtx.gz")

data_genes = read.csv("/mnt/volume1/project/pbmc_10k/features.tsv.gz",header=F,sep="\t",stringsAsFactors = F)

data_cells = read.csv("/mnt/volume1/project/pbmc_10k/barcodes.tsv.gz",header=F,sep="\t",stringsAsFactors = F)

data_original@Dimnames[[1]] = data_genes$V2

data_original@Dimnames[[2]] = data_cells$V1

pbmc.rna.collapsed <- CollapseSpeciesExpressionMatrix(data_original[1:33538,])

pbmc <- CreateSeuratObject(pbmc.rna.collapsed,min.cells = 100, min.features = 500, project = "pbmc")

mito.features <- grep(pattern = "^MT-", x = rownames(x = pbmc), value = TRUE)

percent.mito <- Matrix::colSums(x = GetAssayData(object = pbmc, slot = 'counts')[mito.features, ]) / Matrix::colSums(x = GetAssayData(object = pbmc, slot = 'counts'))

pbmc[['percent.mito']] <- percent.mito

VlnPlot(object = pbmc, features = c("nFeature_RNA", "nCount_RNA", "percent.mito"), ncol = 3)

pbmc <- subset(x = pbmc, subset = nFeature_RNA > 500 & nFeature_RNA < 3000 & percent.mito < 0.15)

pbmc <- NormalizeData(pbmc)

pbmc <- FindVariableFeatures(pbmc,selection.method = "vst",nfeatures = 1000)

pbmc <- ScaleData(pbmc, display.progress = FALSE)

pbmc <- RunPCA(pbmc,features = VariableFeatures(object = pbmc))

ElbowPlot(pbmc)

pbmc <- FindNeighbors(object = pbmc, reduction = "pca", dims = 1:15)

pbmc <- FindClusters(pbmc, resolution = 0.1)

pbmc <- RunTSNE(pbmc, dims = 1:15)

pbmc.markers <- FindAllMarkers(object = pbmc, only.pos = TRUE, min.pct = 0.25, logfc.threshold = 0.25)

pbmc.markers %>% group_by(cluster) %>% top_n(n = 5, wt = avg_logFC) %>% print(n=26)

filter(pbmc.markers, cluster == "0") %>% top_n(n=20,wt=avg_logFC)

new.cluster.ids <- c("CD14 Mono", "CD4 T", "CD4 T","CD8 T", "NK", "B", "Plasma","DC")

names(x = new.cluster.ids) <- levels(x = pbmc)

pbmc <- RenameIdents(object = pbmc, new.cluster.ids)

DimPlot(object = pbmc, reduction = 'tsne', label = TRUE, pt.size = 0.5)

table(pbmc@active.ident)

pbmc[["rna_7clusters"]] <- Idents(object = pbmc)

\# load the ADT UMI data

pbmc.adt <- as.matrix(data_original[33539:33555,])

pbmc.adt = pbmc.adt[,names(pbmc@active.ident)]

pbmc.cite <- CreateSeuratObject(counts = pbmc.adt)

pbmc.cite = NormalizeData(object = pbmc.cite, method = "CLR")

pbmc.cite <- ScaleData(pbmc.cite, features = NULL, display.progress = FALSE)

\###### cluster based on ADT data

\# Cluster based on PCA

seed = 1

pbmc.cite <- RunPCA(pbmc.cite, features = rownames(pbmc.cite@assays$RNA))

pbmc.cite <- FindNeighbors(object = pbmc.cite, reduction = "pca", dims = 1:16)

pbmc.cite <- FindClusters(pbmc.cite, resolution = 0.55, Reduction.type = "pca")

pbmc.cite[["rna_7clusters"]] <- Idents(object = pbmc)

adt_pca_labels = pbmc.cite@active.ident

pbmc.cite <- RunTSNE(pbmc.cite, dims = 1:16, seed.use = seed)

DimPlot(object = pbmc.cite, reduction = 'tsne', label = TRUE, pt.size = 0.5)

pbmc.small <- SubsetData(pbmc.cite, max.cells.per.ident = 300)

adt.markers <- FindAllMarkers(pbmc.small, only.pos = TRUE)

DoHeatmap(pbmc.small, features = unique(adt.markers$gene),size=3)

DoHeatmap(pbmc.small, features = unique(adt.markers$gene),size=3,group.by = "rna_7clusters")

\# Cluser based on UMAP

seed = 42

pbmc.cite <- RunUMAP(pbmc.cite,features = rownames(pbmc.cite@assays$RNA))

pbmc.cite <- FindNeighbors(object = pbmc.cite, reduction = "umap", dims = 1:2)

pbmc.cite <- FindClusters(pbmc.cite, resolution = 0.02, Reduction.type = "umap")

DimPlot(object = pbmc.cite, reduction = 'umap', label = TRUE, pt.size = 0.5)

pbmc.small <- SubsetData(pbmc.cite, max.cells.per.ident = 300)

adt.markers <- FindAllMarkers(pbmc.small, only.pos = TRUE)

DoHeatmap(pbmc.small, features = unique(adt.markers$gene),size=3)

\# load data after filtering

data=benchmarkDataList$PBMC # in normal domain after filtering

colnames(data) <- paste0(colnames(data),"-1")

\# Intersection of data and the original data

data = data[, colnames(data) %in% colnames(pbmc@assays$RNA@data)]

\# Run RCA global panel to label cells

data_obj = dataConstruct(as.matrix(data))

data_obj = geneFilt(obj_in = data_obj)

data_obj = cellNormalize(data_obj)

data_obj = dataTransform(data_obj)

data_obj = featureConstruct(data_obj,method = "GlobalPanel",power = 4)

data_obj = cellClust(data_obj,deepSplit_wgcna=0)

RCAPlot(data_obj)

\# Run RCA Monaco panel to label cells

data_obj = dataConstruct(as.matrix(data))

data_obj = geneFilt(obj_in = data_obj)

data_obj = cellNormalize(data_obj)

data_obj = dataTransform(data_obj)

source('/mnt/volume1/project/code/featureConstructMonaco.R')

data_obj = featureConstructMonaco(data_obj,method = "MonacoPanel",power = 4)

data_obj = cellClust(data_obj,deepSplit_wgcna=3)

RCAPlot(data_obj)

#get the seurat label for filtered data

data_obj$seurat_labels = pbmc@active.ident[colnames(data)]

data_obj$adt_umap_labels = pbmc.cite@active.ident[colnames(data)]

data_obj$adt_pca_labels = adt_pca_labels[colnames(data)]

#Umap using RCA global panel clustering

seed = 42

seurat_obj = CreateSeuratObject(data_obj$fpkm_for_clust)

seurat_obj <- ScaleData(object = seurat_obj, display.progress = FALSE)

seurat_obj = RunUMAP(obj=seurat_obj,features = rownames(seurat_obj@assays$RNA@data))

seurat_obj@active.ident = as.factor(data_obj$group_labels_color$groupLabel)

names(seurat_obj@active.ident) = colnames(data)

colors = as.character(unique(data_obj$group_labels_color$dynamicColors))

labels = unique(data_obj$group_labels_color$groupLabel)

colors_sorted = colors[sort.int(labels,index.return = T)$ix]

DimPlot(object = seurat_obj, reduction = 'umap',group.by = "ident",cols = colors_sorted)

\# label using seurat labels

seurat_obj@active.ident = as.factor(data_obj$seurat_labels)

names(seurat_obj@active.ident) = colnames(data)

colors = unique(labels2colors(data_obj$seurat_labels))

DimPlot(object = seurat_obj, reduction = 'umap',group.by = "ident",cols = colors)

\# label using ADT labels

seurat_obj@active.ident = as.factor(data_obj$adt_pca_labels)

names(seurat_obj@active.ident) = colnames(data)

colors = unique(labels2colors(data_obj$adt_pca_labels))

DimPlot(object = seurat_obj, reduction = 'umap',group.by = "ident",cols = colors)

source('/mnt/volume1/project/plotHeatmap.R')

cellTypeDEPlot(data_obj$fpkm_for_clust,cellTree = data_obj$cellTree,clusterLabels = data_obj$group_labels_color$groupLabel,dynamicColors = data_obj$dynamicColors,filename = "heatmap_RCA_Monacopanel_deepsplit3")

\# Find the consensus groundtruth based on RCA and Seurat clustering

groundtruth = as.data.frame(data_obj$group_labels_color$groupLabel)

rownames(groundtruth) = colnames(data)

colnames(groundtruth) = "RCA"

groundtruth$RNA = data_obj$seurat_labels

groundtruth$ADT_Umap = data_obj$adt_umap_labels

groundtruth$ADT_pca = data_obj$adt_pca_labels

groundtruth$consensus = 0

groundtruth[groundtruth$RCA == 1 | groundtruth$RCA == 17 | groundtruth$RCA == 18,]$consensus = "C_Mono"

groundtruth[groundtruth$ADT_pca==0 & groundtruth$consensus == "C_Mono",]$consensus = "CD14 Mono"

groundtruth[groundtruth$ADT_pca==8 & groundtruth$consensus == "C_Mono",]$consensus = "CD16 Mono"

groundtruth[groundtruth$ADT_pca==10 & groundtruth$consensus == "C_Mono",]$consensus = "CD14 Mono"

groundtruth[groundtruth$ADT_pca==12 & groundtruth$consensus == "C_Mono",]$consensus = "CD16 Mono"

groundtruth[groundtruth$RCA == 2,]$consensus = "T1"

groundtruth[groundtruth$ADT_pca==1 & groundtruth$consensus == "T1",]$consensus = "CD4 T naive"

groundtruth[groundtruth$ADT_pca==3 & groundtruth$consensus == "T1",]$consensus = "CD4 T Memory"

groundtruth[groundtruth$ADT_pca==4 & groundtruth$consensus == "T1",]$consensus = "CD8 T Memory"

groundtruth[groundtruth$ADT_pca==6 & groundtruth$consensus== "T1",]$consensus = "TIGIT+ CD4 T"

groundtruth[groundtruth$ADT_pca==7 & groundtruth$consensus == "T1",]$consensus = "CD8 T Naive"

groundtruth[groundtruth$ADT_pca==9 & groundtruth$consensus == "T1",]$consensus = "TIGIT+ CD8 T"

groundtruth[groundtruth$ADT_pca==11 & groundtruth$consensus == "T1",]$consensus = "CD25+ CD4 T"

groundtruth[groundtruth$RCA == 3,]$consensus = "T2"

groundtruth[groundtruth$ADT_pca==1 & groundtruth$consensus == "T2",]$consensus = "CD4 T naive"

groundtruth[groundtruth$ADT_pca==3 & groundtruth$consensus == "T2",]$consensus = "CD4 T Memory"

groundtruth[groundtruth$ADT_pca==4 & groundtruth$consensus == "T2",]$consensus = "CD8 T Memory"

groundtruth[groundtruth$ADT_pca==6 & groundtruth$consensus== "T2",]$consensus = "TIGIT+ CD4 T"

groundtruth[groundtruth$ADT_pca==9 & groundtruth$consensus == "T2",]$consensus = "TIGIT+ CD8 T"

groundtruth[groundtruth$ADT_pca==11 & groundtruth$consensus == "T2",]$consensus = "CD25+ CD4 T"

groundtruth[groundtruth$RCA == 4,]$consensus = "NK"

groundtruth[groundtruth$RCA == 5,]$consensus = "B1"

groundtruth[groundtruth$RCA == 6,]$consensus = "T3"

groundtruth[groundtruth$ADT_pca==1 & groundtruth$consensus == "T3",]$consensus = "CD4 T naive"

groundtruth[groundtruth$ADT_pca==3 & groundtruth$consensus == "T3",]$consensus = "CD4 T Memory"

groundtruth[groundtruth$ADT_pca==4 & groundtruth$consensus == "T3",]$consensus = "CD8 T Memory"

groundtruth[groundtruth$ADT_pca==6 & groundtruth$consensus== "T3",]$consensus = "TIGIT+ CD4 T"

groundtruth[groundtruth$ADT_pca==7 & groundtruth$consensus== "T3",]$consensus = "CD8 T Naive"

groundtruth[groundtruth$ADT_pca==9 & groundtruth$consensus == "T3",]$consensus = "TIGIT+ CD8 T"

groundtruth[groundtruth$RCA == 7 | groundtruth$RCA == 19,]$consensus = "MAIT"

groundtruth[groundtruth$RCA == 8,]$consensus = "T4"

groundtruth[groundtruth$ADT_pca==1 & groundtruth$consensus == "T4",]$consensus = "CD4 T naive"

groundtruth[groundtruth$ADT_pca==3 & groundtruth$consensus == "T4",]$consensus = "CD4 T Memory"

groundtruth[groundtruth$ADT_pca==4 & groundtruth$consensus == "T4",]$consensus = "CD8 T Memory"

groundtruth[groundtruth$ADT_pca==6 & groundtruth$consensus== "T4",]$consensus = "TIGIT+ CD4 T"

groundtruth[groundtruth$ADT_pca==9 & groundtruth$consensus == "T4",]$consensus = "TIGIT+ CD8 T"

groundtruth[groundtruth$ADT_pca==11 & groundtruth$consensus == "T4",]$consensus = "CD25+ CD4 T"

groundtruth[groundtruth$RCA == 9 | groundtruth$RCA == 16,]$consensus = "T5"

groundtruth[groundtruth$ADT_pca==1 & groundtruth$consensus == "T5",]$consensus = "CD4 T naive"

groundtruth[groundtruth$ADT_pca==3 & groundtruth$consensus == "T5",]$consensus = "CD4 T Memory"

groundtruth[groundtruth$ADT_pca==4 & groundtruth$consensus == "T5",]$consensus = "CD8 T Memory"

groundtruth[groundtruth$ADT_pca==6 & groundtruth$consensus== "T5",]$consensus = "TIGIT+ CD4 T"

groundtruth[groundtruth$ADT_pca==7 & groundtruth$consensus== "T5",]$consensus = "CD8 T Naive"

groundtruth[groundtruth$ADT_pca==9 & groundtruth$consensus == "T5",]$consensus = "TIGIT+ CD8 T"

groundtruth[groundtruth$RCA == 10,]$consensus = "T6"

groundtruth[groundtruth$ADT_pca==1 & groundtruth$consensus == "T6",]$consensus = "CD4 T naive"

groundtruth[groundtruth$ADT_pca==3 & groundtruth$consensus == "T6",]$consensus = "CD4 T Memory"

groundtruth[groundtruth$ADT_pca==4 & groundtruth$consensus == "T6",]$consensus = "CD8 T Memory"

groundtruth[groundtruth$ADT_pca==6 & groundtruth$consensus== "T6",]$consensus = "TIGIT+ CD4 T"

groundtruth[groundtruth$ADT_pca==7 & groundtruth$consensus== "T6",]$consensus = "CD8 T Naive"

groundtruth[groundtruth$ADT_pca==9 & groundtruth$consensus == "T6",]$consensus = "TIGIT+ CD8 T"

groundtruth[groundtruth$RCA == 11,]$consensus = "NC_Mono"

groundtruth[groundtruth$ADT_pca==0 & groundtruth$consensus == "NC_Mono",]$consensus = "CD14 Mono"

groundtruth[groundtruth$ADT_pca==8 & groundtruth$consensus == "NC_Mono",]$consensus = "CD16 Mono"

groundtruth[groundtruth$ADT_pca==10 & groundtruth$consensus == "NC_Mono",]$consensus = "CD14 Mono"

groundtruth[groundtruth$RCA == 12,]$consensus = "mDC"

groundtruth[groundtruth$RCA == 13,]$consensus = "B2"

groundtruth[groundtruth$RCA == 14,]$consensus = "pDC"

groundtruth[groundtruth$RCA == 15,]$consensus = "Plasma"

groundtruth[groundtruth$RCA == 20,]$consensus = "Progenitor"

dim(groundtruth[groundtruth$consensus==0,])

sort(table(groundtruth$consensus))

groundtruth[groundtruth$consensus == "T1" | groundtruth$consensus == "T2" |

groundtruth$consensus == "T3" | groundtruth$consensus == "T4"|

groundtruth$consensus == "T5" | groundtruth$consensus == "T6",]$consensus = "T"

groundtruth[groundtruth$consensus == "NC_Mono",]$consensus = "CD16 Mono"

groundtruthDataList = list()

groundtruthDataList[["pbmc"]] = data[,rownames(groundtruth[groundtruth$consensus!=0,])]

groundtruthGroupList = list()

groundtruthGroupList[["pbmc"]] = groundtruth[groundtruth$consensus!=0,]$consensus

save(groundtruthDataList,groundtruthGroupList,file="groundtruth_PBMC_Monaco.RData")

\# Find the consensus groundtruth based on RCA and Seurat clustering

groundtruth = as.data.frame(data_obj$group_labels_color$groupLabel)

rownames(groundtruth) = colnames(data)

colnames(groundtruth) = "RCA"

groundtruth$RNA = data_obj$seurat_labels

groundtruth$ADT_Umap = data_obj$adt_umap_labels

groundtruth$ADT_pca = data_obj$adt_pca_labels

groundtruth$consensus = 0

groundtruth[groundtruth$RCA == 1 | groundtruth$RCA == 17 | groundtruth$RCA == 18,]$consensus = "C_Mono"

groundtruth[groundtruth$ADT_pca==0 & groundtruth$consensus == "C_Mono",]$consensus = ""

groundtruth[groundtruth$ADT_pca==2 & groundtruth$RCA == 1,]$consensus = "CD4 T Memory"

groundtruth[groundtruth$ADT_pca==4 & groundtruth$RCA == 1,]$consensus = "CD8 T Memory"

groundtruth[groundtruth$ADT_pca==6 & groundtruth$RCA == 1,]$consensus = "CD8 T naive"

groundtruth[groundtruth$ADT_pca==7 & groundtruth$RCA == 1,]$consensus = "TIGIT+ CD8 T"

groundtruth[groundtruth$ADT_pca==9 & groundtruth$RCA == 1,]$consensus = "TIGIT+ CD4 T"

groundtruth[groundtruth$RCA == 1 | groundtruth$RCA == 17 | groundtruth$RCA == 18,]$consensus = "C_Mono"

groundtruth[groundtruth$ADT_pca==0 & groundtruth$consensus == "C_Mono",]$consensus = "CD14 Mono"

groundtruth[groundtruth$ADT_pca==1 & groundtruth$RCA == 2,]$consensus = "CD16 Mono"

groundtruth[groundtruth$ADT_pca==10 & groundtruth$RCA == 2,]$consensus = "CD14 Mono"

groundtruth[groundtruth$ADT_pca==11 & groundtruth$RCA == 2,]$consensus = "CD16 Mono"

groundtruth[groundtruth$RCA == 3,]$consensus = "NK"

groundtruth[groundtruth$RCA == 4,]$consensus = "B"

groundtruth[groundtruth$RCA == 5,]$consensus = "DC"

groundtruth[groundtruth$RCA == 6,]$consensus = "Plasma"

groundtruth[groundtruth$RCA == 7,]$consensus = "precursors"

dim(groundtruth[groundtruth$consensus==0,])

table(groundtruth$consensus)

groundtruthDataList = list()

groundtruthDataList[["pbmc"]] = data[,rownames(groundtruth[groundtruth$consensus!=0,])]

groundtruthGroupList = list()

groundtruthGroupList[["pbmc"]] = groundtruth[groundtruth$consensus!=0,]$consensus

save(groundtruthDataList,groundtruthGroupList,file="groundtruth_PBMC.RData")

###############################VDJ###############################################

\# load the original data

data_original = readMM("/mnt/volume1/project/vdj_8k/matrix.mtx.gz")

data_genes = read.csv("/mnt/volume1/project/vdj_8k/features.tsv.gz",header=F,sep="\t",stringsAsFactors = F)

data_cells = read.csv("/mnt/volume1/project/vdj_8k/barcodes.tsv.gz",header=F,sep="\t",stringsAsFactors = F)

data_original@Dimnames[[1]] = data_genes$V2

data_original@Dimnames[[2]] = data_cells$V1

vdj.rna.collapsed <- CollapseSpeciesExpressionMatrix(data_original[1:33538,])

vdj <- CreateSeuratObject(vdj.rna.collapsed,min.cells = 100, min.features = 500, project = "vdj")

mito.features <- grep(pattern = "^MT-", x = rownames(x = vdj), value = TRUE)

percent.mito <- Matrix::colSums(x = GetAssayData(object = vdj, slot = 'counts')[mito.features, ]) / Matrix::colSums(x = GetAssayData(object = vdj, slot = 'counts'))

vdj[['percent.mito']] <- percent.mito

VlnPlot(object = vdj, features = c("nFeature_RNA", "nCount_RNA", "percent.mito"), ncol = 3)

vdj <- subset(x = vdj, subset = nFeature_RNA > 500 & nFeature_RNA < 4000 & percent.mito < 0.3)

vdj <- NormalizeData(vdj)

vdj <- FindVariableFeatures(vdj,selection.method = "vst",nfeatures = 1000)

vdj <- ScaleData(vdj, display.progress = FALSE)

vdj <- RunPCA(vdj,features = VariableFeatures(object = vdj))

ElbowPlot(vdj)

vdj <- FindNeighbors(object = vdj, reduction = "pca", dims = 1:15)

vdj <- FindClusters(vdj, resolution = 0.1)

vdj <- RunTSNE(vdj, dims = 1:15)

vdj.markers <- FindAllMarkers(object = vdj, only.pos = TRUE, min.pct = 0.25, logfc.threshold = 0.25)

vdj.markers %>% group_by(cluster) %>% top_n(n = 5, wt = avg_logFC) %>% print(n=26)

filter(vdj.markers, cluster == "0") %>% top_n(n=20,wt=avg_logFC)

new.cluster.ids <- c("CD14 Mono", "CD4 T", "B","CD8 T", "NK", "Unknown", "FCGR3A Mono","DC","CD8 T","Mk")

names(x = new.cluster.ids) <- levels(x = vdj)

vdj <- RenameIdents(object = vdj, new.cluster.ids)

DimPlot(object = vdj, reduction = 'tsne', label = TRUE, pt.size = 0.5)

table(vdj@active.ident)

vdj[["rna_10clusters"]] <- Idents(object = vdj)

\# load the ADT UMI data

vdj.adt <- as.matrix(data_original[33539:33555,])

vdj.adt = vdj.adt[,names(vdj@active.ident)]

vdj.cite <- CreateSeuratObject(counts = vdj.adt)

vdj.cite = NormalizeData(object = vdj.cite, method = "CLR")

vdj.cite <- ScaleData(vdj.cite, features = NULL, display.progress = FALSE)

\###### cluster based on ADT data

\# Cluster based on PCA

vdj.cite <- RunPCA(vdj.cite, features = rownames(vdj.cite@assays$RNA))

vdj.cite <- FindNeighbors(object = vdj.cite, reduction = "pca", dims = 1:16)

vdj.cite <- FindClusters(vdj.cite, resolution = 0.2, Reduction.type = "pca")

vdj.cite[["rna_10clusters"]] <- Idents(object = vdj)

adt_pca_labels = vdj.cite@active.ident

vdj.cite <- RunTSNE(vdj.cite, dims = 1:16)

DimPlot(object = vdj.cite, reduction = 'tsne', label = TRUE, pt.size = 0.5)

vdj.small <- SubsetData(vdj.cite, max.cells.per.ident = 300)

adt.markers <- FindAllMarkers(vdj.small, only.pos = TRUE)

DoHeatmap(vdj.small, features = unique(adt.markers$gene),size=3)

DoHeatmap(vdj.small, features = unique(adt.markers$gene),size=3,group.by = "rna_10clusters")

\# Cluser based on UMAP

vdj.cite <- RunUMAP(vdj.cite,features = rownames(vdj.cite@assays$RNA))

vdj.cite <- FindNeighbors(object = vdj.cite, reduction = "umap", dims = 1:2)

vdj.cite <- FindClusters(vdj.cite, resolution = 0.01, Reduction.type = "umap")

DimPlot(object = vdj.cite, reduction = 'umap', label = TRUE, pt.size = 0.5)

vdj.small <- SubsetData(vdj.cite, max.cells.per.ident = 300)

adt.markers <- FindAllMarkers(vdj.small, only.pos = TRUE)

DoHeatmap(vdj.small, features = unique(adt.markers$gene),size=3)

\# load data after filtering

data=benchmarkDataList$VDJ # in normal domain after filtering

colnames(data) <- paste0(colnames(data),"-1")

\# Intersection of data and the original data

data = data[, colnames(data) %in% colnames(vdj@assays$RNA@data)]

\# Run RCA global panel to label cells

data_obj = dataConstruct(as.matrix(data))

data_obj = geneFilt(obj_in = data_obj)

data_obj = cellNormalize(data_obj)

data_obj = dataTransform(data_obj)

#data_obj = featureConstruct(data_obj,method = "GlobalPanel",power = 4)

source('/mnt/volume1/project/code/featureConstructMonaco.R')

data_obj = featureConstructMonaco(data_obj,method = "MonacoPanel",power = 4)

data_obj = cellClust(data_obj,deepSplit_wgcna=2)

RCAPlot(data_obj)

#get the seurat label for filtered data

data_obj$seurat_labels = vdj@active.ident[colnames(data)]

data_obj$adt_umap_labels = vdj.cite@active.ident[colnames(data)]

data_obj$adt_pca_labels = adt_pca_labels[colnames(data)]

#Umap using RCA global panel clustering

seed = 42

seurat_obj = CreateSeuratObject(data_obj$fpkm_for_clust)

seurat_obj <- ScaleData(object = seurat_obj, display.progress = FALSE)

seurat_obj = RunUMAP(obj=seurat_obj,features = rownames(seurat_obj@assays$RNA@data))

seurat_obj@active.ident = as.factor(data_obj$group_labels_color$groupLabel)

names(seurat_obj@active.ident) = colnames(data)

colors = as.character(unique(data_obj$group_labels_color$dynamicColors))

labels = unique(data_obj$group_labels_color$groupLabel)

colors_sorted = colors[sort.int(labels,index.return = T)$ix]

DimPlot(object = seurat_obj, reduction = 'umap',group.by = "ident",cols = colors_sorted)

\# label using seurat labels

seurat_obj@active.ident = as.factor(data_obj$seurat_labels)

names(seurat_obj@active.ident) = colnames(data)

colors = unique(labels2colors(data_obj$seurat_labels))

DimPlot(object = seurat_obj, reduction = 'umap',group.by = "ident",cols = colors)

\# label using ADT labels

seurat_obj@active.ident = as.factor(data_obj$adt_umap_labels)

names(seurat_obj@active.ident) = colnames(data)

colors = unique(labels2colors(data_obj$adt_umap_labels))

DimPlot(object = seurat_obj, reduction = 'umap',group.by = "ident",cols = colors)

source('/mnt/volume1/project/plotHeatmap.R')

cellTypeDEPlot(data_obj$fpkm_for_clust,cellTree = data_obj$cellTree,clusterLabels = data_obj$group_labels_color$groupLabel,dynamicColors = data_obj$dynamicColors,filename = "heatmap_RCA_Monacopanel_deepsplit2")

\# Find the consensus groundtruth based on RCA and Seurat clustering

groundtruth = as.data.frame(data_obj$group_labels_color$groupLabel)

rownames(groundtruth) = colnames(data)

colnames(groundtruth) = "RCA"

groundtruth$RNA = data_obj$seurat_labels

groundtruth$ADT_Umap = data_obj$adt_umap_labels

groundtruth$ADT_pca = data_obj$adt_pca_labels

groundtruth$consensus = 0

groundtruth[groundtruth$RCA == 1,]$consensus = "Mono"

groundtruth[groundtruth$RCA == 2,]$consensus = "T"

groundtruth[groundtruth$ADT_pca==1 & groundtruth$RCA == 2,]$consensus = "CD4 T"

groundtruth[groundtruth$ADT_pca==3 & groundtruth$RCA == 2,]$consensus = "CD8 T"

groundtruth[groundtruth$RCA == 3,]$consensus = "B"

groundtruth[groundtruth$RCA == 4,]$consensus = "MAIT"

groundtruth[groundtruth$ADT_pca==5 & groundtruth$RCA == 5,]$consensus = "NK"

groundtruth[groundtruth$ADT_pca!=5 & groundtruth$RCA == 5,]$consensus = "CD8 TE"

groundtruth[groundtruth$RCA == 6,]$consensus = "mDC"

groundtruth[groundtruth$RCA == 7,]$consensus = "pDC"

groundtruth[groundtruth$RCA == 8,]$consensus = "Mono"

groundtruth[groundtruth$RCA == 9,]$consensus = "plasma"

groundtruth[groundtruth$RCA == 10,]$consensus = "basophil"

groundtruth[groundtruth$RCA == 11,]$consensus = "B"

groundtruth[groundtruth$RCA == 12,]$consensus = "neutrophil"

groundtruth[groundtruth$RCA == 15,]$consensus = "progenitor"

dim(groundtruth[groundtruth$consensus==0,])

table(groundtruth$consensus)

groundtruthDataList = list()

groundtruthDataList[["vdj"]] = data[,rownames(groundtruth[groundtruth$consensus!=0,])]

groundtruthGroupList = list()

groundtruthGroupList[["vdj"]] = groundtruth[groundtruth$consensus!=0,]$consensus

save(groundtruthDataList,groundtruthGroupList,file="groundtruth_vdj.RData")

#### **Supplementary Note 4** – Preprocessing and scConsensus Execution for FACS-sorted PBMC data

##### FACS sorted PBMC data from Zheng et al. (2017)

##### # 26k FACS_sorted PBMC object preparation

pipestance_path1 = "~/cd14_monocytes/outs/filtered_gene_bc_matrices_mex/hg19/"

pipestance_path2 = "~/CD19+ B Cells/outs/filtered_gene_bc_matrices_mex/hg19/"

pipestance_path3 = "~/CD34+ Cells/outs/filtered_gene_bc_matrices_mex/hg19/"

pipestance_path4 = "~/CD4+ Helper T Cells/outs/filtered_gene_bc_matrices_mex/hg19/"

pipestance_path5 = "~/CD4+CD25+ Regulatory T Cells/outs/filtered_gene_bc_matrices_mex/hg19/"

pipestance_path6 = "~/CD4+CD45RA+CD25- Naive T cells/outs/filtered_gene_bc_matrices_mex/hg19/"

pipestance_path7 = "~/CD4+CD45RO+ Memory T Cells/outs/filtered_gene_bc_matrices_mex/hg19/"

pipestance_path8 = "~/CD56+ Natural Killer Cells/outs/filtered_gene_bc_matrices_mex/hg19/"

pipestance_path9 = "~/CD8+ Cytotoxic T cells/outs/filtered_gene_bc_matrices_mex/hg19/"

pipestance_path10 = "~/CD8+CD45RA+ Naive Cytotoxic T Cells/outs/filtered_gene_bc_matrices_mex/hg19/"

set.seed(1)

pbmc1.data <- Read10X(data.dir = pipestance_path1)

pbmc1.data <- pbmc1.data[,sample(1:ncol (pbmc1.data), 2600)]

pbmc1 <- CreateSeuratObject(counts = pbmc1.data, project = "CD14_Monocytes")

pbmc2.data <- Read10X(data.dir = pipestance_path2)

pbmc2.data <- pbmc2.data[,sample(1:ncol (pbmc2.data), 2600)]

pbmc2 <- CreateSeuratObject(counts = pbmc2.data, project = "B_Cells")

pbmc3.data <- Read10X(data.dir = pipestance_path3)

pbmc3.data <- pbmc3.data[,sample(1:ncol (pbmc3.data), 2600)]

pbmc3 <- CreateSeuratObject(counts = pbmc3.data, project = "CD34_Cells")

pbmc4.data <- Read10X(data.dir = pipestance_path4)

pbmc4.data <- pbmc4.data[,sample(1:ncol (pbmc4.data), 2600)]

pbmc4 <- CreateSeuratObject(counts = pbmc4.data, project = "CD4_T-Helper")

pbmc5.data <- Read10X(data.dir = pipestance_path5)

pbmc5.data <- pbmc5.data[,sample(1:ncol (pbmc5.data), 2600)]

pbmc5 <- CreateSeuratObject(counts = pbmc5.data, project = "T_Reg")

pbmc6.data <- Read10X(data.dir = pipestance_path6)

pbmc6.data <- pbmc6.data[,sample(1:ncol (pbmc6.data), 2600)]

pbmc6 <- CreateSeuratObject(counts = pbmc6.data, project = "CD4_T-Naive")

pbmc7.data <- Read10X(data.dir = pipestance_path7)

pbmc7.data <- pbmc7.data[,sample(1:ncol (pbmc7.data), 2600)]

pbmc7 <- CreateSeuratObject(counts = pbmc7.data, project = "CD4_T-Memory")

pbmc8.data <- Read10X(data.dir = pipestance_path8)

pbmc8.data <- pbmc8.data[,sample(1:ncol (pbmc8.data), 2600)]

pbmc8 <- CreateSeuratObject(counts = pbmc8.data, project = "NK_Cells")

pbmc9.data <- Read10X(data.dir = pipestance_path9)

pbmc9.data <- pbmc9.data[,sample(1:ncol (pbmc9.data), 2600)]

pbmc9 <- CreateSeuratObject(counts = pbmc9.data, project = "CD8_T-Cytotoxic")

pbmc10.data <- Read10X(data.dir = pipestance_path10)

pbmc10.data <- pbmc10.data[,sample(1:ncol (pbmc10.data), 2600)]

pbmc10 <- CreateSeuratObject(counts = pbmc10.data, project = "CD8_T-Naive")

pbmc_26k_S.v3 <- merge(pbmc1, y = c(pbmc2, pbmc3, pbmc4, pbmc5, pbmc6, pbmc7, pbmc8, pbmc9, pbmc10), add.cell.ids = c("1", "2", "3", "4", "5", "6", "7", "8", "9", "10"), project = "PBMC26K")

\# QC filtering

pbmc_26k_S.v3[["percent.mt"]] <- PercentageFeatureSet(pbmc_26k_S.v3, pattern = "^MT-")

VlnPlot(pbmc_26k_S.v3, features = c("nFeature_RNA", "nCount_RNA", "percent.mt"), ncol = 3, pt.size = 0)

plot1 <- FeatureScatter(pbmc_26k_S.v3, feature1 = "nCount_RNA", feature2 = "percent.mt")

plot2 <- FeatureScatter(pbmc_26k_S.v3, feature1 = "nCount_RNA", feature2 = "nFeature_RNA")

CombinePlots(plots = list(plot1, plot2))

pbmc_26k_S.v3_Filtered <- subset(pbmc_26k_S.v3, subset = nFeature_RNA > 300 & nFeature_RNA < 2000 & percent.mt < 8)

\# Unsupervised clustering (seurat pipeline)

pbmc_26k_S.v3_Filtered <- NormalizeData(pbmc_26k_S.v3_Filtered, normalization.method = "LogNormalize", scale.factor = 10000)

pbmc_26k_S.v3_Filtered <- FindVariableFeatures(pbmc_26k_S.v3_Filtered, selection.method = "vst", nfeatures = 2000)

top10 <- head(VariableFeatures(pbmc_26k_S.v3_Filtered), 10)

plot1 <- VariableFeaturePlot(pbmc_26k_S.v3_Filtered)

plot2 <- LabelPoints(plot = plot1, points = top10, repel = TRUE)

CombinePlots(plots = list(plot1, plot2))

pbmc_26k_S.v3_Filtered <- ScaleData(pbmc_26k_S.v3_Filtered, vars.to.regress = "percent.mt")

pbmc_26k_S.v3_Filtered <- RunPCA(pbmc_26k_S.v3_Filtered, features = VariableFeatures(object = pbmc_26k_S.v3_Filtered))

DimHeatmap(pbmc_26k_S.v3_Filtered, dims = 1:15, cells = 500, balanced = TRUE)

pbmc_26k_S.v3_Filtered <- JackStraw(pbmc_26k_S.v3_Filtered, num.replicate = 100)

pbmc_26k_S.v3_Filtered <- ScoreJackStraw(pbmc_26k_S.v3_Filtered, dims = 1:20)

JackStrawPlot(pbmc_26k_S.v3_Filtered, dims = 1:20)

ElbowPlot(pbmc_26k_S.v3_Filtered)

pbmc_26k_S.v3_Filtered <- FindNeighbors(pbmc_26k_S.v3_Filtered, reduction = "pca", dims = 1:20)

pbmc_26k_S.v3_Filtered <- FindClusters(pbmc_26k_S.v3_Filtered, resolution = 0.2, algorithm = 1)

pbmc_26k_S.v3_Filtered <- RunUMAP(pbmc_26k_S.v3_Filtered, dims = 1:20)

DimPlot(pbmc_26k_S.v3_Filtered, reduction = "umap", group.by = "nFeature_RNA")

DimPlot(pbmc_26k_S.v3_Filtered, reduction = "umap", group.by = "orig.ident")

FeaturePlot(pbmc_26k_S.v3_Filtered, features = c("IL7R", "CD8A", "GNLY", "CD79A","CD14","FCER1A", "FCGR3A", "SERPINF1", "PPBP"))

pbmc.markers <- FindAllMarkers(pbmc_26k_S.v3_Filtered, only.pos = TRUE, min.pct = 0.25, logfc.threshold = 0.25)

\# Save the seurat object:

saveRDS(pbmc_26k_S.v3_Filtered, file = "pbmc_26k_S.v3_Filtered.rds")
